# Enhancer-AAVs allow genetic access to oligodendrocytes and diverse populations of astrocytes across species

**DOI:** 10.1101/2023.09.20.558718

**Authors:** John K. Mich, Smrithi Sunil, Nelson Johansen, Refugio A. Martinez, Jiatai Liu, Bryan B. Gore, Joseph T. Mahoney, Mckaila Leytze, Yoav Ben-Simon, Darren Bertagnolli, Ravi Bhowmik, Yemeserach Bishaw, Krissy Brouner, Jazmin Campos, Ryan Canfield, Tamara Casper, Nicholas P. Donadio, Nadezhda I. Dotson, Tom Egdorf, Amanda Gary, Shane Gibson, Jeff Goldy, Erin L. Groce, Kenta M. Hagihara, Daniel Hirschstein, Han Hou, Will D. Laird, Elizabeth Liang, Luke V. Loftus, Nicholas Lusk, Jocelin Malone, Naomi X. Martin, Deja Monet, Josh S. Nagra, Dakota Newman, Nhan-Kiet Ngo, Paul A. Olsen, Victoria Omstead, Ximena Opitz-Araya, Aaron Oster, Christina A. Pom, Lydia Potekhina, Melissa Reding, Christine Rimorin, Augustin Ruiz, Adriana E. Sedeño-Cortés, Nadiya V. Shapovalova, Michael Taormina, Naz Taskin, Michael Tieu, Nasmil J. Valera Cuevas, Sharon W. Way, Natalie Weed, Vonn Wright, Zizhen Yao, Thomas Zhou, Delissa A. McMillen, Michael Kunst, Medea McGraw, Bargavi Thyagarajan, Jack Waters, Trygve E. Bakken, Nick Dee, Shenqin Yao, Kimberly A. Smith, Karel Svoboda, Kaspar Podgorski, Yoshiko Kojima, Greg D. Horwitz, Hongkui Zeng, Tanya L. Daigle, Ed S. Lein, Bosiljka Tasic, Jonathan T. Ting, Boaz P. Levi

**Affiliations:** Allen Institute for Brain Science, Seattle WA, USA; Allen Institute for Neural Dynamics, Seattle WA, USA; Department of Otolaryngology Head and Neck Surgery, University of Washington, Seattle, WA, USA; Department of Neurobiology and Biophysics, University of Washington, Seattle, WA, USA; Division of Medical Genetics, University of Washington, Seattle, WA, USA; Washington National Primate Research Center, University of Washington, Seattle, WA, USA; Department of Neurological Surgery, Univ. of Washington, Seattle WA; Department of Laboratory Medicine and Pathology, Univ. of Washington, Seattle WA

## Abstract

Proper brain function requires the assembly and function of diverse populations of neurons and glia. Single cell gene expression studies have mostly focused on characterization of neuronal cell diversity; however, recent studies have also revealed substantial diversity of glial cells, particularly astrocytes. To better understand glial cell types and their roles in neurobiology, we built a new suite of adeno-associated viral (AAV)-based genetic tools to enable genetic access to astrocytes and oligodendrocytes. These oligodendrocyte and astrocyte enhancer-AAVs are highly specific (usually > 95% cell type specificity) with variable expression levels, and the astrocyte enhancer-AAVs show multiple distinct expression patterns reflecting the spatial distribution of astrocyte cell types. To provide the best glial-specific functional tools, several enhancer-AAVs were: optimized for higher expression levels, shown to be functional and specific in rat and macaque, shown to maintain specific activity across transgenes and in epilepsy where traditional promoters changed activity, and used to drive functional transgenes in astrocytes including Cre recombinase and acetylcholine-responsive sensor iAChSnFR. The astrocyte-specific iAChSnFR revealed a clear reward-dependent acetylcholine response in astrocytes of the nucleus accumbens during reinforcement learning. Together, this collection of glial enhancer-AAVs will enable characterization of astrocyte and oligodendrocyte populations and their functional roles.

## Introduction

Glial cell types play critical roles in CNS development, function, and homeostasis^1–3^. Astrocytes sculpt neuronal circuits^4–9^, regulate and support synaptic transmission^9–11^, and drive beneficial and/or pathologic injury response^12–15^. Oligodendrocytes form myelin sheaths^16^, strengthen circuits adaptively^17–19^, and contribute to pathologic disease progression^20,21^. Transcriptomic characterization of glial cells has revealed an array of astrocyte and oligodendrocyte cell types, often with pronounced regional signatures^22–24^. Furthermore, species-specific features have been described^25^, although the functional significance of these differences is unknown. Glial cell types have also been shown to play critical roles in CNS diseases ranging from epilepsy^26^ to neurodegenerative diseases^27^ to cancer^28,29^. To understand how glia differ between cell types, regions, species, and disease states, a set of tools is needed to grant targeted genetic access to these specific populations across species.

Adeno-associated virus (AAV) vectors are exceptionally useful tools for somatic transgenesis across mammalian species including human^30–35^. Short enhancer or promoter regulatory elements work effectively in AAV expression cassettes to drive cell-type selective gene expression in the brain or other organs. Recent work has shown that selective AAVs can be rationally designed by using enhancers identified from epigenetic datasets that are selectively active and fit in an AAV vector^36–46^. Enhancer-AAVs were recently developed to target different populations of excitatory and inhibitory neurons in the brain, and some enhancers have shown successful targeting of glial cell populations as well^41,47–49^.

However, the field largely relies on glial promoters that have some undesirable characteristics, most notably loss of specificity or change in strength in different contexts as seen for astrocytic *GFAP* promoter fragments^50–52^. Furthermore, single cell genomics studies have revealed region-specific astrocyte cell types for which no current tools are available^22–24^.

Here we present characterization of a collection of enhancer-AAVs that selectively target astrocytes and oligodendrocytes. Twenty-five astrocyte and 21 oligodendrocyte enhancer AAVs were identified from mouse and human neocortical epigenetic data that produced reporter expression that was highly specific for the intended populations, often labeling more than half of the intended cells in the area, and with a wide range of expression strengths. Multiple astrocyte-targeting vectors exhibited distinct CNS region-specific expression patterns, whereas oligodendrocyte-selective vectors generally drove expression throughout the entire CNS. Several enhancer-AAVs maintained selective expression for astrocytes or oligodendrocytes across rat and macaque, in one model of epileptic disease, and even for difficult transgenes. Lastly, several astrocyte tools were adapted to drive expression of functional transgenes like Cre or the detection of neurotransmitters to reveal the role of astrocytes in neurobiology. We used astrocyte-selective AAV expressing iAChSnFR^53^ to measure the dynamics of acetylcholine in astrocytes of the nucleus accumbens during reinforcement learning. This collection of tools opens up new opportunities for selective labeling and functional interrogation of glial cell types across species and in some disease states, and could have translational applications via AAV-based therapeutics^54–56^.

## Results

### Generation of astrocyte- and oligodendrocyte-specific enhancer-AAVs

We identified putative enhancers specific for astrocytes and oligodendrocytes from single cell/single nucleus assay for transposase-accessible chromatin (sc/snATAC-seq^39,40^) and single nucleus methyl-cytosine sequencing (snmC-seq) studies from neocortex^57–60^. Thousands of astrocyte- and oligodendrocyte-selective scATAC-seq peaks were identified previously in both human middle temporal gyrus (MTG) and mouse primary visual cortex (VISp), averaging approximately 300-600 bp in size (**Figure 1A**). Additional ATAC-seq datasets confirmed these peaks^61,62^. Generally, astrocyte and oligodendrocyte candidate enhancers were accessible in non-neuronal cells but not in neuronal cells across the human forebrain^39,61^ (**Figure 1—figure supplement 1A-F**), and in the corresponding astrocyte or oligodendrocyte subclasses across the mouse forebrain without strong cell type preferences^62^ (**Figure 1—figure supplement 1G-L**).

**Figure 1:**
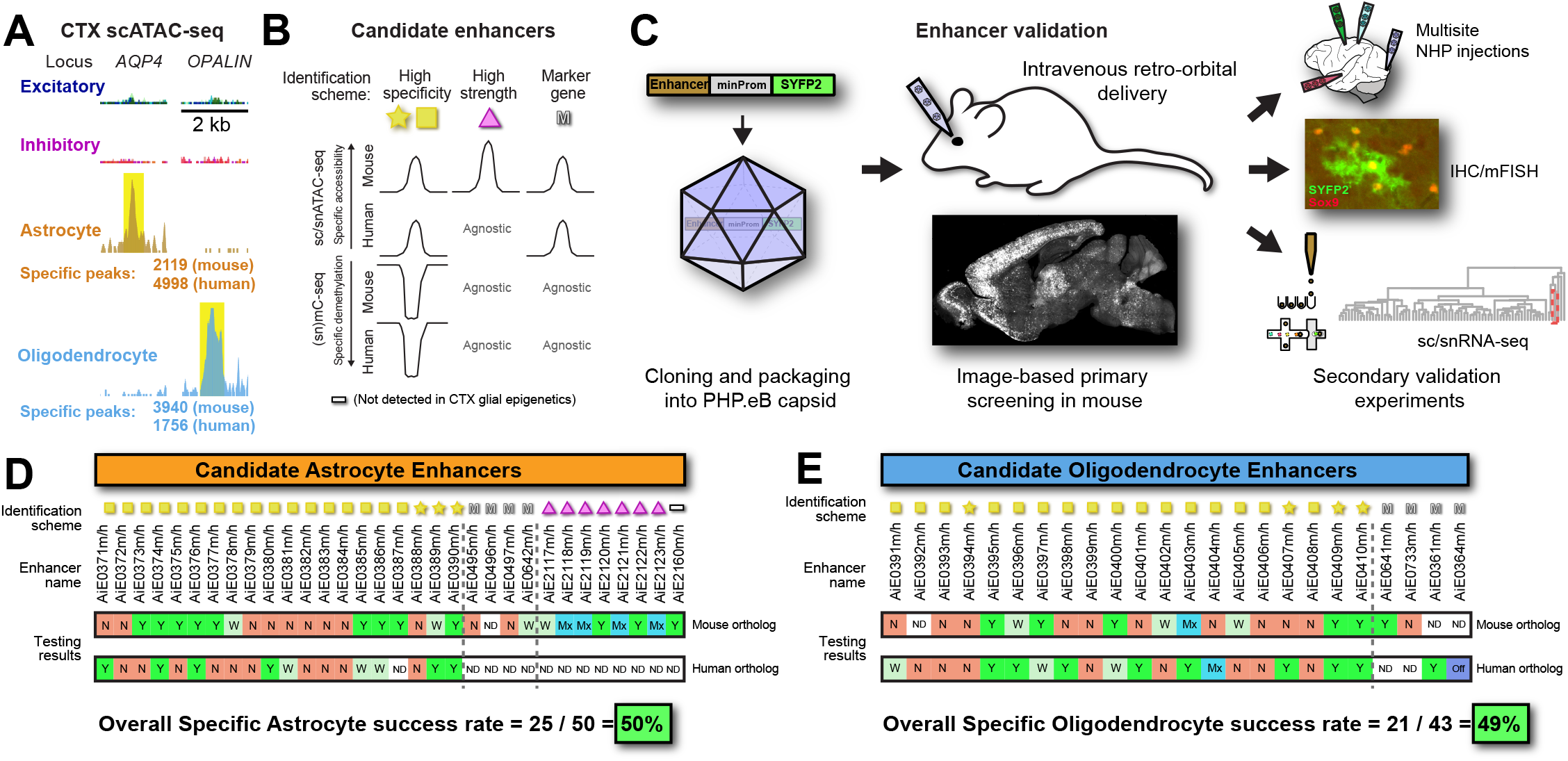
Astrocyte and oligodendrocyte enhancer discovery from single cell epigenetics. **(A)** Example astrocyte- and oligodendrocyte-specific peaks near the loci of astrocyte-specific gene *AQP4* and oligodendrocyte-specific gene *OPALIN*, identified in human MTG snATAC-seq data^39^. **(B)** Differing approaches to identify candidate enhancers. Specific accessibility peaks are depicted as peaks, and specifically demethylated regions are depicted as troughs. Schemes not utilizing a particular data modality are shown as “Agnostic”. Marker gene selection criteria can use accessibility from either mouse or human. Icons represent identification schemes; gold star candidate enhancers undergo more stringent criteria than those with gold squares (see Methods for details). **(C)** Workflow for enhancer cloning, packaging, screening, and validation. Enhancers are cloned into a pAAV plasmid upstream of a minimal human beta-globin promoter and SYFP2 reporter, and plasmids are packaged into PHP.eB AAVs. Enhancer-AAVs are injected intravenously into retro-orbital sinus, and expression is assessed by imaging. Promising enhancer-AAVs then go on to secondary validation experiments consisting of cross-species validation, molecular characterization by IHC and/or multiplexed FISH, and flow cytometry for single cell RNA-seq. (**D-E**) Candidate identification schemes and summarized screening results for astrocyte-specific (**D**) and oligodendrocyte-specific enhancers (**E**). Overall 50% (25/50) of candidate astrocyte enhancers, and 49% (21/43) of candidate oligodendrocyte enhancers, show the intended specificity after intravenous delivery of PHP.eB AAVs. Testing result bar: Y = yes, enhancer-AAV gives strong or moderate on-target expression pattern; N = no, enhancer-AAV fails to express; W = weak on-target expression pattern; Mx = mixed expression pattern consisting of on-target cells plus unwanted neuronal populations; Off = off-target expression pattern; ND = no data. Note both enhancers giving strong/moderate (“Y”) and weak (“W”) specific expression are grouped here for overall success rate analysis.

We used three strategies to identify putative enhancers for testing: “high specificity”, “high strength”, and “marker gene” (**Figure 1B-E**). The “high specificity” nomination criteria (gold square or star icons) required enhancers and their orthologs to show accessibility specifically for both mouse and human astrocytes or oligodendrocytes but not other cell types^39,40^. In addition, we required that these putative enhancers not be detected in demethylated genomic regions in both mouse and human neuron populations^58^. Some “high specificity” enhancers also showed specific demethylation in bulk human and mouse glial cells^57^ (marked by gold star icons). “High strength” putative enhancers were selected on the basis of strong astrocyte-specific peaks using only mouse scATAC-seq data^40^, with strength measured by accessibility read count within peaks. Finally, “marker gene” putative enhancers showed specific and strong accessibility near known astrocyte- and oligodendrocyte-specific marker genes.

We tested putative enhancer function in AAV vectors upstream of a minimal promoter driving the reporter SYFP2 and evaluated expression throughout the mouse brain after systemic administration of PHP.eB-serotyped AAVs. Enhancer-AAVs that showed anticipated reporter expression, were further evaluated for specificity, completeness of expression, and cross-species activity (**Figure 1C** and **Table S1**). All three strategies were effective, with approximately half of the candidates for both cell types yielding astrocyte or oligodendrocyte-specific expression patterns during primary screening (**Figure 1D, E**), which were confirmed on-target by immunohistochemistry (IHC) and/or scRNA-seq (see figures below). Some of these vectors were also characterized in parallel large-scale efforts^41^, all raw data from which are available online at https://portal.brain-map.org/genetic-tools/genetic-tools-atlas, and the full set of experimental mice described below are summarized in **Table S2**.

### A collection of astrocyte-specific enhancer-AAVs

We screened 50 candidate astrocyte-specific enhancer-AAVs, and 25 (50%) of them labeled astrocytes specifically with SYFP2 expression (**Figure 2—figure supplement 1, 2**). We defined specificity as the number of SYFP2+Sox9+ / total SYFP2+ cells x 100%. Completeness is defined as the number of SYFP2+Sox9+ / total Sox9+ cells x 100%.

Astrocyte-specific enhancer-AAVs showed a range of expression strengths and patterns (**Figure 2A-O**) and vectors were categorized based on their labeling as: “Most of the CNS”, “Regional”, “Sparse”, “Weak” and “Mixed specificities”. “Most of the CNS” astrocyte enhancer-AAVs, including AiE0380h and the human *GFAP* promoter (GfaABC1D^63^), labeled cells with astrocyte morphology in both brain and spinal cord (SpC, **Figure 2A-B, G-H**). Other examples in this category include AiE0387m, AiE0390h, AiE0390m, and the synthetic element ProB12^47^ (**Figure 2—figure supplement 1, 2**). “Regional” astrocyte enhancer-AAVs showed regionally restricted expression, such as AiE0385m that labeled astrocytes primarily in the telencephalon (**Figure 2C, F, Figure 2—figure supplement 1, 2**). Other “Regional” enhancer-AAVs labeled astrocytes primarily in subcortical domains, such as AiE0381h and AiE2160m (**Figure 2—figure supplement 1, 2**); while AiE0375m only labeled Bergmann glia, specialized astrocytes in the cerebellar cortex (CBX, **Figure 2—figure supplement 1, 2**). Interestingly, enhancers AiE2120m and AiE2160m labeled astrocytes in different regions, with some overlap in midbrain regions like inferior colliculus (**Figure 2—figure supplement 1, 2, 3**). “Sparse” enhancer-AAVs labeled astrocytes strongly but sparsely in most brain regions. These enhancer-AAVs include AiE0374m and its ortholog AiE0374h (**Figure 2—figure supplement 1, 2**). Enhancer-AAVs labeled as “Weak” gave astrocyte-specific patterns with low expression of SYFP (e.g., AiE0373m and AiE0386m, **Figure 2—figure supplement 1, 2**). Last, we designated several enhancer-AAVs as “Mixed specificities” because they labeled astrocytes in forebrain as designed, and populations of neurons in other brain regions. For example, AiE2118m labels many astrocytes strongly and specifically within the telencephalon, but also labels neurons strongly in non-telencephalic structures like midbrain (MB), deep cerebellar nuclei (CBN), and globus pallidus, external segment (GPe) (**Figure 2—figure supplement 1, 2**).

**Figure 2:**
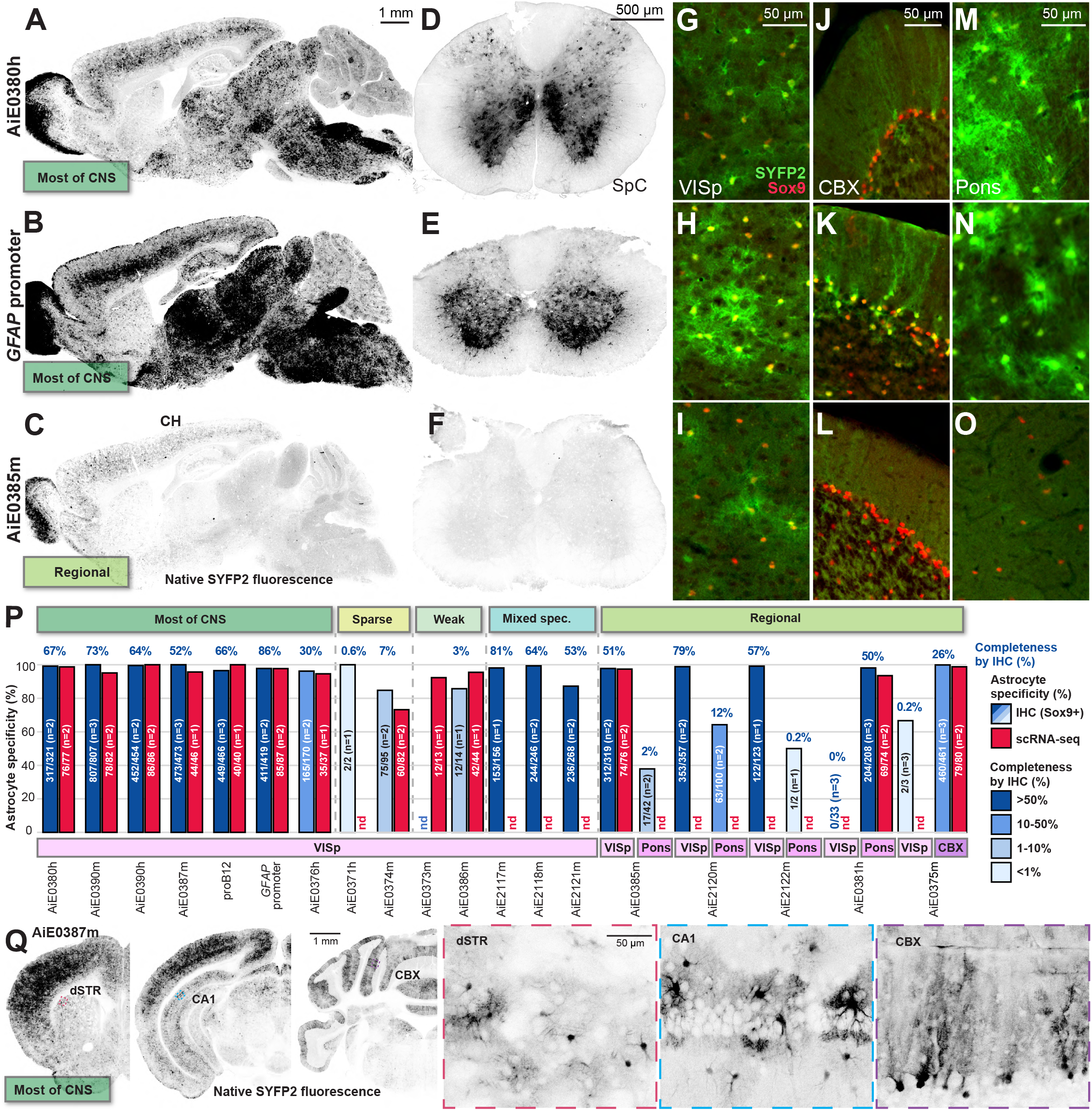
A collection of astrocyte-specific enhancer-AAV vectors with varying regional specificities and expression densities. (**A-B**) Astrocyte-specific enhancer-AAVs marking many astrocytes throughout most of the CNS. AiE0380h (**A**, n = 8 animals tested) and *GFAP* promoter (**B**, n = 3) mark many astrocytes throughout gray matter in FB, MB, HB, and CBX. (**C**) AiE0385m shows a regional pattern labeling astrocytes in cerebrum (CH) but not in MB, HB, or CBX (n = 6). (**D-F**) Astrocyte-specific enhancer-AAVs labeling astrocytes in lumbar SpC. AiE0380h (**D**) and *GFAP* promoter (**E**) label many astrocytes in SpC gray matter, but AiE0385m (**F**) does not label SpC astrocytes. (**G-O**) Positive confirmation of molecular astrocyte identity across brain regions. SYFP2+ astrocytes are colabeled with anti-Sox9 immunoreactivity in VISp, CBX, and Pons. Images in G-O represent n = 2 animals each assessed by IHC. (**P**) Quantification of specificity for astrocytes by astrocyte enhancer-AAVs. Specificity and completeness for astrocyte labeling by enhancer-AAVs was quantified by costaining with anti-Sox9 antibody in VISp, Pons, and CBX. Specificity is defined as the number of SYFP2+Sox9+ / total SYFP2+ cells x 100%. Completeness is defined as the number of SYFP2+Sox9+ / total Sox9+ cells x 100%, and is represented as a color scale on the bargraph with the exact value displayed as text above the bar. Brains from one to three mice per condition were analyzed, with range 131-827 cells counted (median 311) per brain region analyzed. AiE0375m-labeling was only quantified in the Purkinje cell layer of CBX, not in the granule or molecular layers. Specificity was also quantified by scRNA-seq, defined as the percentage of sorted SYFP2+ cells mapping as astrocytes within the VISp molecular taxonomy^97^. Overall, specificity is high for many astrocyte-specific vectors, with “Sparse” and “Weak” vectors showing low completeness, and “Regional” vectors showing more completeness in certain regions. One to three animals were analyzed per vector per modality, as indicated. The same IHC specificity data for *GFAP* promoter-SYFP2 are repeated in **Figure 6B**. (**Q**) Distinct astrocyte morphologies throughout the brain with AiE0387m enhancer-AAV targeting “Most of CNS”. Images were acquired on a serial two-phtoton tomography blockface imaging platform (STPT, TissueCyte, n = 4 tested). Abbreviations: CH cerebrum, dSTR dorsal striatum, CA1 cornu ammonis 1, CBX cerebellar cortex, SpC spinal cord, VISp primary visual cortex.

We quantified the specificity of many of these astrocyte-specific enhancer-AAVs using multiple independent techniques. First, we characterized SYFP2-expressing cells with immunohistochemistry (IHC) for Sox9, a marker of astrocytes throughout the brain^64^ (**Figure 2G-O**). Many of the astrocyte-specific enhancer-AAV vectors show high specificity, which we define as >80% specificity for the target cell population^39^. Astrocyte-specific enhancer-AAVs are usually >95% specific, and often >99% specific in VISp for Sox9-expressing astrocytes (**Figure 2P**). Second, we also observed high specificity when we isolated single SYFP2+ cells by flow cytometry and profiled them by scRNA-seq (**Figure 2P**).

Additionally, we assessed completeness of astrocyte labeling using IHC, and we observed that vectors scored as “Most of CNS” often label >50% of astrocytes in VISp, but “Weak” or “Sparse” vectors labeled many fewer astrocytes (**Figure 2P**). “Regional” vectors showed differing completeness across brain regions as expected (**Figure 2P**). Whole-brain serial two-photon tomography (STPT) of mouse brain transduced with astrocyte-specific enhancer-AAVs demonstrated distinct astrocyte morphologies in multiple brain regions (**Figure 2Q**). Thus, this collection of astrocyte-specific enhancer-AAVs are diverse with regard to the density of labeled cells, expression strength, and regionalization.

### A collection of oligodendrocyte-specific enhancer-AAVs

We screened 43 candidate oligodendrocyte enhancers, of which 21 (49%) gave oligodendrocyte-specific expression patterns (**Figure 3—figure supplement 1**). Unlike the astrocyte collection, the oligodendrocyte enhancer-AAVs all produced similar expression patterns throughout the gray matter and white matter tracts without any obvious regional specificity (**Figure 3A-F**). Oligodendrocyte-specific enhancer-AAV vectors ranged in expression from strong (for example AiE0410m, **Figure 3A**, AiE0641m, AiE0395h, and AiE0396h), to moderate (for example AiE0409h, **Figure 3B**, and the Myelin Basic Protein (*MBP*) promoter^31,65^, **Figure 3—figure supplement 1**), to weak (for example AiE0400h, **Figure 3C**). These vectors also labeled oligodendrocytes throughout the spinal cord (**Figure 3D-F**). We confirmed molecular oligodendrocyte characteristics of the vector-labeled cells by co-staining with CC1, a marker of oligodendrocytes^66^, which showed most vectors were highly specific across multiple brain regions (**Figure 3G-O**).

**Figure 3:**
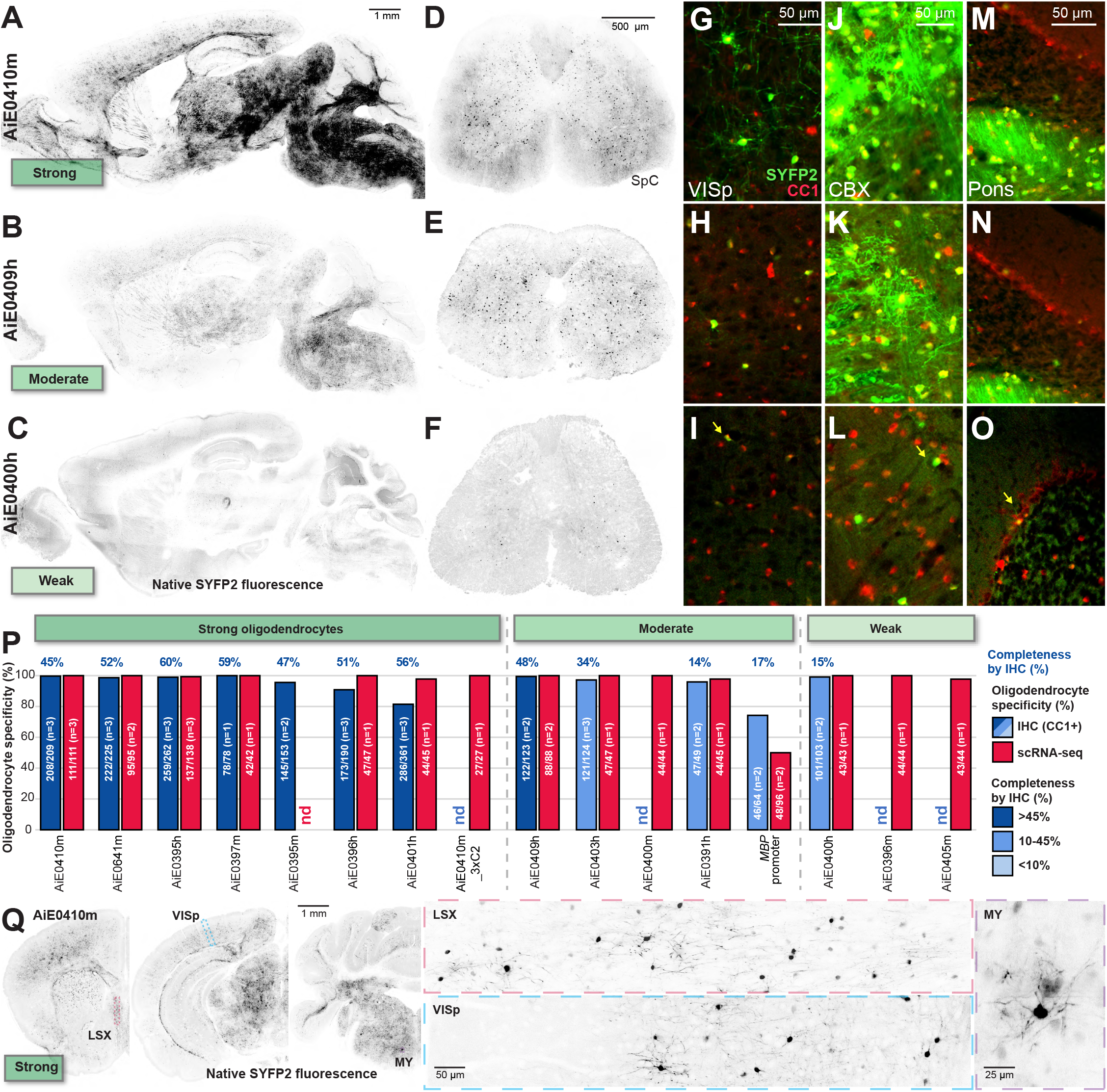
A collection of oligodendrocyte-specific enhancer-AAV vectors with varying levels of expression. (**A-C**) Oligodendrocyte enhancer-AAVs marking many oligodendrocytes throughout most of the CNS. AiE0410m (**A**, n = 7 animals tested), AiE0409h (**B**, n = 5), and AiE0400h (**C**, n = 4) label many oligodendrocytes throughout FB, MB, HB, and CBX, but at differing expression levels. (**D-F**) Oligodendrocyte enhancer-AAVs marking oligodendrocytes in lumbar SpC. AiE0410m (**D**), AiE0409h (**E**), and AiE0400h (**F**) mark oligodendrocytes in gray and white matter of SpC, but at different intensities. (**G-O**) Positive confirmation of molecular oligodendrocyte identity across brain regions. SYFP2+ oligodendrocytes are colabeled with CC1 immunoreactivity in VISp, CBX, and Pons. Two to three animals per enhancer-AAV were analyzed. **(P)** Quantification of specificity for oligodendrocytes by oligodendrocyte enhancer-AAVs. Specificity and completeness for oligodendrocyte labeling by enhancer-AAVs was quantified by costaining with CC1 antibody in VISp. Specificity is defined as the number of SYFP2+CC1+ / total SYFP2+ cells x 100%. Completeness is defined as the number of SYFP2+CC1+ / total CC1+ cells x 100%, and is represented as a color scale on the bargraph with the exact value displayed as text above the bar. Brains from one to three mice per condition were analyzed, with range 101-332 cells counted (median 147) per brain region analyzed. Specificity was also quantified by scRNA-seq, defined as the percentage of sorted SYFP2+ cells mapping as oligodendrocytes within the VISp molecular taxonomy^97^. One to three animals were analyzed per vector per modality, as indicated. Overall, specificity is high for many oligodendrocyte-specific vectors, with “Weak” vectors showing low completeness. **(Q)** Myelinating oligodendrocyte morphologies throughout the brain with AiE0410m. Sections were visualized with STPT (n = 2 tested). Abbreviations: SpC spinal cord, VISp primary visual cortex, CBX cerebellar cortex, LSX lateral septal complex, MY medulla.

Quantification by immunohistochemistry and scRNA-seq on sorted SYFP2-expressing cells showed >99% specificity and >45% completeness of labeling in VISp for multiple vectors (**Figure 3P**). STPT demonstrated myelinating oligodendrocyte morphologies in multiple parts of mouse brain (**Figure 3Q**). This collection of oligodendrocyte-specific enhancer-AAV vectors shows a diversity of expression strengths, but appears to label a homogeneous population of oligodendrocytes.

### Transcriptomic identities of astrocytes and oligodendrocytes

To investigate distinctions among enhancer-AAV-transduced cells, we performed SMARTerV4 scRNA-seq on sorted SYFP2-expressing cells. We characterized 2040 cells from 47 mice injected with 31 different enhancer-AAVs (1-2 mice per enhancer-AAV). After removing low-quality single-cell transcriptomes and cells not expressing the SYFP2 transcript, we focused the analysis on 1946 high-quality single cells. Astrocytes and oligodendrocytes separated in the UMAP space, as did astrocytes sorted from the distinct brain regions including VISp, midbrain/hindbrain (MB/HB), and CBX (**Figure 4A**). The molecular distinctions among regional astrocyte populations agree with findings from recent whole-brain atlases^24^. Indeed, mapping to a whole-brain taxonomic atlas indicates that, with high confidence, VISp-profiled astrocytes are predominantly mapped to the *Gja1*- and *Gfap*-expressing cluster “5112 Astro-TE NN_3”^24^, whereas MB/HB-profiled astrocytes marked by AiE0381h and AiE2160m mapped primarily to the *Gja1-* and *Agt*-expressing cluster “5109 Astro-NT NN_2”^24^. Likewise, the CBX-profiled Bergmann glia astrocytes mapped primarily to cluster identity “5102 Bergmann NN” as expected (**Figure 4B-D**). In contrast, labeled oligodendrocytes largely mapped to *Cldn11-* and *Mog*-expressing and most abundant oligodendrocyte cluster “5158 MOL NN”^24^ regardless of the enhancer used to label them (**Figure 4E**), confirming that oligodendrocyte enhancer-AAVs label a largely homogeneous population of oligodendrocytes.

**Figure 4:**
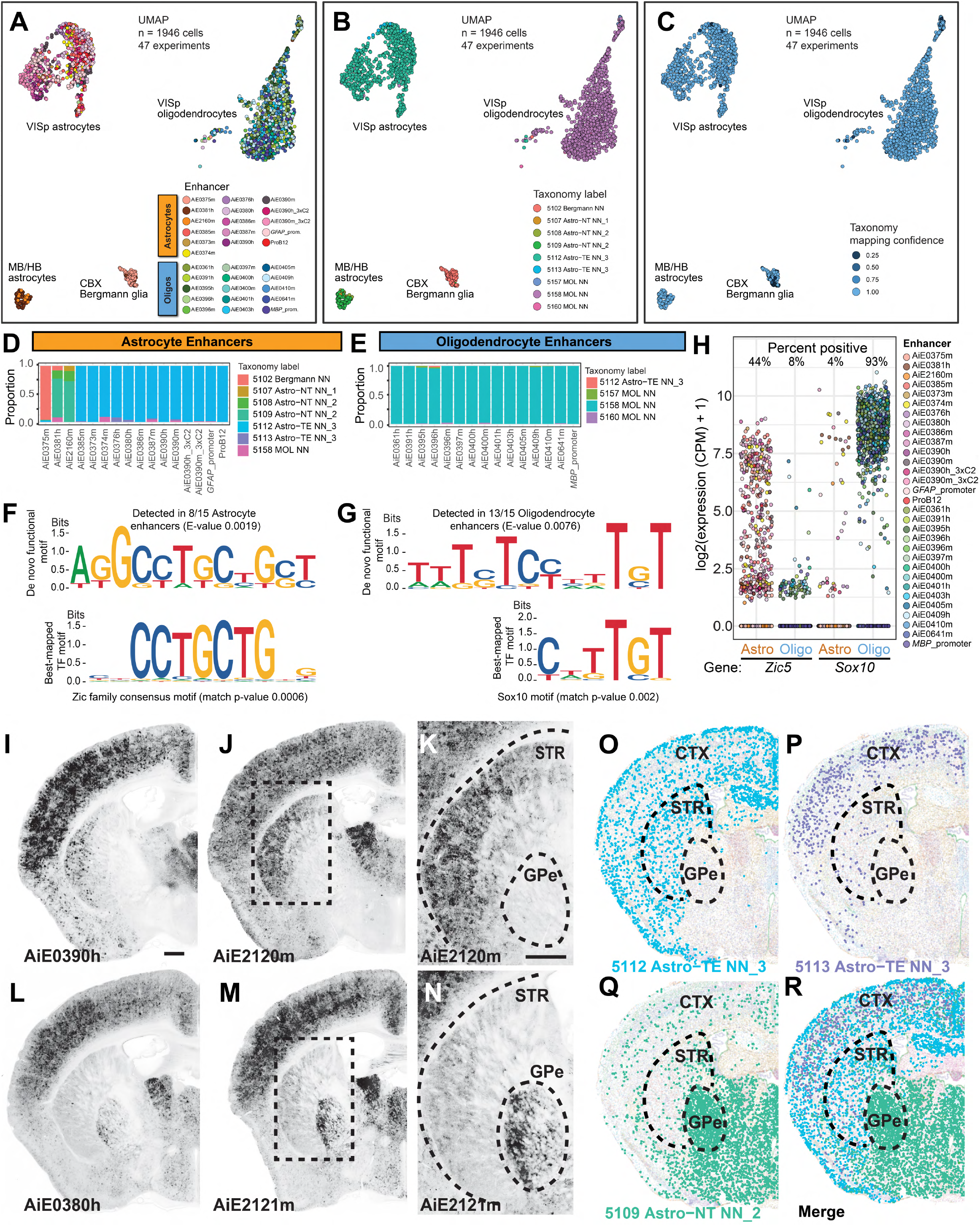
Transcriptomic identities of prospectively targeted astrocytes and oligodendrocytes. (**A-C**) Groups of transcriptomically profiled single cells, as visualized by UMAP. Single cells labeled by various astrocyte- and oligodendrocyte-specific enhancer-AAVs (n = 1946 quality-filtered cells) were profiled from 47 brains in 47 independent experiments by SMARTerV4^97^, one to three animals per enhancer. Libraries were aligned to mm10 and transformed into UMAP space for visualization, with coloring by enhancer (**A**), mapped taxonomic cell type cluster (**B**), and taxonomic mapping confidence (**C**). Overall CTX astrocytes group away from CTX oligodendrocytes as expected, and MB/HB astrocytes and Bergmann glia astrocytes group away from CTX astrocytes, consistent with recent results^24^. Note that AiE0381h- and AiE2160m-labeled astrocytes were dissected from MB/HB region, and AiE0375m-labeled Bergmann glia were dissected from CBX region, but the remainder of the cells were dissected fromVISp. (**D-E**) Quantifications of taxonomic cell type cluster mapping by enhancer vector. Prospectively labeled astrocytes from all enhancer-AAV vectors dissected from VISp predominantly map to cluster “5112 Astro-TE NN_3”, whereas those from MB/HB dissections (AiE0381h and AiE2160m) predominantly map to cluster “5109 Astro-NT NN_2”, and AiE0375m-labeled astrocytes from CBX dissections predominantly map to cluster “5102 Bergmann NN”. In contrast, all prospectively labeled oligodendrocytes predominantly map to cluster “5158 MOL NN”. Cluster identities are from a recent whole mouse brain taxonomy study^24^. (**F-H**) De novo motif detection in astrocyte- and oligodendrocyte-specific enhancer sequences using MEME-CHIP^67^ identifies one strong consensus motif in each set of sequences (top). These de novo motifs were mapped against databases of known TF motifs using TomTom (bottom), which identified the top hits as the Zic family consensus motifs for astrocytes, and Sox family motif for oligodendrocytes (Sox10 shown). These TFs (Zic5 and Sox10) show highly specific expression differences between astrocytes and oligodendrocytes from prospective scRNA-seq profiling (**H**). *** non-parametric Wilcoxon rank-sum test W = 577624 (*Zic5*) or W = 9838 (*Sox10*), p < 1e-16 each. (**I-N**) Intrinsic SYFP2 expression from the indicated enhancer-AAVs after retro-orbital administration. Images were generated by STPT. Boxes in I and L correspond to K and N, respectively. Scale in I and K is 500 µm. (**O-R**) MERFISH data showing the distribution of three astrocyte cell types revealed by single cell gene expression from the whole mouse brain^24^. Abbreviations: CTX cerebral cortex, STR striatum, GPe globus pallidus, external segment.

To understand the molecular regulation of these astrocyte and oligodendrocyte-selective enhancer-AAVs, we performed de novo motif detection on a collection of specific and strong astrocyte and oligodendrocyte enhancers (n = 15 each) using MEME-CHIP^67^. This analysis yielded one motif occurring in the majority of enhancers in each set (**Figure 4F, G**). Thes motifs had strong enrichments as measured by MEME-CHIP E-values less than 0.01, corresponding to the expected number of equally sized motifs of same or greater log likelihood ratio occurring in a set of random sequences of equal nucleotide content. We mapped these motifs against known transcription factor (TF) motif databases^68–70^, which revealed top matches to the Zic family consensus motifs for astrocytes (JASPAR accession numbers MA0697.2, MA1628.1, and MA1629.1; average of these three shown) and the Sox family motif for oligodendrocytes (JASPAR accession number MA0442.1 [Sox10 shown], and also Uniprobe accession numbers UP00030.1 and UP00062.1; **Figure 4F, G**). These analyses suggest that Zic and Sox family transcription factors might be key determinants of astrocyte versus oligodendrocyte identity in the CNS^71–74^, confirming the results of independent analysis of these sequences^42^. Moreover, Zic and Sox gene family members were differentially expressed between the profiled astrocytes and oligodendrocytes (*Zic5* 32-fold mean difference, *Sox10* 455-fold mean difference, p < 1e-16 each; **Figure 4H**). These results suggest Zic5 and Sox10 each play key roles in determining specificity of these glial enhancer-AAVs.

### Regional expression correlates with astrocyte cell type distribution

Using STPT imaging, we observed astrocyte-specific enhancer-AAVs to have two distinct expression patterns within the basal ganglia circuit. Several enhancer-AAVs showed elevated expression in astrocytes of the dorsolateral striatum and depletion in the globus pallidus (GP; **Figure 4I-K**), and several other enhancer-AAVs drove stronger transgene expression in astrocytes in the GP compared with those of the lateral striatum (**Figure 4L-N, Figure 2—figure supplement 2**). To determine if these enhancer-AAV expression patterns correspond to transcriptomically defined astrocyte cell types, we evaluated the spatial distributions of all astrocyte cell types in the Allen Brain Cell Whole Mouse Brain taxonomy^24^.

Interestingly, two closely related astrocyte cell types were strongly enriched in the dorsolateral striatum and cortex (“5112 Astro-TE NN_3” and “5113 Astro-TE NN_3”), while another was strongly enriched in the GP (“5109 Astro-NT NN_2”), demarcating the same boundaries observed with the collection of astrocyte enhancer-AAVs (**Figure 4O-R**).

### Measuring and optimizing enhancer strength

In some cases, enhancer-AAVs might not drive sufficient levels of a transgene to functionally affect the target cell. We sought to boost the expression levels of some enhancers by assembling concatemers of “core” sequences. These core sequences are responsible for the selective expression patterns and are often found in the central third of the original enhancer region identified by snATAC-seq^39^ (that is, ∼100-200 bp core from ∼300-600 bp original enhancer, **Figure 5A**). We observed that concatenation of the core can substantially increase expression from the original enhancer, such as AiE0387m concatenated to AiE0387m_3xC1 (**Figure 5B, C**), AiE0390h concatenated to AiE0390h_3xC2 (**Figure 5D, E**), or AiE0390m concatenated to AiE0390m_3xC2 (**Figure 5F, G**) while retaining similar expression patterns (**Figure 5H-J**) and cell type specificity (**Figure 5K**). However, concatenation sometimes resulted in a less dramatic effect on expression, (e.g., AiE0410m_3xC and AiE0641m_3xC; **Figure 3—figure supplement 1**).

**Figure 5:**
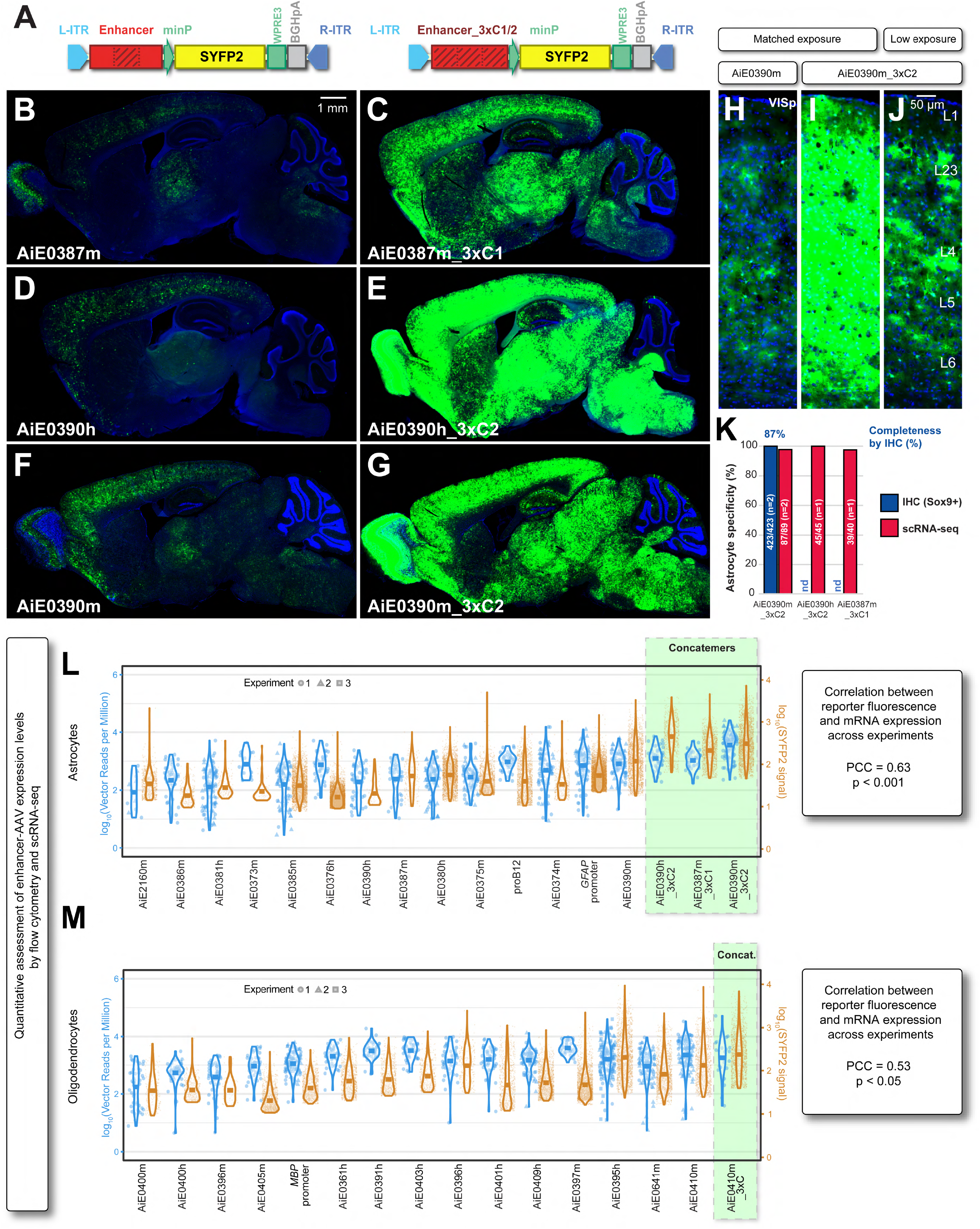
Optimizing astrocyte and oligodendrocyte enhancer strength. (**A**) Native Enhancer and for Enhancer_3xC2 vector designs. The central approximate third of the enhancer (the “Core2” element) is marked by dark hatches, and this element is triply concatemerized in the Enhancer_3xC2 vector. Alternatively, the first or third segment (“Core1” or “Core3”) may be concatemerized (determined empirically). (**B-J**) Dramatic increase in expression levels while maintaining specificity using Enhancer_3xC1/2 vector designs. Brains from mice injected with the Enhancer or Enhancer_3xC1/2 vectors were processed and imaged in parallel in these experiments. (**H-J**) Zoom in view of AiE0390m- and AiE0390m_3xC2-injected mouse VISp shows high specificity for morphological astrocytes throughout cortical layers in both cases. Data represent one to two animals per vector with parallel tissue processing and imaging. (**K**) Quantification of specificity for astrocytes by concatemer astrocyte enhancer-AAVs within VISp by IHC and scRNA-seq as described in **Figure 2P**. The same IHC data for AiE0390m_3xC2 are repeated in **Figure 6B**. (**L-M**) Direct correlated quantification of enhancer strength by flow cytometry and scRNA-seq, for both astrocyte-(**L**) and oligodendrocyte-specific (**M**) enhancer-AAVs. The left (blue) y-axis represents the log-transformed vector transgene reads per million in individual sorted scRNA-seq-profiled cells. The right (brown) y-axis represents the log-transformed SYFP2 signal intensity of positively gated vector-expressing cells observed on the flow cytometer, quantified as the fold signal of positive cells normalized to non-expressing cell autofluorescence (taken as background). Points represent individual cells observed by scRNA-seq and by flow cytometry, visualized also as violins, and with the horizontal bar representing mean expression levels across all cells expressing that enhancer-AAV, across one to three replicate experiments per vector. Across all experiments, we observe significant correlation between mean expression intensity at the RNA level by scRNA-seq, and mean SYFP2 reporter expression by signal intensity (astrocytes: n = 26 experiments, Pearson correlation coefficient [PCC] 0.63, t = 3.97, df = 24, p = 0.00057; oligodendrocyte n = 22 experiments, PCC 0.53, t = 2.82, df = 20, p = 0.011; correlation t-tests). Furthermore, 3xC astrocyte enhancers are among the strongest enhancers we have characterized, typically several fold stronger than their native counterparts. One to three animals were analyzed per enhancer-AAV vector (the same dataset described in **Figure 4A**). Abbreviations: VISp primary visual cortex.

We established single-cell measurements of reporter expression to compare enhancer strengths. We found that single-cell reporter fluorescence by flow cytometry correlated with vector read counts from SMARTerV4 scRNA-seq for both astrocytes and oligodendrocytes (astrocyte Pearson correlation coefficient = 0.63, p < 0.001; oligodendrocyte Pearson correlation coefficient = 0.53, p < 0.05; **Figure 5L, M**, **Figure 5—figure supplement 1**). These measurements revealed that several concatenated enhancer-AAVs, including AiE0390m_3xC2 and AiE0390h_3xC2, drove the strongest expression among the vectors we have tested (**Figure 5L**), consistent with the microscopy results (**Figure 5B-J**). Conversely, MB/HB (AiE0381h and AiE2160m) and Bergmann glia (AiE0375m) astrocyte enhancers have among the weakest expression levels we have tested (**Figure 5L**), likely a consequence of selecting glial enhancers from cortical accessibility datasets.

### Enhancers can yield more robust specific expression across transgenes than classic promoters

Previously it was demonstrated that the human *GFAP* promoter (GfaABC1D^63^) can change specificity from astrocytes to neurons over time when driving certain difficult transgene cargos including NeuroD1-mCherry fusion protein^50^. Accordingly we tested whether astrocyte-specific enhancers might be more robust to transgenes than *GFAP* promoter. As demonstrated previously, we found that at seventeen days post transduction, *GFAP*-driven NeuroD1-mCherry is expressed predominantly in NeuN+ neurons in primary visual cortex (VISp, 90 ± 2% of labeled cells, mean ± standard deviation), unlike *GFAP*-driven SYFP2 which is expressed with high Sox9+ astrocyte specificity (98 ± 2% of labeled cells, **Figure 6A-B**, p < 0.001, see also **Figure 2**). In sharp contrast an optimized astrocyte-specific enhancer, AiE0390m_3xC2 maintains high levels of astrocyte specificity whether driving SYFP2 (100 ± 0% of labeled cells) or NeuroD1-mCherry (95 ± 4% of labeled cells, **Figure 6A-B**, p < 1e-7, see also **Figure 5K**). These results suggest that optimized astrocyte-specific enhancers may provide more predictable expression for certain transgenes, and confirm the previous results that astrocytic NeuroD1-mCherry expression alone is insufficient to drive astrocyte transdifferentiation to neurons^50^.

**Figure 6:**
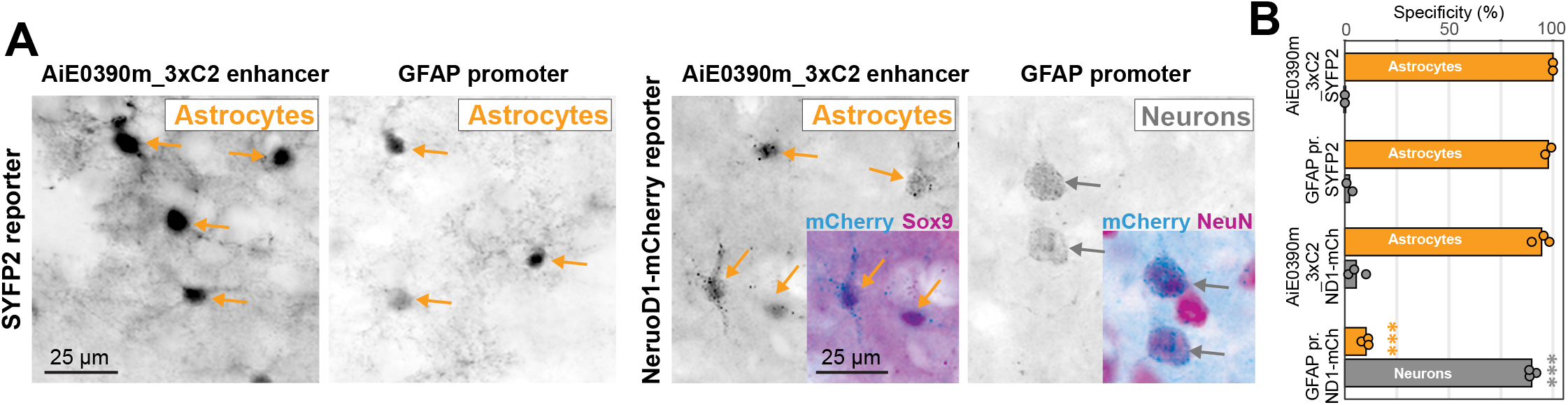
Robust specificities of astrocyte enhancer-AAVs with a difficult transgene cargo. (**A-B**) Specific delivery of a difficult transgene cargo with an optimized astrocyte-specific transgene. Mice received the indicated enhancer-AAVs driving either SYFP2 or NeuroD1-mCherry by intravenous PHP.eB AAVs (5e11 gc per animal), and expression analyzed in brain by IHC with NeuN neuronal marker or Sox9 astrocyte marker. Specificity for neurons and astrocytes was quantified as (Reporter+[NeuN or Sox9]+ cells) / (All Reporter+ cells) in primary visual cortex (VISp, Layer 5 cells shown, n=2-3 animals per condition). The same IHC specificity data for *GFAP* promoter-SYFP2 and AiE0390m_3xC2-SYFP2 are repeated in **Figures 2P and 5K**, respectively. Two to three animals were analyzed per condition (each dot represents one animal). *** ANOVA with Tukey’s post hoc adjustment for multiple comparisons, F = 647, p < 1e-7, for *GFAP* promoter-driven NeuroD1-mCherry compared to every other condition.

### Predictability of enhancer-AAV expression across tissues and in a disease state

Recent work suggests that AAV-mediated transduction and high transgene expression in organs such as the liver and dorsal root ganglia is associated with toxicity^35,75^. We tested if we could predict off-target activity from enhancer accessibility profiles in a human body-wide epigenetic dataset^76^. Different astrocyte-specific enhancers showed either moderate or low accessibility across many body organs (**Figure 7A**), since this was not a factor in their selection for testing. We assessed off-target transgene expression in liver after intravenous delivery since PHP.eB capsid transduces the liver^77^. We observed that astrocyte enhancers with moderate levels of accessibility in liver (AiE0381h, AiE0371m, AiE0371h, and AiE0386m) expressed SYFP2 in many hepatocytes, whereas the enhancers with little or no liver accessibility (AiE0387m, AiE0375m, AiE0390h, and AiE0390m) expressed SYFP2 in many fewer hepatocytes (p < 0.001, **Figure 7B-C**), demonstrating strong predictability of enhancer expression profiles in off-target tissues. In contrast, the *GFAP* promoter can drive bright expression in few hepatocytes, which is not predictable from any epigenetic or transcriptomic atlases.

**Figure 7:**
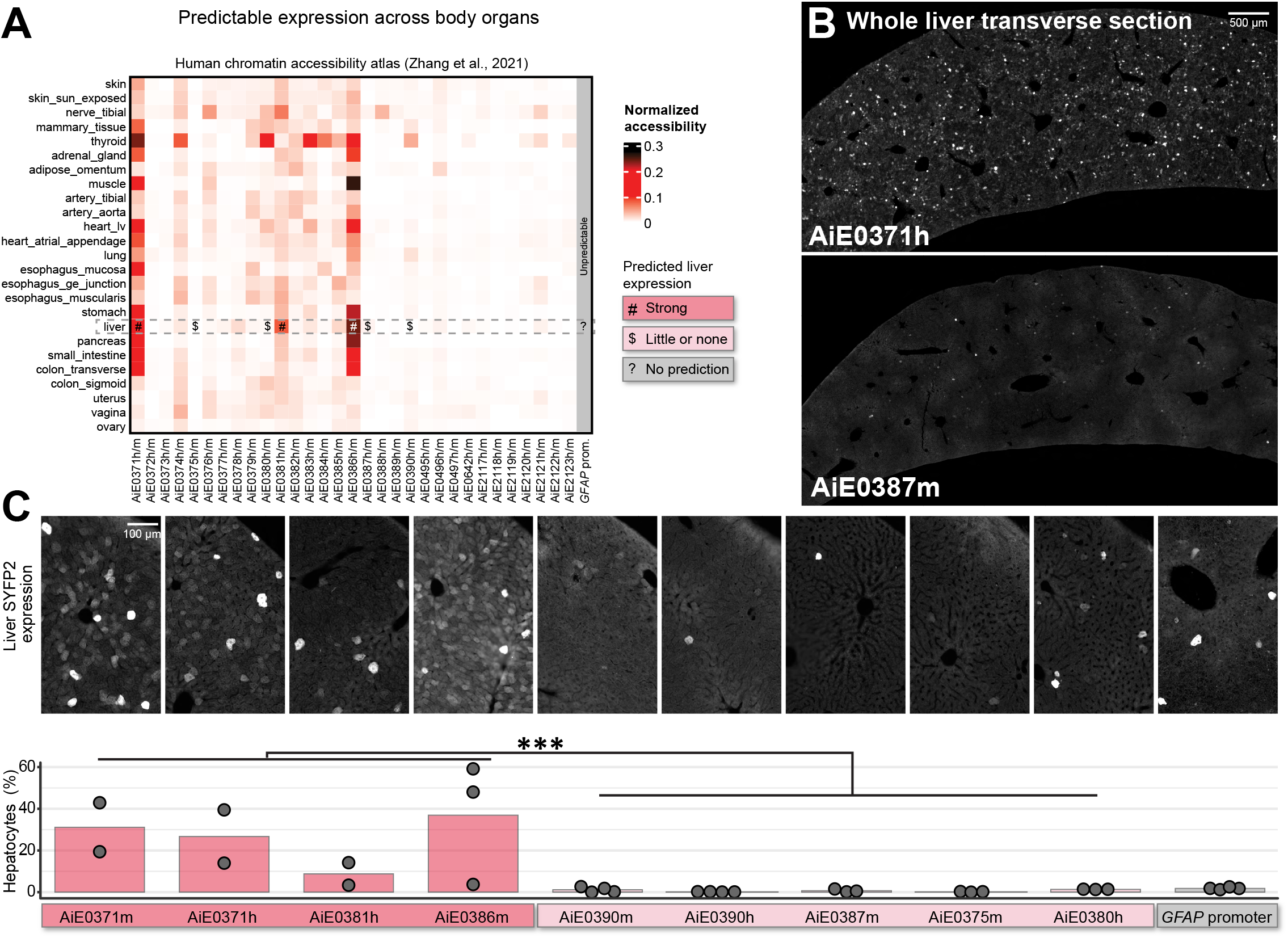
Predictability of astrocyte enhancer-AAV expression patterns across body organs. **(A)** Accessibility profiles of astrocyte-specific enhancers in the human whole-body accessibility atlas^76^. Single-cell profiles were grouped within each tissue into pseudo-bulk aggregates, then normalized according to the signal (reads in peaks) within the dataset. Accessibility profiles are likely to predict enhancer activities within each tissue. Focusing on liver, some astrocyte-specific enhancers are predicted to have “Strong” expression, and some are predicted to have very “Little or none” expression. In contrast, accessibility atlases do not predict expression of *GFAP* promoter across tissues. **(B)** Whole livers from mice injected intravenously with AiE0371h- and AiE0387m-enhancer-AAV vectors, stained with anti-GFP antibody. AiE0371h has high liver accessibility, is predicted to have high liver expression, and shows many strong SYFP2-expressing hepatocytes throughout the liver as predicted. In contrast, AiE0387m has little liver accessibility and so is predicted to have little liver expression, and in fact shows few positive SYFP2-expressing hepatocytes as predicted. **(C)** Agreement between liver expression predictions and liver expression measurements across several astrocyte-specific enhancer-AAV vectors. AiE0371m, AiE0371h, AiE0381h, and AiE0386m all show many SYFP2-expressing hepatocytes as predicted. AiE0390m, AiE0390h, AiE0375m, AiE0387m, and AiE0380h all show few weak SYFP2-expressing hepatocytes as predicted. *GFAP* promoter shows few but strongly expressing hepatocytes, which was not predictable from the accessibility atlases. Liver images in **B** and **C** represent two to four mice analyzed for each vector (each dot represents one mouse). *** non-parametric Wilcoxon rank sum test, W = 153, p < 1e-4 for the comparison of Strong versus “Little or none”-predicted enhancers.

*GFAP* expression can change expression in the context of disease or injury^14,78^, and the synthetic *GFAP* promoter can change specificity when delivering different transgenes^50–52^, suggesting this might have limitations for genetic access to astrocytes in disease. We compared *GFAP* promoter and one of the best enhancer-AAVs (AiE0390m) in *Scn1a^+/-^* Dravet syndrome model mice since they have strong epilepsy-associated reactive astrogliosis^79^. We injected SYFP2-expressing enhancer-AAVs into these mice prior to the Dravet syndrome critical period of high susceptibility to seizures and mortality at P21 and analyzed tissues at P42 (**Figure 8A**). Significant hippocampal reactive astrogliosis was seen in Dravet syndrome model mice, revealed by elevated endogenous GFAP immunoreactivity in all hippocampal layers (**Figure 8B**). Concomitant with this reactive astrogliosis, the *GFAP* promoter-driven AAV reporter changed its expression pattern. Normally, this promoter drives moderate levels of astrocyte-specific reporter expression. However, in the Dravet mice experiencing epilepsy, expression strength in astrocytes was considerably elevated, and ectopic expression was observed in many dentate gyrus granule cell neurons (DGCs, 12 ± 6% of DGCs labeled, p < 0.01, **Figure 8B, C**). In sharp contrast, the AiE0390m enhancer-AAV vector maintains astrocyte specificity at moderate levels despite the profound reactive astrogliosis in these diseased animals (0 ± 0% of DGCs labeled, **Figure 8B, C**).

**Figure 8:**
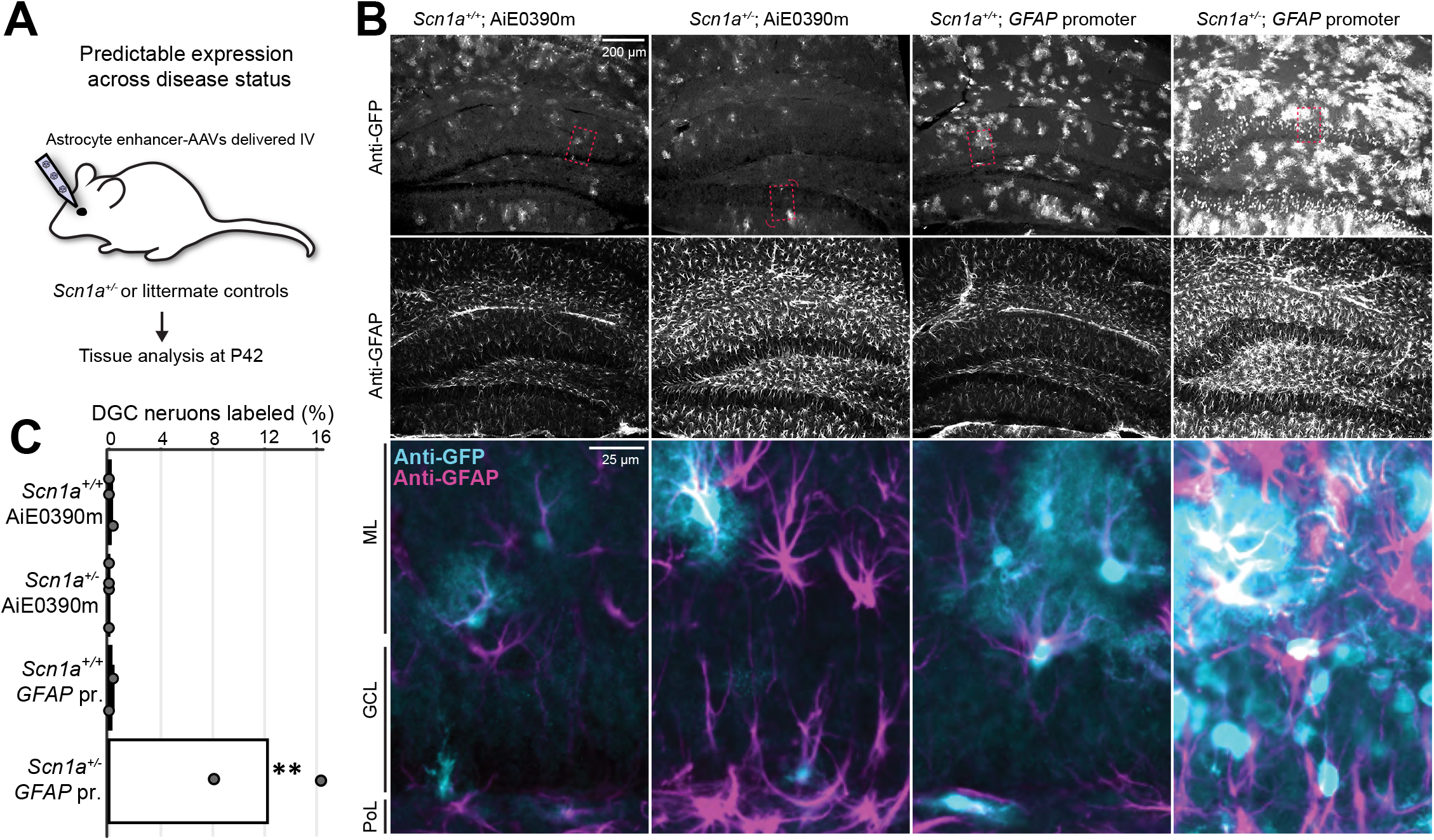
Predictability of astrocyte enhancer-AAV expression patterns in a mouse model of epilepsy. (**A-C**) Testing fidelity of enhancer-AAV expression in a mouse model of epilepsy. We used Dravet syndrome model *Scn1a^+/-^* mice to induce epilepsy-associated hippocampal reactive astrogliosis, injected enhancer-AAVs prior to the critical period, and analyzed tissue for expression patterns after the critical period (**A**). We assessed hippocampal reactive astrogliosis with anti-GFAP antibody and enhancer-AAV expression with anti-GFP antibody (**B**). AiE0390m maintained specific expression and similar levels in hippocampal astrocytes regardless of epileptic reactive astrogliosis. In contrast, *GFAP* promoter expression strongly increased in reactive astrocytes, and also was observed in dentate gyrus granule cells. Red dashed rectangles indicate the position of the expanded zoomed view, and the curved red arrows indicate a rotated view for one zoom. (**C**) Quantification of DGC labeling reveals a much greater number of DGC neurons labeled with GFAP promoter in Dravet syndrome model mice. Two to four animals were analyzed per condition (each dot represents one mouse). ** ANOVA with Tukey’s post hoc adjustments, F = 16, p < 0.01 for the *GFAP* promoter in *Scn1a^+/-^* mice compared to each other condition. Abbreviations: ML molecular layer, GCL granule cell layer, PoL polymorphic layer.

### Astrocyte specific sensing of cholinergic signals in the nucleus accumbens during behavior

We next asked whether the vectors would drive sufficient expression to obtain functional signals in a cell-type specific manner. We created a vector driving astrocyte-specific expression of the acetylcholine indicator iAChSnFR^53^ (**Figure 9A,B**), to detect extracellular acetylcholine fluctuations in the nucleus accumbens (NAc) in an awake and behaving animal using fiber photometry. After stereotaxic injection, we implanted optical fibers above the injection site to perform fiber photometry. We trained mice to perform a dynamic foraging reinforcement learning task while we recorded photometry signals to assess bulk acetylcholine fluctuations in the NAc. In the task, water-restricted mice chose freely between two lick ports for a water reward after an auditory cue. Reward probabilities of the two lick ports were changed in a block-design manner, which resulted in both rewarded and unrewarded trials (**Figure 9C**). During these trials, the astrocyte-specific iAChSnFR vector drove sufficient expression to observe fluorescence intensity fluctuations (**Figure 9D**) which can be seen to differ during individual rewarded and unrewarded licks (**Figure 9D**, bottom left). Both rewarded and unrewarded trials showed an increase in fluorescent signal at the time of choice, followed by a deviation in signal depending on whether the trial was rewarded (**Figure 9E**). Astrocyte acetylcholine signals decreased more in rewarded trials than in unrewarded trials (**Figure 9E**). In summary, these results indicate that glial-selective enhancer-AAVs can be applied to measure functional acetylcholine dynamics in the NAc.

**Figure 9:**
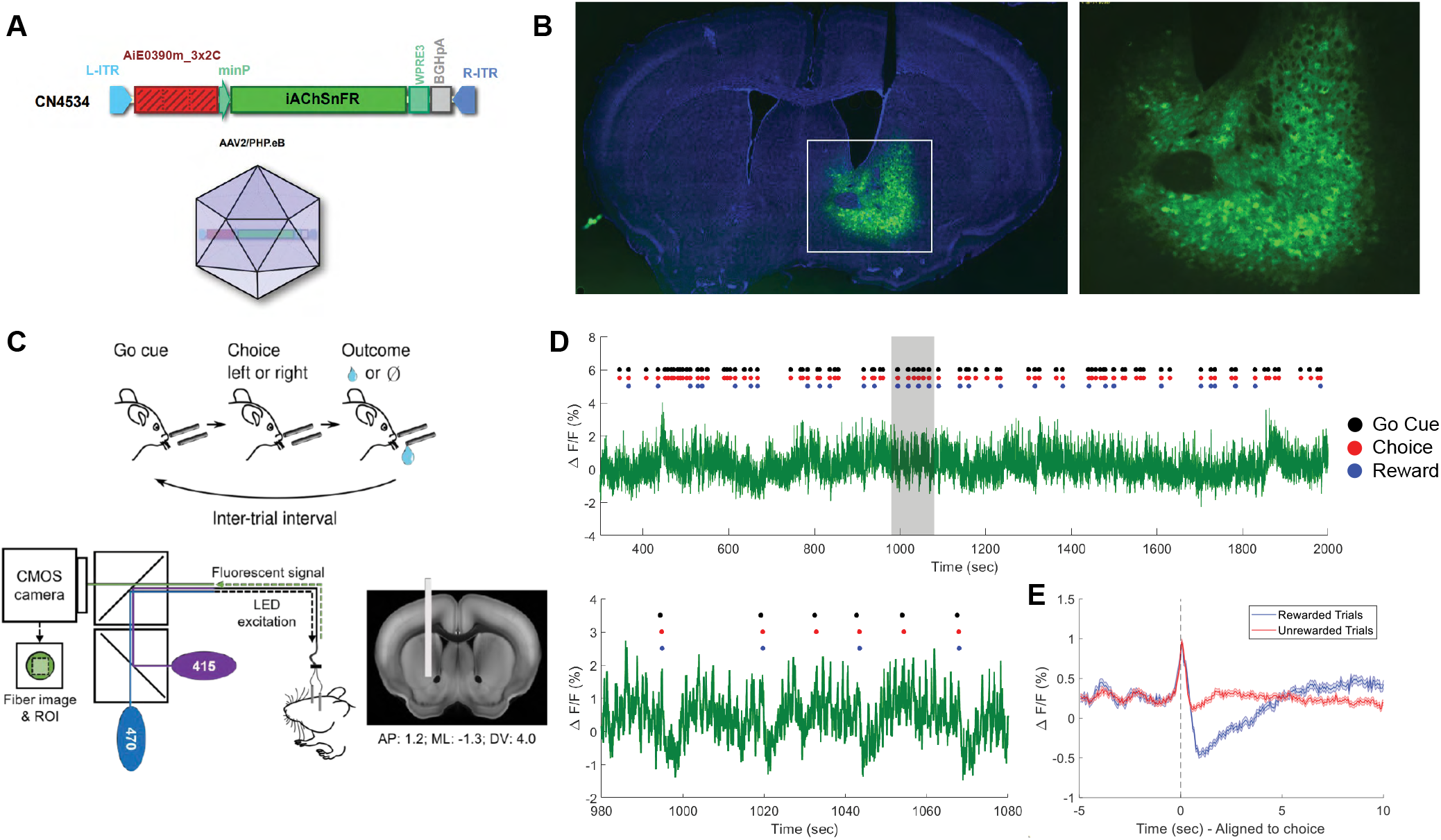
Astrocyte specific sensing of cholinergic signals in the nucleus accumbens during behavior. **(A)** AiE0390m_3xC2 driving expression of iAChSnFR. Enhancer vector is cloned into a pAAV plasmid and packaged into PHP.eB AAVs. **(B)** Coronal section showing stereotaxic injection of enhancer virus expressing iAChSnFR in the nucleus accumbens (injection coordinates: AP: 1.2, ML: 1.3, DV: 4.1). **(C)** Behavior and imaging experiment setup. Top: dynamic foraging behavior task schematic. Bottom: Fiber photometry instrumentation schematic and fiber location in a coronal section. **(D)** Fiber photometry signals of acetylcholine fluctuations during task performance. Top: ∼30 min segment of a 2-hour session of dynamic foraging. Black dots represent the auditory cue, red dots represent time of first lick, blue dots represent water reward delivery. Bottom: 100 second (980-1080 seconds) zoom in on above session with 6 individual trials (4 rewarded and 2 unrewarded trials). Data from one mouse shown, representative of three mice assayed. **(E)** Trial-averaged signals of rewarded and unrewarded trials aligned to time of first lick (mean±sem).

### Astrocyte- and oligodendrocyte-specific AAV-Cre vectors

AAVs that can selectively drive Cre recombinases are valuable tools for mouse genetics since the AAV can be delivered somatically for cell type-specific recombination of floxed alleles. We used some of the best astrocyte- and oligodendrocyte-specific enhancers to express an attenuated R297T mutant Cre recombinase in *Ai14* reporter mice^80,81^ (**Figure 10A**). AiE0390m_3xC2-driven Cre vector drove astrocyte-specific recombination in many parts of mouse brain, including specific (95 ± 3%) and complete (98 ± 1%) labeling in VISp gray matter astrocytes (**Figure 10B-D**). Similarly, an AiE0410m-driven Cre vector also incorporating three miR126 binding sites to prevent unwanted endothelial cell recombination^82^ drove oligodendrocyte-specific recombination in many parts of mouse brain, including specific (97 ± 3%) and complete (81 ± 0.1%) labeling in VISp gray matter oligodendrocytes (**Figure 10E-G**). Additional vectors utilizing AiE0390m and AiE0387m also showed specific and complete astrocyte labeling in VISp (**Figure 10H-I**). These tools may permit fine interrogation of the roles of astrocyte and oligodendrocyte gene expression in brain biology.

**Figure 10:**
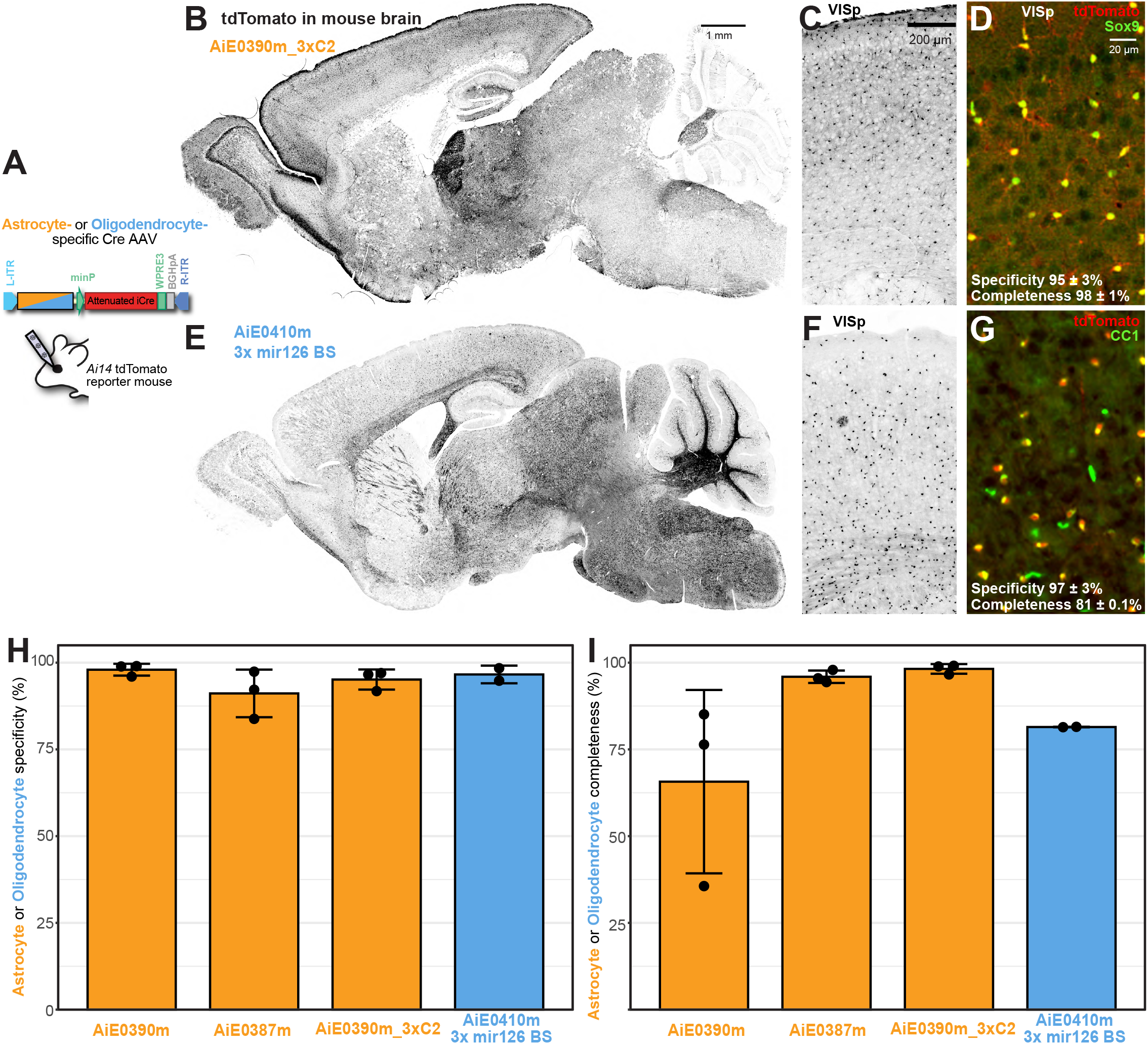
Astrocyte- and oligodendrocyte-specific Cre AAVs. (**A**) Design and testing of astrocyte- and oligodendrocyte-specific attenuated Cre-expressing enhancer-AAVs. (**B-D**) Specific Cre recombination in astrocytes driven by AiE0390m_3xC2. Ai14 reporter recombination is observed in astrocytes throughout the mouse brain (**B**). Recombination within VISp is highly specific (**C**), as evidenced by highly specific and complete recombination in Sox9+ astrocytes (**D**). (**E-G**) Specific Cre recombination in astrocytes driven by AiE0410m. This vector also includes three 3’UTR mir126 binding sites to eliminate unwanted expression in mir126-expressing endothelial cells^82^. Ai14 reporter recombination is observed in oligodendrocytes throughout the mouse brain (**E**). Recombination within VISp is highly specific (**F**), as evidenced by highly specific and complete recombination in CC1+ oligodendrocytes (**G**). (**H-I**) Quantification of recombination specificity (**H**) and completeness (**I**). Specificity is defined as (tdTomato+Sox9 or CC1+ cells) / (all tdTomato+ cells) x 100%. Completeness is defined as (tdTomato+Sox9 or CC1+ cells) / (all Sox9 or CC1+ cells) x 100%. Two to three animals were analyzed for each condition (each dot represents one mouse).

### Cross-species genetic access to astrocytes and oligodendrocytes

We tested whether several glial-selective enhancer-AAV vectors could maintain specific expression across species. We first tested conservation in neonatal rats after ICV administration (**Figure 11A**). We observed that an optimized AiE0390m_3xC2 vector containing 4X2C miRNA binding sites to prevent any unwanted expression in excitatory neurons^83^ labeled rat cortical astrocytes with high specificity (98% specific, **Figure 11B, C**). We also found that an AiE0641m-driven AAV vector labeled rat cortical oligodendrocytes with high specificity (91% specific, **Figure 11D**). Note that astrocyte labeling completeness and spread to caudal brain structures could not be assessed since ICV administered virus resulted in uneven spread. Thus, some vectors identified in the mouse screen maintained specificity in rats after ICV injection into neonates, consistent with the results from other enhancer-AAV collections^44^.

**Figure 11:**
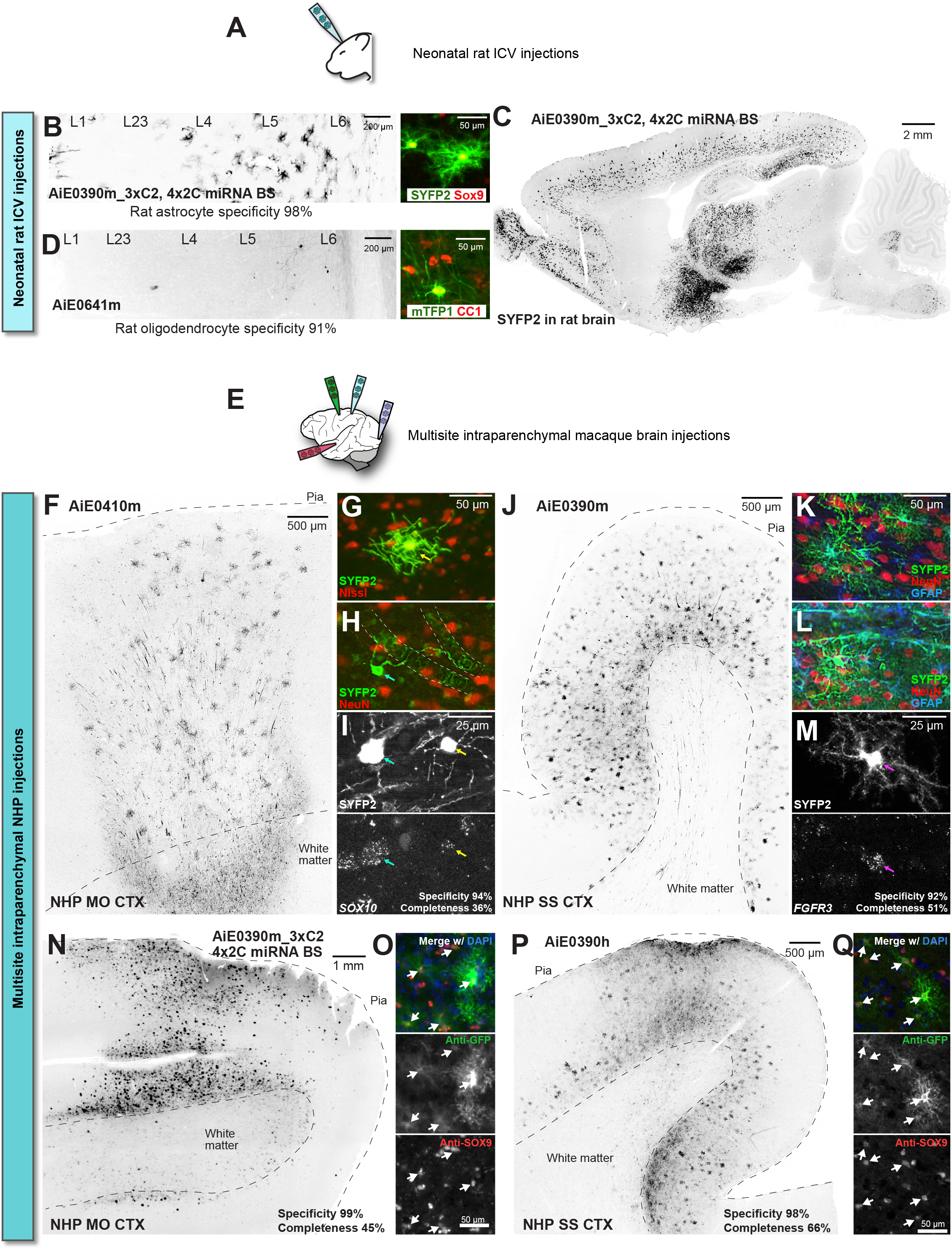
Genetic targeting of astrocytes and oligodendrocytes across species. (**A**) Testing enhancer-AAV vectors by neonatal rat ICV injections. (**B-C**) Validation of astrocyte-specific enhancer-AAV vectors in rat by IHC with Sox9 antibody. AiE0390m_3xC2 vector also incorporates 4X2C 3’UTR miRNA binding sites to prevent any off-target labeling in excitatory neurons^83^. Each image represents one animal tested. (**D**) Validation of oligodendrocyte-specific enhancer-AAV vector in rat. AiE0641m shows specific expression in CC1+ VISp oligodendrocytes. Each image represents one animal tested. (**E**) Multiple stereotactic intraparenchymal injections into macaque brain. (**F-I**) Prospective labeling of macaque oligodendrocytes in vivo. AiE0410m enhancer-AAV vector gives widespread labeling of oligodendrocytes throughout the depth of motor cortex (**F**). Most labeled macaque oligodendrocytes exhibit multipolar ramified morphology indicative of local axon myelination. (**G**). Some labeled oligodendrocytes exhibit morphologies suggesting wrapping around wider tubular structures highlighted with dashed white lines (**H**). SYFP2-expressing cells of both morphological types express the oligodendrocyte/OPC marker *SOX10* with high specificity (**I**). One animal tested. (**J-Q**) Prospective labeling of macaque astrocytes in vivo. AiE0390m enhancer-AAV vector gives widespread labeling of astrocytes throughout the depth of somatosensory cortex (**J**). A few large Layer 5 pyramidal neurons are also labeled. Labeled astrocytes show the expected bushy morphology and GFAP immunoreactivity of astrocytes in parenchyma (**K**) and sometimes reside near walls of large-diameter tubular structures (**L**). SYFP2-expressing astrocytes express the astrocyte marker *FGFR3* with high specificity (**M**). In another animal, an injection of AiE0390m_3xC2 vector with 4X2C 3’UTR miRNA binding sites gives high specificity (99%, **N**) for SOX9+ astrocytes in MO (**O**), without any large Layer 5 pyramidal neurons labeled. In another animal, AiE0390h vector gives high specificity (98%) of labeling for SOX9+ astrocytes (**P-Q**). Each vector represents one animal tested. Abbreviations: VISp primary visual cortex, MO motor cortex, SS somatosensory cortex.

We extended these cross-species tests to primate, using multisite intraparenchymal injections into *Macaca nemestrina* (pig-tailed macaque, **Figure 11E**). We administered AiE0410m AAV vector into motor (MO) cortex and observed cell morphologies of myelinating oligodendrocytes throughout the cortical column (**Figure 11F-G**). Interestingly, we also observed SYFP2-expressing cells with a different morphology: one to three processes that spiral around stretches of tubular structures approximately 15-20 microns in diameter often running perpendicular to the cortical pial surface (**Figure 11H**). These tubular structures have not yet been defined and were not observed in mouse or rat testing, but both morphological types of SYFP2-expressing cells co-expressed the oligodendrocyte marker *SOX10* by mFISH with high specificity (94%, **Figure 11I**).

We also injected the somatosensory cortex (SS) with the astrocyte-specific AiE0390m AAV vector and observed many SYFP2-expressing cells with astrocyte morphology throughout the cortical column (**Figure 11J**; note a small number of large layer 5 pyramidal neurons labeled as well, which we did not observe in mouse testing). SYFP2-expressing astrocytes co-expressed GFAP either in parenchyma (**Figure 11K**) or in apposition to a large blood vessel (**Figure 11L**), and some showed fibrous morphology in white matter (**Figure 11—figure supplement 1**). Enhancer-AAV-labeled astrocytes also expressed the astrocyte-specific transcript *FGFR3* with high specificity (92%) and about half of the *FGFR3+* gray matter astrocytes were labeled through the whole cortical depth near the injection site (51%) (**Figure 11M**). In an additional animal we also tested AiE0390m_3xC2 vector containing 4X2C miRNA binding sites in MO, and we observed 99% specificity for SOX9+ astrocytes and no layer 5 pyramidal neuron labeling (**Figure 11N-O**), and a final injection into a third animal also confirmed 98% specificity of AiE0390h-driven vector for SOX9+ astrocytes in SS (**Figure 11P-Q**). These studies suggest that enhancer-AAV vectors can provide specific and dependable genetic access to astrocytes and oligodendrocytes across multiple species and reveal morphological glial features not observed in the mouse.

## Discussion

Flexible and dependable tools to target glial cell populations will be essential to understand their diverse roles in brain biology. Here we characterize a collection of astrocyte- and oligodendrocyte-specific enhancers that can be used in AAV vectors and applied across species. Some of these enhancer-AAV vectors were also reported in a recent publication (Ben-Simon et al.,^41^), and here we demonstrate deeper characterization of this subset. Most of these enhancer-AAVs generated highly specific labeling of astrocytes or oligodendrocytes, and often substantial completeness of labeling. Detailed characterization revealed the enhancers showed a range of expression strengths, and the astrocyte enhancers frequently exhibited regional enrichment and differences in labeling densities. We demonstrate that this enhancer-AAV toolset can be applied: 1) across species in mouse, rat and macaque, 2) in a model of epilepsy in mice where reactive astrogliosis is occurring without losing specificity, and 3) to deliver Cre selectively to many astrocytes and oligodendrocytes, 4) irrespective of transgene delivered, and 5) to measure circuit dynamics with a neurotransmitter sensor *in vivo*. As a result, these glial enhancer-AAVs will be useful for interrogating the roles of these glial cell types in health and disease.

### Lesson learned from screening

Several lessons were learned through the process of screening for astrocyte and oligodendrocyte enhancer-AAVs. First, multiple selection criteria can identify strong glial enhancer-AAVs. Excellent functional enhancers were derived from genome-wide peak selection across mouse and human datasets of distinct epigenetic modalities, peak selection from one mouse dataset based on peak strength, or peak selection only near marker genes. Second, screening one enhancer-AAV at a time can be efficient for the identification of useful enhancer-AAVs. Nearly half of the candidate enhancers proved to be specific for the targeted type, and this created a large and diverse library of new enhancer-AAVs that labeled astrocytes or oligodendrocytes in different ways. Third, while enhancers were identified from neocortex at the subclass level, brain-wide patterns match well to whole brain cell gene expression atlas patterns^24^. Specifically, AAV-labeled oligodendrocytes appear homogeneous across the brain, while astrocytes show prominent regional enrichments and differences in cell density. Fourth, body-wide specificity of enhancer-AAV expression can be predicted based on body-wide epigenetic datasets^76^. Thus, off-target expression can be predicted and limited during the enhancer selection process, and enhancers can be identified that label different cell types throughout the body. These attributes will make enhancer-AAVs a valuable tool for precision gene therapy where only the cell population of interest is expressing the therapeutic transgene.

### Enhancer-AAVs show diverse expression patterns

We tested a large collection of astrocyte-selective enhancer-AAVs and saw a diversity of expression patterns. Some astrocyte enhancer-AAVs predominantly labeled in telencephalic structures, some in MB/HB structures, and others showed “Sparse” but uniform expression. Within the forebrain, we also observed multiple enhancer-AAVs that showed mutually exclusive enriched or depleted expression in dorsolateral striatum or the globus pallidus. These regional differences were reflected by region-specific astrocyte transcriptomic profiles in agreement with recent results^24^. The GPe astrocytes express high levels of GABA uptake gene *Slc6a11*^22,84,85^ while striatal and cortical (but not GPe) astrocytes express high levels of glutamate uptake gene *Slc1a2*^22,85,86^. This suggests that the astrocytes help selectively maintain glutamate or GABA tone depending on the local neurotransmitter balance. The functional roles of astrocytes in different brain regions will require additional experiments that this collection of enhancer-AAVs may facilitate, with individual or even multiplexed perturbations of multiple astrocyte populations simultaneously. Similarly, it is not yet clear what produces “Sparse” astrocyte reporter expression, and the answer will await future experiments.

### Enhancer-AAVs can be optimized to improve vector function

Achieving functional levels of transgene expression is critical for applying enhancer-AAV tools to learn new biology and deliver gene therapies. We show that two astrocyte enhancers could be optimized through generation of a triple concatenated core to produce significantly higher levels of transgene expression without sacrificing astrocyte specificity. One such optimized tool was used to detect acetylcholine activity in the NAc. The NAc has been implicated in reward-related reinforcement learning and receives inputs from dopaminergic and serotonergic neurons, in addition to local cholinergic signaling^87–90^. We used enhancer-AAVs to deliver the acetylcholine indicator iAChSnFR to astrocytes within the NAc, and measured dynamics with fiber photometry. This experiment showed that the astrocyte enhancer AiE0390m_3xC2 maintained faithful astrocyte specificity after direct injection into NAc and expressed sufficient iAChSnFR for sensing of acetylcholine in live awake animals. It also showed that iAChSnFR expressed by astrocytes could readily detect acetylcholine dynamics in NAc, and that astrocytes are a good cellular compartment for the sensor.

Different optimization efforts enabled the generation of a functional Cre-AAV that was specific for astrocytes in most parts of the brain. Cre recombinases can lose specificity when expressed from enhancer-AAVs, possibly due to low-level expression of this potent enzyme^36^. We were able to successfully generate an astrocyte-selective Cre AAV using an attenuated Cre recombinase with the R297T mutation^80,81^, in combination with the AiE0390m_3xC2 enhancer. As a result we produced a highly specific somatic astrocyte Cre. Similarly we were able to produce a highly specific oligodendrocyte Cre AAV using the AiE0410m enhancer along with miR126 binding sites. We anticipate these vectors may unlock many types of functional experiments dissecting the roles of these important glial cell populations.

### Enhancers-AAVs can be identified that have conserved specificity from mouse to macaque

Some astrocyte and oligodendrocyte enhancer-AAVs are active and selective across species from mouse through macaque. This property will allow these tools to be applied somatically in multiple organisms besides mice. Testing enhancers in macaques revealed interesting morphological differences compared to mouse. Abundant mature oligodendrocytes with large dendritic arbors were labeled in both mouse and primate tissues with AiE0410m, but this enhancer-AAV also labeled *SOX10*+ cells wrapping around radial tubes (presumably blood vessels) in primate but not mouse. Also, the astrocyte enhancer AiE0390m labeled abundant protoplasmic astrocytes in the gray matter in mouse and macaque, but also labeled several large fibrous astrocytes that were not observed in the mouse experiments. The ability to function across species and label cells that are not obviously represented in mouse tissue, makes this collection of enhancer-AAVs a powerful toolset to better understand new biology. It also makes a compelling case that some of these enhancer-AAVs could be suitable for use in human gene therapy where astrocyte or oligodendrocyte expression selectivity is required.

### Enhancer-AAVs can be cell state-independent

AAVs that drive expression using promoters can cause gene expression changes in a state dependent fashion. We showed that the *GFAP* promoter changed expression strength and specificity in the context of epilepsy-induced reactive astrogliosis using an *Scn1a* haploinsufficiency model. This is not surprising since the *GFAP* gene is known to change in response to disease and injury^12,78^. Additionally, *GFAP* promoter-driven expression changed specificity from astrocytes to neurons with the difficult transgene NeuroD1-mCherry, as seen in other reports^50^.

Enhancer-AAVs utilizing AiE0390m, on the other hand, did not show a change of expression in either epileptic reactive astrogliosis conditions or due to differing transgenes. This could be due to the enhancer being selected based on core cell type identity, and most properties of cell types have not been seen to change character dramatically in the context of disease^26,91,92^. Enhancers can be selected that do not change activity during development, aging, or disease, in a similar way to avoid selecting enhancers predicted to have activity in off-target tissues. Alternatively, enhancers can be identified to change with disease state as desired^93^. As epigenetic datasets are expanding to cover these axes of disease, development, and aging, it is becoming feasible to select only putative enhancers with the desired activity profiles.

### Conclusion

We have characterized a large collection of enhancer-AAV vectors for targeting astrocytes and oligodendrocytes. These vectors will provide researchers with the ability to mark and manipulate these critical cell types in a variety of species, genetic backgrounds, ages, and disease contexts, and could also enable delivery of therapeutics. Combined with other recently discovered AAV-based tools^31,32,37–44,46,49,77^, this glial-targeting toolbox will help to advance the understanding of the roles of glial cell types in brain biology, make the complex cellular anatomy of the brain more experimentally tractable, and advance the development of AAV-based therapeutics for human CNS disorders.

## Methods

### Epigenetic analysis and enhancer nomination

We identified candidate astrocyte- and oligodendrocyte-specific enhancers from cortical epigenetic datasets. We used the following datasets: human middle temporal gyrus snATAC-seq^39^, mouse primary visual cortex scATAC-seq^40^, human frontal cortex snmC-seq^58^, mouse frontal cortex snmC-seq^58^, human frontal cortex bulk mC-seq^57^, and mouse frontal cortex bulk mC-seq^57^. A single cell glial snmC-seq dataset^59,60^ became available only after initial identification of most of the enhancers described in this study. From the single nucleus/cell ATAC-seq datasets, we aggregated reads according to cell subclass as in the references, and then called peaks using Homer findPeaks (http://homer.ucsd.edu/homer/) with the-region flag, yielding typically tens of thousands of peaks per subclass, sized approximately 300-600 bp, as previously described^39^. To find differentially methylated regions (DMRs) we either used the published regions by Luo et al. 2017 in Extended Data Tables 5 and 6^58^, and aggregated by subclass and then to all neurons using bedtools merge (https://bedtools.readthedocs.io/en/latest/). Alternatively for bulk non-neuronal DMRs we used methylpy DMRfind with minimum differentially methylated sites set to 1 on the dataset of Lister 2013, as previously described^39,58^. To convert mouse and human peak or DMR regions to each other’s genomic coordinates for direct intersectional analysis, we used liftOver (https://genome.ucsc.edu/cgi-bin/hgLiftOver) with minMatch parameter set to 0.6. All peak regions described in this manuscript successfully liftOver from human to mouse, and vice-versa, except AiE0733m which does not have an obvious human ortholog via liftOver.

To automatically identify peaks and DMRs genome-wide that are astrocyte- or oligodendrocyte-specific within each dataset, we used a series of bedtools intersectBed (https://bedtools.readthedocs.io/en/latest/) operations to filter for regions that are only detected in astrocytes or oligodendrocytes. For the “high specificity” criteria, we found peaks that were specifically detected in both human and mouse cortical astrocytes/oligodendrocytes^39,40^, but did not overlap DMRs from either human or mouse neurons of any subclass^58^, and these candidate enhancers are marked by gold square icons in **Figure 1**. These criteria yielded a set of 87 candidate astrocyte-specific enhancers and 112 candidate oligodendrocyte-specific enhancers, and the top 17 (Astrocyte) or 16 (Oligodendrocyte) candidate enhancers were chosen from this list as ranked by Homer findPeaks score. Homer findPeaks score is a measure of peak significance relative to local background, not peak strength. Additionally, a small number of these “high specificity” criteria candidate enhancers also overlapped with DMRs from both human and mouse non-neuronal cells^57^ (3 Astrocyte and 4 Oligodendrocyte), and these are marked by gold star icons in **Figure 1** and **Figure 1—figure supplement 1**.

For the “high strength” criteria we found peaks that were specifically detectable in cortical astrocytes, using mouse scATAC-seq data only^40^, and agnostic to detection in human and methylation datasets. This analysis yielded 2119 (astrocyte) and 3940 (oligodendrocyte) candidate enhancers, which were ranked by read counts within the region, and the top 7 candidate enhancers for astrocytes from this list were chosen. This ranking led to nomination of peaks that are overall stronger and longer, and these candidate enhancers are marked with a purple triangle in **Figure 1B**, but accessibility profiles were not always conserved in human tissue, as shown in **Figure 1—figure supplement 1A,C**.

Some candidate enhancers were identified manually in the vicinity of known astrocyte or oligodendrocyte marker genes by visual inspection of ATAC-seq read pileups on UCSC browser (marked as “*M”* for Marker genes in **Figure 1B**). Methylation data was not visualized in this manual nomination process. Importantly we found that both automatic and manual approaches can identify peaks with high strength and specificity, as shown in **Figure 1D-E**.

Additionally, enhancer AiE2160m was initially identified as a candidate enhancer for pericytes in cortex using the data of Graybuck et al.^40^, but it was found in the course of this study to instead label mid/hindbrain astrocytes.

To model enhancer screening results as a generalized linear model, we confined analysis to 50 screened enhancers where we observed a clear yes/no screening result for both itself and its cross-species ortholog. These candidate enhancers were AiE0371, 372, 373, 375, 377, 379, 380, 382, 383, 384, 388, 393, 394, 398, 399, 401, 406, 407, 408, 374, 376, 390, 395, 409, and 410, both the m and horthologs for each. For each of these genomic regions we calculated candidate enhancer strength (read CPM within either astrocytes or oligodendrocytes), candidate enhancer specificity (defined as the proportion of astrocyte or oligodendrocyte enhancer strength relative to the summed strength in all populations, using the data of Mich et al. or Graybuck and Daigle et al.^39,40^), candidate enhancer length in base pairs, region-segmented PhyloP using the previous method^39^, and tabulated whether each candidate enhancer’s partner ortholog worked (binary yes [1] or no [0]). We fit a logistic generalized linear model of testing results from these predictors using glm() in R with the following command:

`glm(Screen_result_01 ∼ Length + PhyloP + Specificity + Strength_cpm + Ortholog_result_01, family=binomial(link=’logit’), data = data)`

The significance of each coefficient to predict the screening result was determined from the coefficients of the model, using the data as provided in **Table S1**. Although high peak specificity and strength were important criteria for candidate enhancer identification, these metrics each had little predictive power to explain success or failure of screening collection testing as evidenced by coefficients of fit to a logistic linear model (**Table S1**; strength z-value = 1.43, *p* = 0.15; specificity [defined as proportional strength within target cell subclass] z-value = −1.20, *p* = 0.23), similar to enhancer length (z-value = −0.13, *p* = 0.89), enhancer sequence conservation measured by PhyloP (z-value = 0.87, *p* = 0.38), and the presence of a functional ortholog in testing (z-value = 1.77, *p* = 0.077), which suggests that there are additional undiscovered elements that determine successes versus failures in AAV-based enhancer screening. Overall, the null deviance was 67.3 on 49 degrees of freedom, and the residual deviance was 57.7 on 44 degrees of freedom, again indicating little power of these features to predict the screening results.

### Cloning and packaging enhancer-AAVs

With candidate enhancers chosen, we next found their predicted DNA sequence from genomic reference sequence using Bioconductor package Bsgenome^94^. We extracted the sequence and padded 50 bp to each side of the enhancer to provide room for forward and reverse primer binding sites that capture the entire enhancer. From these padded sequences, we used automatic primer design in Geneious to identify primer pairs within the 50 bp pads to specifically amplify each enhancer, and append a constant 5’ homology arm to each enhancer for automatic Gibson assembly into reporter-AAV plasmid. We amplified the regions from C57Bl/6 tail snip DNA or from human male genomic DNA (Promega catalog # G1471) using FastPhusion 2x Master Mix (Thermo Fisher catalog # F548L), and >90% of the PCR reactions were successful on the first try. In some cases we redesigned primers to attempt a second amplification.

We cloned into reporter backbone CN1244 (Addgene plasmid #163493) using the sites MluI/SacI and the 5’ primer homology arms F: TTCCTGCGGCCGCACGCGT and R: GACTTTTATGCCCAGCCCGAGCTC, using Infusion kit (Takara catalog # 638949). For some enhancers we instead cloned into a next generation reporter vector backbone that includes a SYFP2-P2A-3xFLAG-H2B reporter for detection of cytosolic SYFP2 and nuclear FLAG for simultaneous expression analysis and snRNA-seq, using the same cut sites and homology arms (see **Table S1**). We transformed infusion reactions into Mix N’ Go (Zymo Research catalog # T3001) chemically competent Stbl3 E. coli (Thermo Fisher catalog # C737303) and selected on 100 µg/mL carbenicillin plates. We cultured individual clones at 32 C, verified them by Sanger sequencing, maxiprepped them with 100 µg/mL ampicillin, and saved them as frozen glycerol stocks at −80°C.

We used maxiprep DNA for packaging into PHP.eB AAV particles. For routine enhancer-AAV screening by intravenous delivery in mouse we generated small-scale crude AAV preps by transfecting 15 µg maxiprep enhancer-reporter DNA,15 µg PHP.eB cap plasmid, and 30 µg pHelper plasmid into one 15-cm dish of confluent HEK-293T cells using PEI-Max (Polysciences Inc. catalog # 24765-1). After transfection the next day we changed the medium to 1% FBS, and after 5 days the cells and supernatant were collected, freeze-thawed 3x to release AAV particles, treated with benzonase (1 µL) for 1 hr to degrade free DNA, then clarified (3000g 10min) and then concentrated to approximately 150 µL by using an Amicon Ultra-15 centrifugal filter unit at 5000g for 30-60 min (NMWL 100 kDa, Sigma #Z740210-24EA), yielding a titer of approximately 5e13 genome copies (gc)/mL. Titer values ranged from 4.47e12 to 1.47e14 gc/mL, interquartile range 3.13e13-6.55e13 gc/mL, mean 5.10e13 gc/mL, n = 215 total viral preps. For large-scale gradient preps for intraparenchymal injection into macaque or mouse or ICV injection into rat, we transfected 10 15-cm plates of cells, and also purified preps by iodixanol gradient centrifugation. We assessed viral titer for both crude and gradient AAV preps by digital droplet PCR on a BioRad QX200 system. All vectors showing specific expression patterns are available through Addgene.

### Optimizing enhancer strength through concatemerization

For some native enhancers that showed specific expression patterns, we sought to boost their expression levels through concatemerization. To concatemerize, we segmented the enhancer (typically approximately 400-600 bp) into approximately thirds with approximately 25 bp of overlaps at the junctions (each a candidate “core” of approximately 200 bp), then designed a tandem array of three cores in series (approximately 600 bp). These synthetic tandem array sequences were gene synthesized by Azenta/GeneWiz PRIORITYGene synthesis service with flanking MluI/SacI sites for restriction enzyme digestion and ligation into corresponding sites in CN1244. We then packaged and tested concatemerized PHP.eB enhancer-AAVs as above.

### Mice and injections

All mouse experimentation was approved by Allen Institute Institutional Animal Care and Use Committee (IACUC) as part of protocol #2020-2002. In these studies, we purchased C57Bl/6J mice from The Jackson Laboratory (Stock # 000664). For enhancer screening these C57Bl/6J mice were injected with AAVs in the retro-orbital sinus at age P21 with 5E11 genome copies of AAV/PHP.eB viral vectors with brief isoflurane anesthesia. For enhancer validation studies (IHC) mice were injected the same way but between ages P42 to P56. Tissues from mice were harvested at 3 to 4 weeks post injection for analysis, except in the case of the NeuroD1-mCherry experiments (**Figure 1F**) where mice were analyzed at 17 days post injection as previously reported^50^, and in the liver experiments (**Figure 6C**) in which 1-2 mice for each vector were analyzed at 3 weeks and 1-2 mice were analyzed at 36 weeks to confirm durability across age. We perfused animals with saline then 4%PFA in PBS, and harvested brains or other tissues and post-fixed in 4%PFA overnight, before rinsing and cryoprotecting in 30% sucrose solution before sectioning at 30 micron thickness on sliding microtome with a freezing stage. PFA used in these experiments was prepared by dissolving solid PFA (VWR Chemicals #28794.295) in PBS by heating, which was then stored in frozen aliquots. For enhancer screening we counterstained with DAPI and propidium iodide and mounted in Vectashield Vybrance, and imaged on either a Nikon Ti-Eclipse or Nikon Ti-Eclipse 2 epifluorescent microscope, Olympus FV-3000 confocal microscope, or Leica Aperio slide scanner. In some experiments where noted, we tested enhancer-AAVs after bilateral intracerebroventricular (ICV) injection at age P2 using the technique of Kim et al.^95^ These ICV-injected pups were harvested for tissue analysis at age P21. For whole brain imaging of expression pattern, we performed sequential blockface imaging of brains using the TissueCyte 1000 serial two-photon tomography system^96^.

For testing in Dravet syndrome model mice, 129S1/SvImJ-*Scn1a^em1Dsf/J^* mice (strain # 034129) were purchased from Jackson Laboratories and bred to C57Bl/6J mice to create *Scn1a^R613X/+^* pups on a F1 hybrid C57Bl/6J:129S1/SvImJ background, and these pups were injected retro-orbitally at P21 with tissue analysis at P42. Additionally, we also tested enhancer-AAV vectors in *CMV-Cre;Scn1a^A1783V/+^* pups on C57Bl/6J background, which were generated from crossing B6(Cg)-*Scn1a^tm^*^1^*^.1Dsf/J^* male mice (The Jackson Laboratory, strain #:026133) with homozygous CMV-Cre female mice (*B6.C-Tg(CMV-cre)1Cgn/J*, The Jackson Laboratory, strain # 006054). Data from both lines appeared similar, so we combined them for analysis.

### Mouse immunohistochemistry (IHC)

For IHC and ISH, we transcardially perfused mice with ice-cold 25 mL HBSS (Thermo Fisher Scientific # 14175079) containing 0.25 mM EDTA (Thermo Fisher Scientific # AM9260G), followed by 12 mL of ice-cold 4% paraformaldehyde in 1x PBS, freshly prepared from 16% PFA (Electron Microscopy Sciences #15710). We dissected brains and other tissues from carcasses and post-fixed them at 4 degrees overnight, and the next morning we rinsed the tissues with fresh PBS, and then transferred to 30% sucrose solution in PBS for cryoprotection. For sectioning on Leica CM3050 cryostat, we then embedded tissues in OCT cryo-compound (Tissue-Tek # 4583) at room temperature at least 3 hours, then froze the blocks on dry ice and stored at −80°C until sectioning at 25 micron thickness. Alternatively, we sectioned half-brains at 25 µm thickness on frozen 30% sucrose solution slabs on a sliding microtome (Leica SM2000R) equipped with freezing stage. Sections were stored at 4 degrees in PBS containing 0.1% sodium azide until analysis. IHC of liver was performed as for brain, focusing on the frontal lobe.

For IHC we used the following antibodies: chicken anti-GFP (Aves # GFP-1010, 1/3000), rabbit anti-Sox9 (Cell Signaling clone D8G8H, # 82630S), mouse CC1 antibody (Abcam # ab16794, 1/1000), mouse anti-GFAP (Millipore Sigma clone G-A-5, # G3893, 1/1000), with 5% normal goat serum (Thermo Fisher Scientific # 31872) and 0.1% Triton X-100 (VWR 97062-208) for blocking and permeabilization, and appropriate Alexa Fluor-conjugated secondary antibodies for detection.

### Flow cytometry and single cell transcriptomics

We prepared cell suspensions for flow cytometry and single cell RNA-seq from brain tissue as previously described^97^. Briefly, for flow cytometry, we perfused mice transcardially under anesthesia with ACSF.1. We harvested the brains, embedded in 2% agarose in PBS, then sliced thick 350 micron sections using a compresstome with blockface imaging, then picked the sections containing the region of interest (VISp, or mid- and hindbrain, or cerebellar cortex), and dissected out the regions of interest. We then treated dissected tissues with 30U/mL papain (Worthington LK003176) in ACSF.1 containing 30% trehalose (ACSF.1T) in a dry oven at 35°C for 30 minutes. After papain treatment we quenched digestion with ACSF.1T containing 0.2% BSA, triturated sequentially using fire-polished glass pipettes with 600, 300, and 150 micron bores, filtered the released cell suspensions into ACSF.1T containing 1% BSA, centrifuged cells at 100g for 10 min, then resuspended cells in ACSF.1T containing 0.2% BSA and 1 μg/mL DAPI prior to flow cytometry and sorting on a FACSAria III (Becton-Dickinson). SYFP2 reporter brightness was measured as the ratio of positive cell population mean fluorescence intensity, divided by the low mean fluorescence intensity of autofluorescence in non-expressing cells. This measure of reporter brightness is more consistent than positive cell population mean fluorescence intensity alone, due to differences in raw signal across days, cytometers, and cytometer settings.

For single cell RNA-seq, we sorted single SYFP2+ cells into tubes and processed them via SMARTer v4 using the workflow described previously^97^, on 47 enhancer-AAV-injected mice. In each experiment from one mouse injected with one single enhancer-AAV we sorted and profiled up to 48 cells per experiment, and each measurement was taken from a distinct individual cell. After retroorbital injections, enhancer-AAV SYFP2-expressing cells consisted of on average 7% of the positive brain cells (range 0.1-20.1% of cells, n = 47 experiments). We sequenced single cell-derived SMARTer libraries at 659996 ± 199038 (mean ± standard deviation) reads per library on an Illumina NovaSeq instrument at Broad Institute (Cambridge, MA) or on an Illumina NextSeq instrument at Allen Institute (Seattle, WA). We aligned the libraries to mm10 genome using STAR (https://github.com/alexdobin/STAR), and also aligned them to the synthetic AAV transgene reference construct using bowtie2 (https://bowtie-bio.sourceforge.net/bowtie2/index.shtml). From 2040 initial cells, we excluded from analysis the libraries with poor library quality metrics, consisting of: firstly low-quantity or degraded libraries (judged as less than 65% percentage of cDNA library sized greater than 400 bp, consisting of 71 [3.4%] libraries in this study), and secondly those that lacked AAV transgene-mapping reads (likely mis-sorted events, 23 [1.1%] of remaining libraries). Applying these filtering criteria yielded a dataset for analysis of 1946 high-quality AAV transgene-expressing cells, with alignment rates of 92 ± 3% to mm10 genome and 4654 ± 1285 genes detected per cell (mean ± standard deviation). To assess enhancer specificity within the cortex we mapped the high quality transgene-expressing SMARTer cells to the SMARTer-based VISp cellular taxonomy generated by Tasic et al. using bootstrapped hierarchical approximate nearest neighbor mapping^24,97^, and quantified the specificity as the percentage of positively sorted cells that mapped to the expected cell subclass (astrocytes or oligodendrocytes). To test for significance of correlation of brightness by flow cytometry with expression levels by scRNA-seq, we calculated Pearson’s product-moment correlation coefficient by cor.test() function in R.

To understand different characteristics of different regional astrocyte populations we utilized scrattch.mapping (https://github.com/AllenInstitute/scrattch) from the Allen Institute. To accomplish this we first transformed these cells by principal component analysis and performed UMAP dimensionality reduction on the first 40 principal components for visualization using the default scanpy parameters, which clearly separated oligodendrocytes and regional groupings of astrocytes. For clustering astrocytes we subset the dataset to astrocytes only, then identified the top 2000 genes ranked by variance among them, recomputed UMAP projections from these high-variance genes, then performed Leiden clustering^98^ which identified VISp, MB/HB, and CBX astrocyte clusters as expected, and finally identified differential genes among them (differential gene expression threshold false discovery rate less than 5% and log_2_-fold change greater than 0.5) using scanpy (https://scanpy.readthedocs.io/en/stable/). In doing so we detected two major subgroups of VISp astrocytes that are distinguished by presence or absence of immediate-early gene markers (for example, *Fos*, *Fosl2*, *Nr4a1*, *Irs2*, *Pde10a*, and *Pde7b*). This distinction may be an artifact of the cell dissociation process for scRNA-seq; for the purposes of this study we collapse these cortical astrocyte clusters. In order to understand the different regional characteristics of astrocyte populations we mapped cells to whole brain taxonomy we mapped to the best-correlated mean-aggregated taxonomic cluster from the ABC-WMB atlas^24^ with 100 bootstrapped iterations using the top 10% of high-variance genes and omitting a variable number of genes (10-50%) each round. We interpret the frequency of correct mapping rounds as the mapping confidence. We also used CELLxGENE for single cell visualization (https://github.com/chanzuckerberg/cellxgene). Spatial transcriptomic analysis was performed as described in the recent whole brain transcriptomic taxonomy study^24^, and cell type location data was visualized using Cirrocumulus (https://cirrocumulus.readthedocs.io/en/latest/index.html).

For determination of *Zic5* and *Sox10* differential gene expression between astrocytes and oligodendrocytes, we used two-sided ANOVA on expression measurements from individually profiled cells from all the experiments with no exclusion, and no covariates were tested. Testing for normality by the Shapiro-Wilk test revealed that *Zic5* and *Sox10* expression are not normally distributed (*Zic5* W = 0.539, p-value < 2.2e-16; *Sox10* W = 0.958, p-value = 5.1e-16), so we used a non-parametric Wilcoxon rank-sum test. No significance thresholds adjustments were made for multiple comparisons since only one comparison was performed. For the comparison of *Sox10* versus *Zic5* expression (mean counts per million +/- standard deviation, n cells): astrocyte *Zic5* expression 32 ± 71, n = 864; astrocyte *Sox10* expression 0.3 ± 5, n = 864; oligodendrocyte *Zic5* expression 0.6 ± 6, n = 964, oligodendrocyte *Sox10* expression 456 ± 276, n = 964.

### Motif analysis

We performed de novo motif discovery from the sets of astrocyte and oligodendrocyte enhancers that showed specific and strong expression patterns, excluding those enhancers scored as weak. For astrocytes this list consisted of: AiE0375m, AiE0376h, AiE0376m, AiE0377m, AiE0380h, AiE0381h, AiE0385h, AiE0385m, AiE0380h, AiE0390h, AiE0390m, AiE2120m, AiE2122m, AiE2160m, and ProB12. For oligodendrocytes this list consisted of: AiE0361h, AiE0395h, AiE0395m, AiE0396h, AiE0397m, AiE0398h, AiE0400m, AiE0401h, AiE0403h, AiE0407h, AiE0409h, AiE0409m, AiE0410h, AiE0410m, and AiE0641m. We used MEME-CHIP^67^ to identify recurrent de novo motifs in these sets of sequences, using the parameter-meme-maxw 12, and comparing to a background set of random sequence with the same nucleotide content. This analysis revealed one strong motif in each set of sequences, as measured by its E-value, which is an estimate of the number of motifs expected by chance to have as strong a log likelihood ratio as itself within the given sequences. These de novo motifs where then mapped to known sequences using TomTom^67^ which revealed several possible matches to known motifs at significant p-values, but the strongest motif match (lowest p-value) in each case is shown. In the case of the Zic family transcription factors, for simplicity we averaged together the highly correlated strongest hits in the Zic family (JASPAR accession numbers MA0697.2, MA1628.1, and MA1629.1 covering Zic1, Zic2, and Zic3), since Zic5 itself is not present in databases. In the case of Sox10, the highly correlated Sox family members Sox4 and Sox11 (Uniprobe accession numbers UP00062.1 and UP00030.1) showed slightly stronger motif match p-values than Sox10 (JASPAR accession number MA0442.1), but these were excluded from analysis due to lack of expression in almost all oligodendrocytes as observed by Tasic et al.^97^ and in this study.

### Enhancer-AAV testing in rat

The Allen Institute Institutional Animal Care and Use Committee (IACUC) approved the following in vivo testing experiments in rat under protocol 2010. We procured timed-pregnant female Sprague-Dawley rats from Charles River laboratories. We tattooed and injected ice-anesthetized neonatal pups at P1 with 1.5e11 viral genomes of enhancer-AAV virus, diluted with PBS to a total volume of 10 µL, unilaterally into the forebrain lateral ventricle (ICV delivery) with a 31-gauge, 4 point, 12° bevel 1 inch needle (custom ordered from Hamilton) and 25 µL capacity removable needle syringe (Hamilton, 7636-01). Between injections we washed the needle and syringe with 100% ethanol, and then nuclease-free water. We targeted the ICV space at 2 mm posterior to bregma, 2 mm lateral to the anterior-posterior midline, and at a depth of 2 mm perpendicular to the surface of the skull. We injected into the ventricle slowly over approximately 30 seconds. After injection, we held the needle in place for approximately 10 seconds to prevent viral leakage, then slowly withdrew the needle at the same relative angle as injection and then placed the animal onto a prewarmed heating pad in a clean cage. We sacrificed pups at 18 days post injection, prior to weaning, and transcardially perfused with 1X PBS and then 4% PFA in PBS. We hemisected each brain and cryoprotected in 30% sucrose in deionized water for a minimum of 24 hours before sectioning. We sectioned each brain at 30 µm thickness using a sliding microtome (Leica part number SM2000R) on a leveled mount of Tissue-Tek® O.C.T. Compound, collecting 3 sagittal planes separated by approximately 500 µm. We counterstained sections with 1 µg/mL DAPI and 2.5 µg/mL propidium iodide (Thermo catalog # P1304MP) overnight at 4°C and mounted in Vectashield HardSet Antifade Mounting Medium (Vector Laboratories # H-1400-10) prior to imaging by epifluorescence.

### Macaque enhancer-AAV testing

*Macaca nemestrina* (pig-tailed macaque) animals were housed and injected at the Washington National Primate Center according to NIH guidelines and as approved by the University of Washington Animal Care and Use Committee under UW IACUC protocol #41-6701. These animals received several intraparenchymal injections under general anesthesia at spatially distinct sites located at least ∼1cm apart throughout the brain. During injection, over the course of 10 minutes we expelled a total of approximately 1e11 gc iodixanol gradient-purified PHP.eB-packaged viral vectors in a total volume of 5 µL at 10 depths ranging from 200 to 2000 microns deep in the animals. After injection the animal rested for 10 minutes between injections. These numbers are approximate and timing, volume, and depths may be adjusted according to animal anatomy and surgical considerations. The experiments described here result from four injection sites in three *Macaca nemestrina* animals (two males, one female). We harvested tissue from these animals after necropsy at 38, 47, and 113 days post injection.

After locating the injection sites and cutting out tissue blocks about 1-2cm on each side surrounding the injection sites, we fixed these tissue blocks in 4% PFA for 24 hrs. Then we rinsed the blocks with PBS, cut 350 µm thick slices on the sliding microtome, and postfixed the slices in 4% PFA for 2 hours at room temperature (RT), and then washed three times in PBS for 10 min each. For ISH analysis using HCR we transferred slices to 70% EtOH at 4°C for a minimum of 12 hours, and up to 30 days. To stain we first incubated the slices in 8% SDS in PBS at RT for two hours with agitation, then washed the slices at RT with 5X sodium chloride sodium citrate (SSC) for three hours, exchanging with fresh 5X SSC every hour. Next we performed HCR v3.0 using reagents and a modified protocol from Molecular Technologies and Molecular Instruments^99^. We first incubated slices in pre-warmed 30% probe hybridization buffer (30% formamide, 5X sodium chloride sodium citrate (SSC), 9 mM citric acid pH 6.0, 0.1% Tween 20, 50 µg/mL heparin, 1X Denhardt’s solution, 10% dextran sulfate) at 37°C for 5 min.

Then we exchanged hybridization buffer for hybridization buffer containing probes added at a concentration of 2 nM. Molecular Instruments designed the probes using the following accession numbers: SLC17A7 – XM_011768126.1, GAD1 – XM_011744029.1, FGFR3 – XM_011744842.2, SOX10 – XM_011712410.2. Hybridization proceeded overnight at 37°C, and afterwards we washed the tissue thrice with 5X SSC for 10 minutes each (total 30 minutes), then 30% probe wash buffer (30% formamide, 5X SSC, 9 mM citric acid pH 6.0, 0.1% Tween 20, 50 µg/mL heparin) for one hour at 37°C. Then we exchanged probe wash buffer with 5X SSC, then amplification buffer (5X SSC, 0.1% Tween 20, 10% dextran sulfate) for 5 min at room temperature. Meanwhile we pooled even and odd amplification hairpins for each of the three genes and snap-cooled them by heating to 95°C for 90 seconds then cooling to room temperature for 30 min, and afterwards we added the snap-cooled hairpins to amplification buffer at a final concentration of 60 nM, and finally centrifuged at 18000g for 1 minute. Then we incubated tissue slices in amplification solution containing amplification hairpins for 4 hours at room temperature, followed by staining in DAPI (10µg/mL in 2X SSC) for 1 hour at room temperature, and finally washing twice for 30 min in 2X SSC at room temperature before imaging. We prepared a fresh aliquot of 67% 2,2’-Thiodiethanol (TDE) solution for use as a clearing and immersion fluid by mixing ≥99% TDE (Sigma-Aldrich) with deionized water to create a 67% TDE solution with a refractive index of 1.46. We transferred slices to 67% TDE and allowed them to equilibrate for at least 1 hour at room temperature prior to imaging on a confocal microscope (Olympus FV-3000). For IHC analysis of injected NHP tissue we subsectioned 350 µm thick slices to 30 µm and stained as for mouse tissue.

### Stereotaxic injection and fiber implant surgery

Virus injection and optic fiber implantation surgery was performed in C57BL/6J mice (The Jackson Laboratory, #000664) at around P60. Mice were anesthetized with isoflurane and monitored throughout the surgery using breathing rate and tail pinch. The skin above the skull surface was removed to make room for the fiber implant and headframe. After leveling the skull, a craniotomy was drilled above the injection and fiber coordinates (AP: 1.2 mm, ML: - 1.3 mm, DV: 4.1 mm). First, a glass pipette positioned at the injection coordinates was lowered through the craniotomy and virus injection was performed (100 nl, titer: 4e13 gc/mL). Once the injection was complete, the pipette was slowly raised, and the optic fiber probe was positioned at the same AP and ML coordinates as the injection. The tip of the fiber was then lowered to 100 μm above the injection site and glued in place where the base of the fiber ferrule contacts the skull. A custom headframe was then glued to the skull to allow head-fixed behavior and imaging. After surgery, the mouse was returned to the home cage and allowed to recover for at least two weeks prior to start of water restriction for behavior and imaging.

### Dynamic foraging reinforcement learning task

Water-restricted and head-restrained mice were trained to perform a reinforcement learning task where they freely choose between two lick ports that delivered a water reward with nonstationary probabilities. This is a variation on the task described in Bari et al.^100^. The base reward probability of both lick ports summed to 0.6 where the probabilities of the two lick ports were selected from two sets of ratios (0.53/0.07, 0.51/0.09). Block lengths that corresponded to each ratio lasted for about 30 trials (min trials per block: 40, max trials per block: 60). Each trial began with an auditory “go cue” that signaled the start of a trial. The mouse was free to choose between the left or right lick port immediately after the “go cue”. The trials were separated by a variable inter-trial-interval (range between 1-7 seconds). The data shown in this study was from a two-hour behavior session that consisted of 438 trials (170 rewarded trials).

### Fiber photometry and analysis

Fiber photometry was performed using a commercially available photometry system (Neurophotometrics LLC, FP3002). A 470 nm LED was used to excite the iAChSnFR fluorophore, Venus, and the emitted fluorescence signals were collected using a CMOS camera. The 470 nm excitation was interleaved with a 415 nm LED as an isosbestic control to remove motion artifacts. Bonsai acquisition software was used to record the photometry signals as well as the behavior trigger signals events (go-cue, left and/or right lick choices, reward/no reward) for offline alignment of imaging data to behavioral events. Prior to start of acquisition, an ROI was drawn over the fiber image seen on the camera, and fluorescence intensity within this ROI was averaged for real-time signal visualization and offline analysis. First, the fiber photometry acquisition was started, following which the behavior task was initialized. Photometry signals were analyzed using custom python scripts. First, the fluorescence signal was detrended for photobleaching using a fourth order polynomial function and then corrected for motion using the control signal from the 415 nm excitation using standard photometry analysis techniques^101^. Acetylcholine signal changes were calculated as a change in fluorescence intensity over the mean fluorescence (ΔF/F as a percentage). The photometry signals were then aligned to behavior events using simultaneously acquired TTL readouts of behavior events (go-cue, left and/or right lick choices, reward/no reward) using a NI USB card. These behavior events were then used to calculate trial averaged traces of rewarded and unrewarded signals.

## Data Availability

All AAV viral vector plasmids are freely available for research use at Addgene (addgene.org/). Mouse scRNA-seq generated from this study are available at GEO with the accession number GSE235987 (https://www.ncbi.nlm.nih.gov/geo/). Mouse primary testing data are available at the Allen Genetic Tools Atlas (https://portal.brain-map.org/genetic-tools/genetic-tools-atlas). Serial two photon tomography datasets will be made available through the Brain Imaging Library (https://www.brainimagelibrary.org/). All other data will be made available upon request.

## Acknowledgements

We would like to thank the Washington National Primate Center and staff for animal care, as well as the supporting grant P51OD010425 from the National Institutes of Health to support primate research. The WaNPRC SPF *M. nemestrina* colony is supported by grant U42OD011123 from the NIH Office of Research Infrastructure Programs. We would like to acknowledge Kathryn Gudsnuk for programmatic support. This work was supported by the following grants: RF1MH114126-01 from the National Institute of Mental Health to BPL, JTT, and ESL; UG3MH120095-01, −02, −03 from the National Institute of Mental Health to BPL, JTT, and ESL, UF1MH128339-01 from the National Institute of Mental Health to BT, TEB, TLD, BPL, and JTT, RF1MH121274-01 from the National Institute for Mental Health to BT and U19MH114830 to HZ. We also would like to acknowledge the estate of Paul G. Allen for his vision, encouragement, and support.

---

**Figure 1—figure supplement 1:**
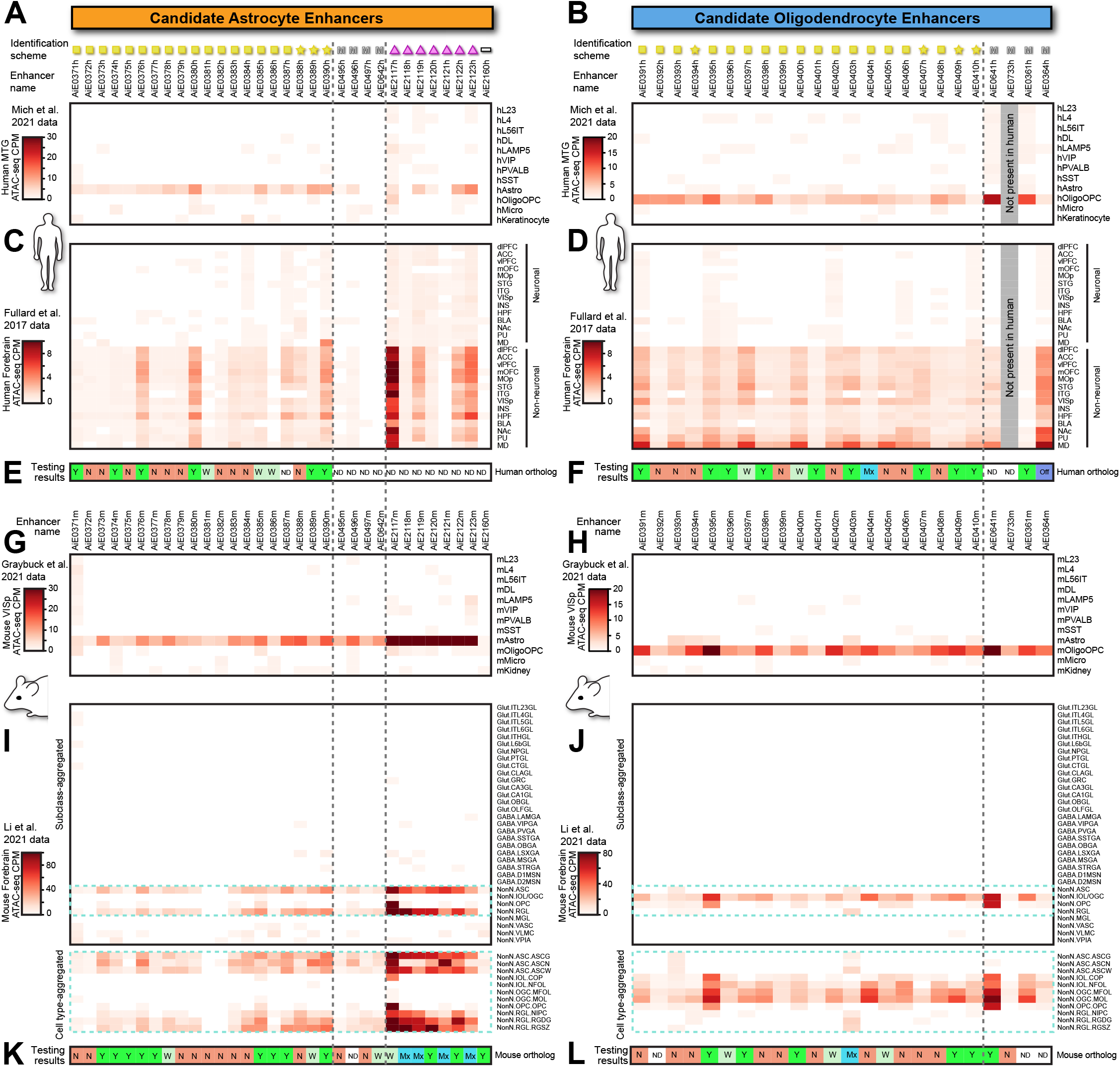
Epigenetic characterization of candidate enhancers in additional chromatin accessibility datasets. (**A-D**) Accessibility profiles of all tested candidate human astrocyte-specific (**A**, **C**) and human oligodendrocyte-specific (**B**, **D**) enhancers. Human enhancer regions are characterized in the datasets of Mich et al.^39^ (**A-B**), who performed snATAC-seq on neurosurgical MTG samples, and of Fullard et al.^102^ (**C-D**), who performed bulk ATAC-seq on neuronal (sorted NeuN^+^) and non-neuronal (sorted NeuN^-^) nuclei from dissections spanning multiple regions of human postmortem forebrain. Overall, many candidate astrocyte- and oligodendrocyte-specific enhancers show accessibility specific to non-neuronal cells across much of the human forebrain. For each genomic region we show their peak nomination scheme matching to **Figure 1B**, enhancer name. (**E-F**) Screening results from testing human candidate enhancers (repeated from **Figure 1D-E**, provided again for visualization). Testing result bar: Y = yes, enhancer-AAV gives strong or moderate on-target expression pattern; N = no, enhancer-AAV fails to express; W = weak on-target expression pattern; Mx = mixed specificities consisting of on-target cells plus unwanted neuronal populations; Off = off-target expression pattern, ND = no data. (**G-J**) Accessibility profiles for all tested candidate mouse astrocyte-specific (**G**, **I**) and oligodendrocyte-specific enhancers (**H**, **J**). Mouse enhancer regions are characterized in the VISp scATAC-seq dataset of Graybuck et al.^40^ (**G-H**) and in the full mouse cerebrum dataset of Li et al.^62^ (**I-J**). (**K-L**) Screening results from testing mouse candidate enhancers (repeated from **Figure 1D-E**, provided again for visualization).

**Figure 2—figure supplement 1:**
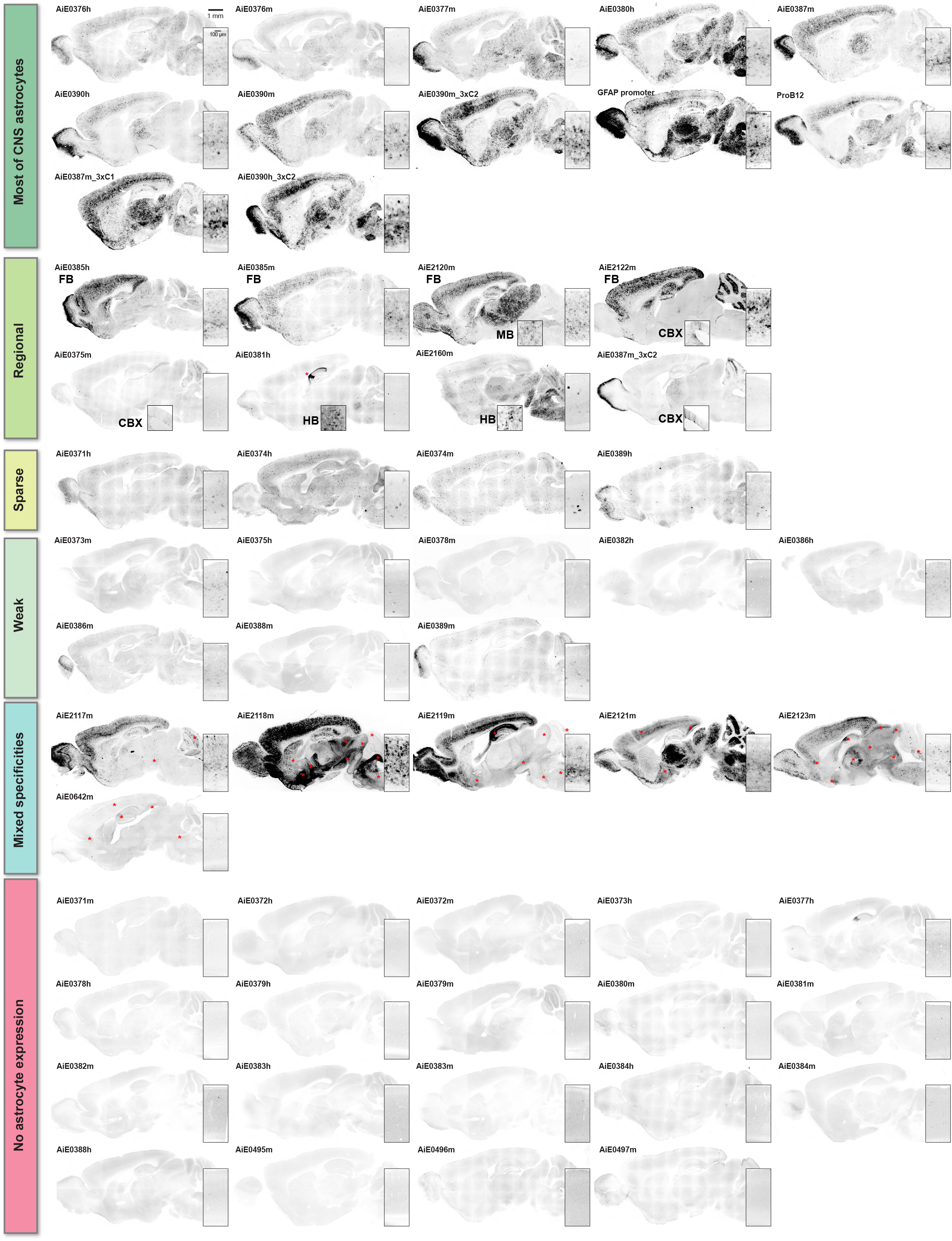
Full screening results of all candidate enhancer-AAVs targeting astrocytes. We injected mice with the indicated enhancer-AAV vectors between P42 and P56, then after 3-4 weeks we harvested brains, sliced them on a sliding microtome with freezing stage at 30 µm thickness, co-stained the sections with DAPI, then mounted them with Vectashield Vybrance. Insets show a full cortical column from VISp (primary visual cortex), and in some cases also the labeling in MB (midbrain) or HB (hindbrain) or CBX (cerebellar cortex) is also shown. Astrocyte-specific enhancer-AAV vectors are broadly grouped by expression pattern into the following categories: “Most of CNS astrocytes”, “Regional” meaning present at medium-to-high levels in one or more broad brain regions but not all, “Sparse” meaning a few astrocytes are strongly labeled throughout the brain, “Weak” meaning many astrocytes throughout the brain are labeled at low level, “Mixed specificities” meaning one or more off-target neuron populations are also labeled in addition to astrocytes, and “No astrocyte expression” meaning failure to detect any clear astrocytes in these whole-brain sagittal images. Red asterisks indicate regions where off-target neuronal labeling was observed. These epifluorescent screening images represent n = 1 to 27 animals tested per enhancer (median 2 animals), and were sometimes taken on multiple different microscopes (see **Table S2** for full summary of screened animals).

**Figure 2—figure supplement 2:**
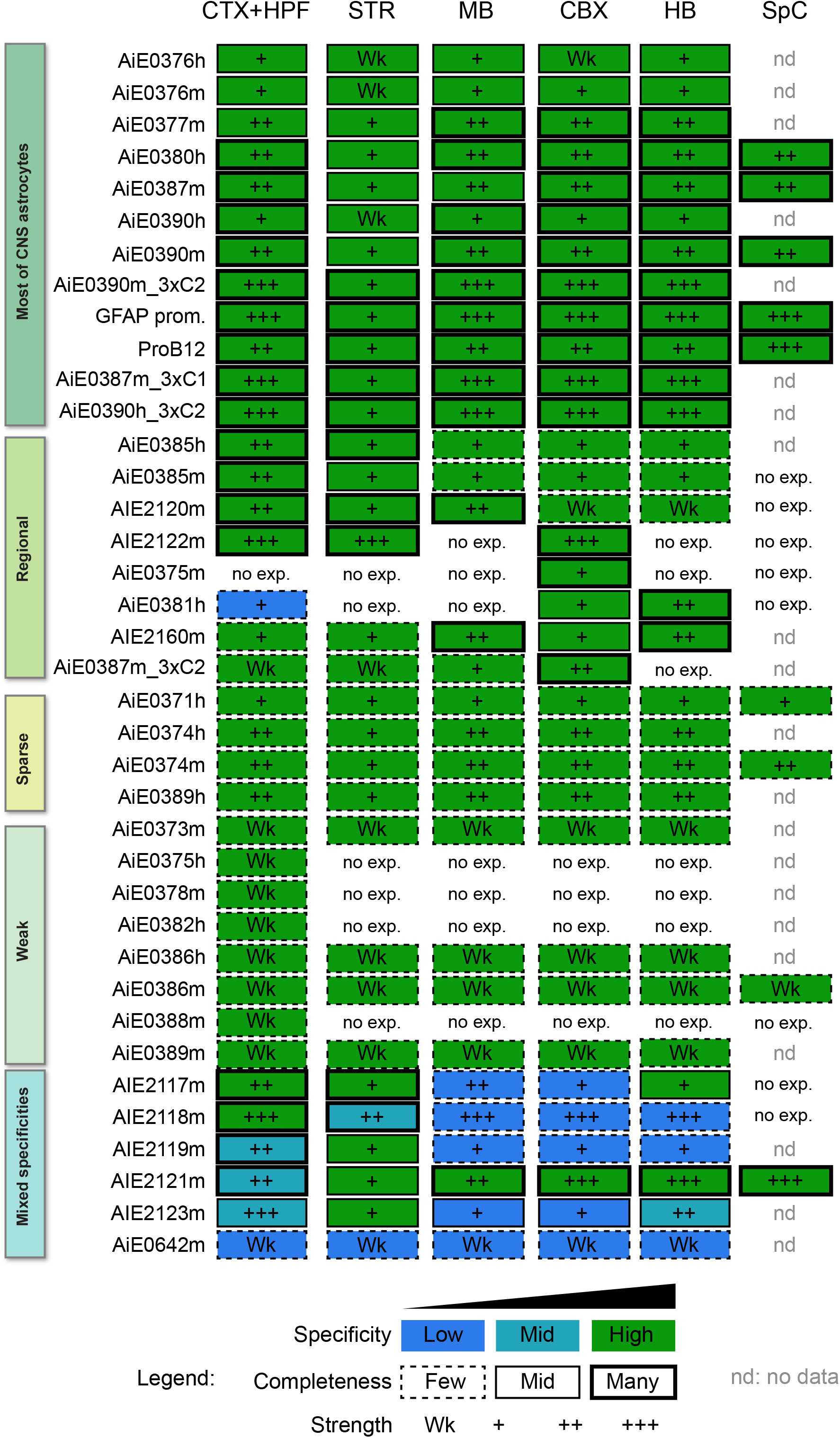
Summary matrix describing screening results of all astrocyte-specific enhancer-AAV vectors. We characterized expression patterns from each of the enhancer-AAV vectors showing astrocyte specificity in regions of the CNS. Specificity ranges from blue (low specificity, many neurons observed) to green (high specificity, >80% specific for astrocytes in the indicated regions). Completeness ranges from dashed line (few astrocytes labeled, estimate <10%) to thick black line (many astrocytes labeled, estimate >50%). Strength of expression ranges from “Wk” (weak expression) to +++ (strong expression). No data is represented by “nd”, and no observable expression in that region is represented by “no exp.”.

**Figure 2—figure supplement 3:**
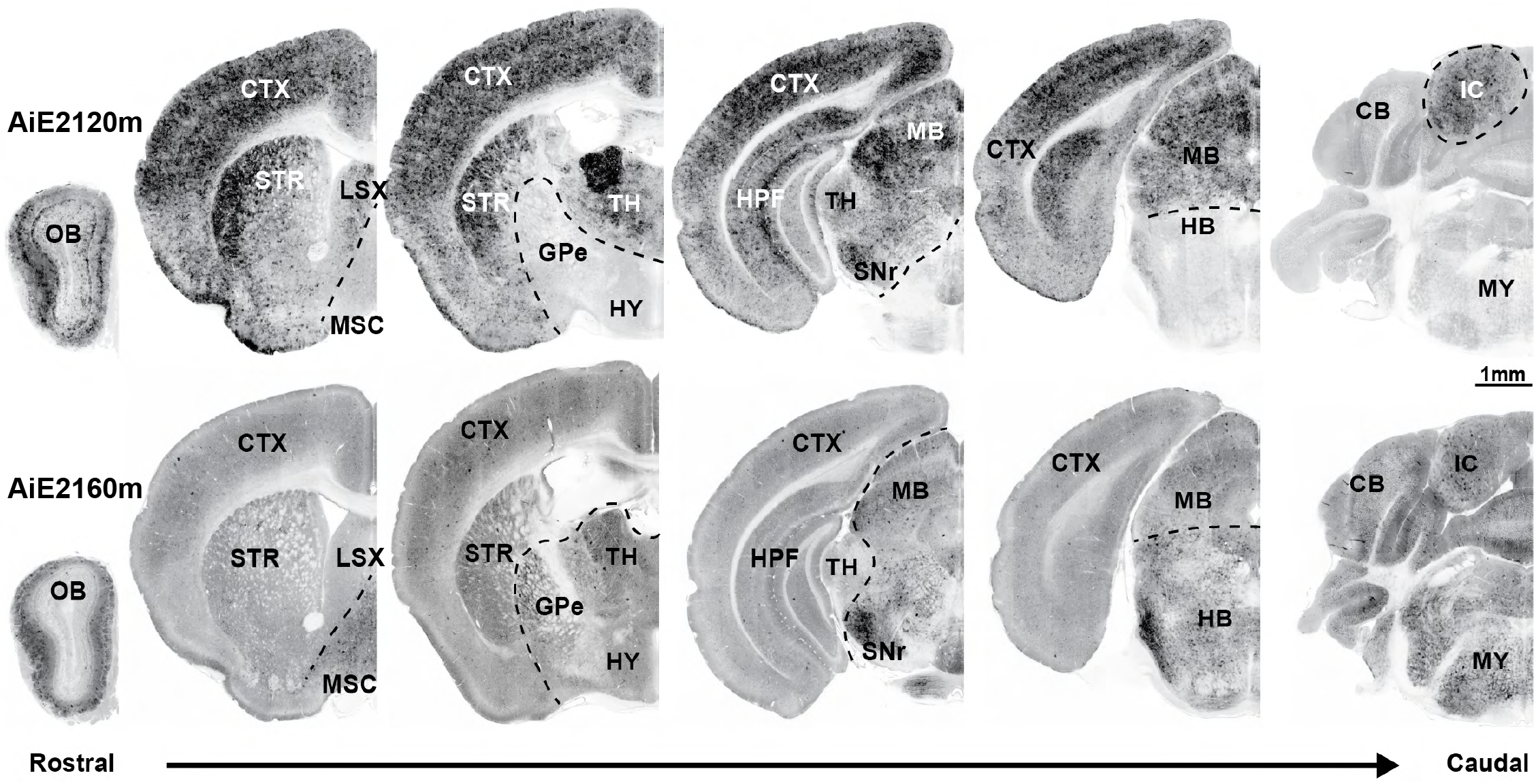
Distinct astrocyte-specific expression domains of AiE2120m and AiE2160m. We injected mice with the indicated astrocyte-specific SYFP2-expressing enhancer-AAVs and performed whole-brain serial two-photon tomography using the TissueCyte platform^96^. These vectors display largely non-overlapping zones of astrocyte expression: AiE2120m is expressed in astrocytes within multiple forebrain structures including CTX, STR, OB, LSX, HPF, and TH, as well as MB, whereas AiE2160m is expressed in MB, CBX, and HB structures as well as complementary forebrain structures including HY, MSC, and GPe, and OB. In the OB AiE2120m is expressed in astrocytes within the granule cell layer, internal plexiform layer, and periglomerular cell layer, whereas AiE2160m is expressed in a complementary pattern of astrocytes within the external plexiform layer. Each image series shows one animal representative of two to three animals tested. Abbreviations: CTX cerebral cortex, STR striatum, OB olfactory bulb, LSX lateral septal complex, HPF hippocampal formation, TH thalamus, MB midbrain, CBX cerebellar cortex, HB hindbrain, HY hypothalamus, MSC medial septal complex, GPe globus pallidus, external layer.

**Figure 3—figure supplement 1:**
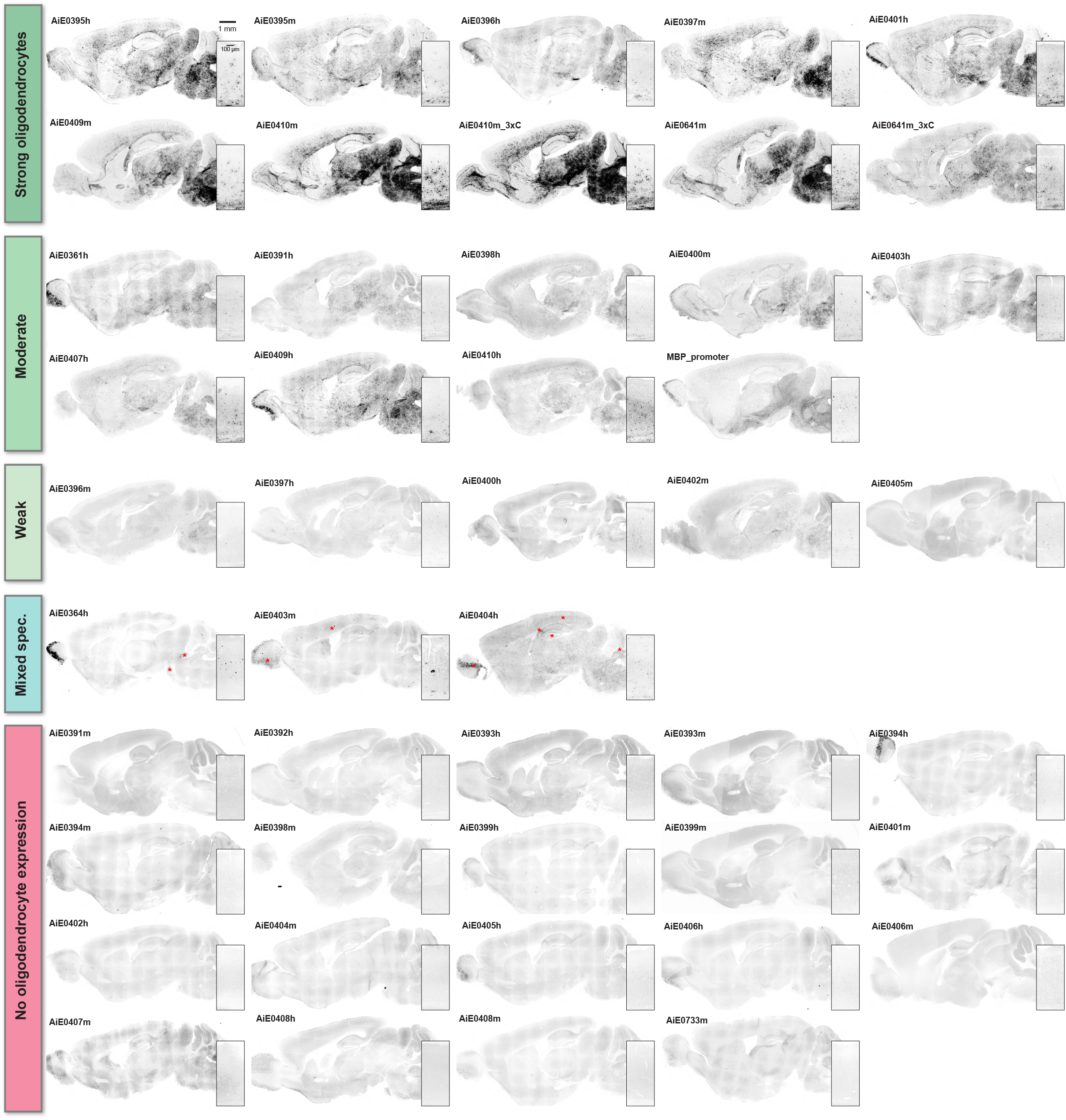
Full screening results of all candidate enhancer-AAVs targeting oligodendrocytes. We injected mice with the indicated enhancer-AAV vectors between P42 and P56, then after 3-4 weeks we harvested brains, sliced them on a sliding microtome with freezing stage at 30 µm thickness, co-stained the sections with DAPI, and mounted them with Vectashield Vybrance. Oligodendrocyte-specific enhancer-AAV vectors are broadly grouped by expression pattern into the following categories: “Strong oligodendrocytes”, “Weak” meaning many oligodendrocytes throughout the brain are labeled at low level, “Mixed specificities” meaning several off-target neuron or astrocyte populations are also present in addition to oligodendrocytes, and “No oligodendrocyte expression” meaning failure to detect any clear oligodendrocytes in these whole-brain sagittal images. Red asterisks indicate regions where off-target neuronal or astrocytic labeling was observed. These epifluorescent screening images represent n = 1 to 20 animals tested per enhancer (median 2 animals), and were sometimes taken on multiple different microscopes (see **Table S2** for full summary of screened animals).

**Figure 5—figure supplement 1:**
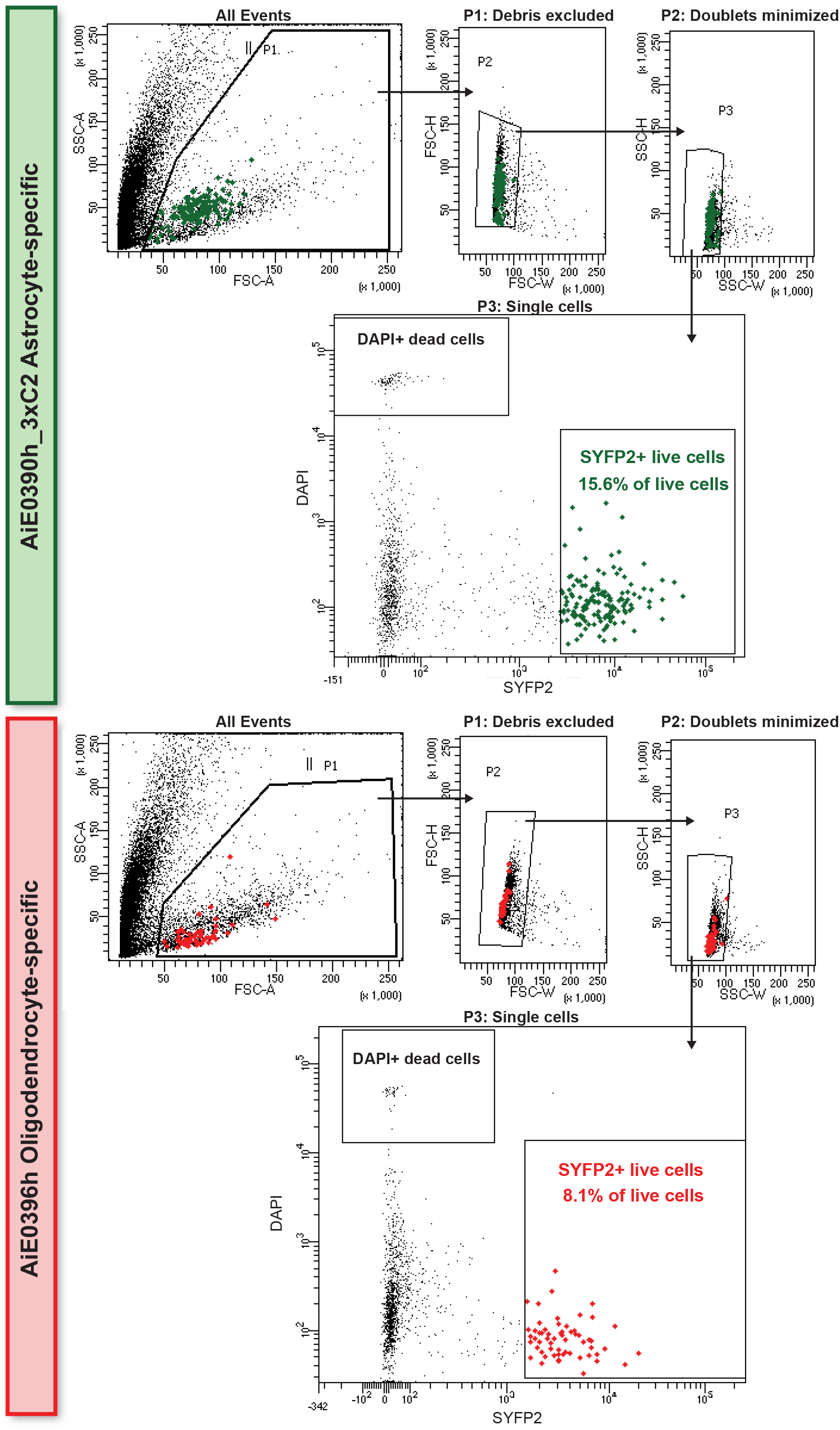
Sorting enhancer-AAV-labeled astrocytes and oligodendrocytes. Example gating strategies for sorting AiE0390h_3xC2-labeled astrocytes and AiE0396h-labeled oligodendrocytes from mouse VISp. Each flow gating strategy represents one animal tested for that vector (n = 26 total animals for astrocytes and n = 21 for oligodendrocytes).

**Figure 11—figure supplement 1:**
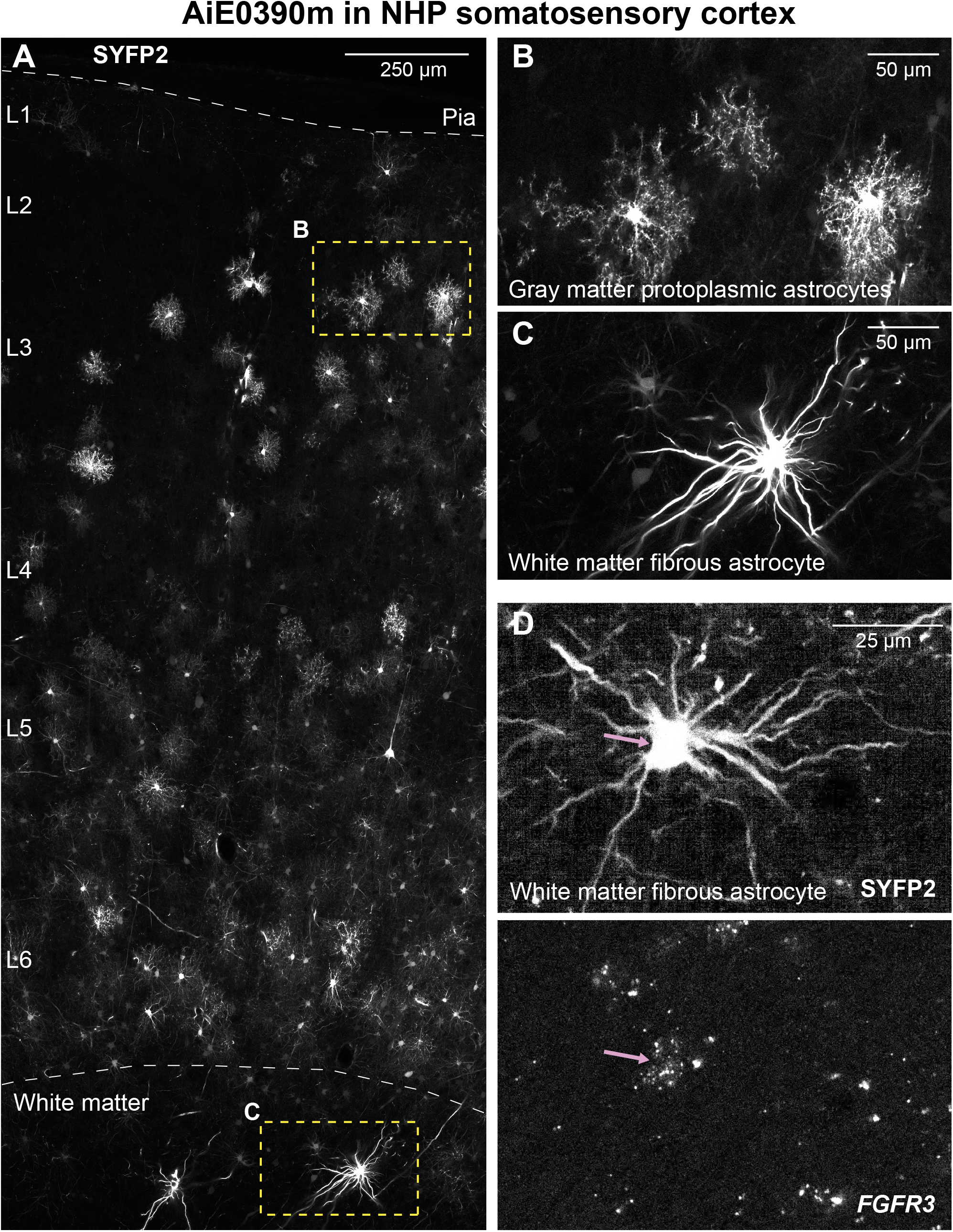
Diverse morphologies of macaque astrocytes labeled by enhancer-AAVs. (**A-C**) Labeling of both gray matter protoplasmic astrocytes and white matter fibrous astrocytes by AiE0390m enhancer-AAV. We show full cortical column of a somatosensory cortex injection site in A, with expanded insets to show protoplasmic astrocytes in gray matter (**B**) and fibrous astrocytes in white matter (**C**). Images represent one animal tested. **(D)** Confirmation of astrocyte identity by mFISH. Fibrous astrocytes in white matter express the astrocyte marker *FGFR3*, similar to gray matter protoplasmic astrocytes (**Figure 8M**).

**Table S1:**
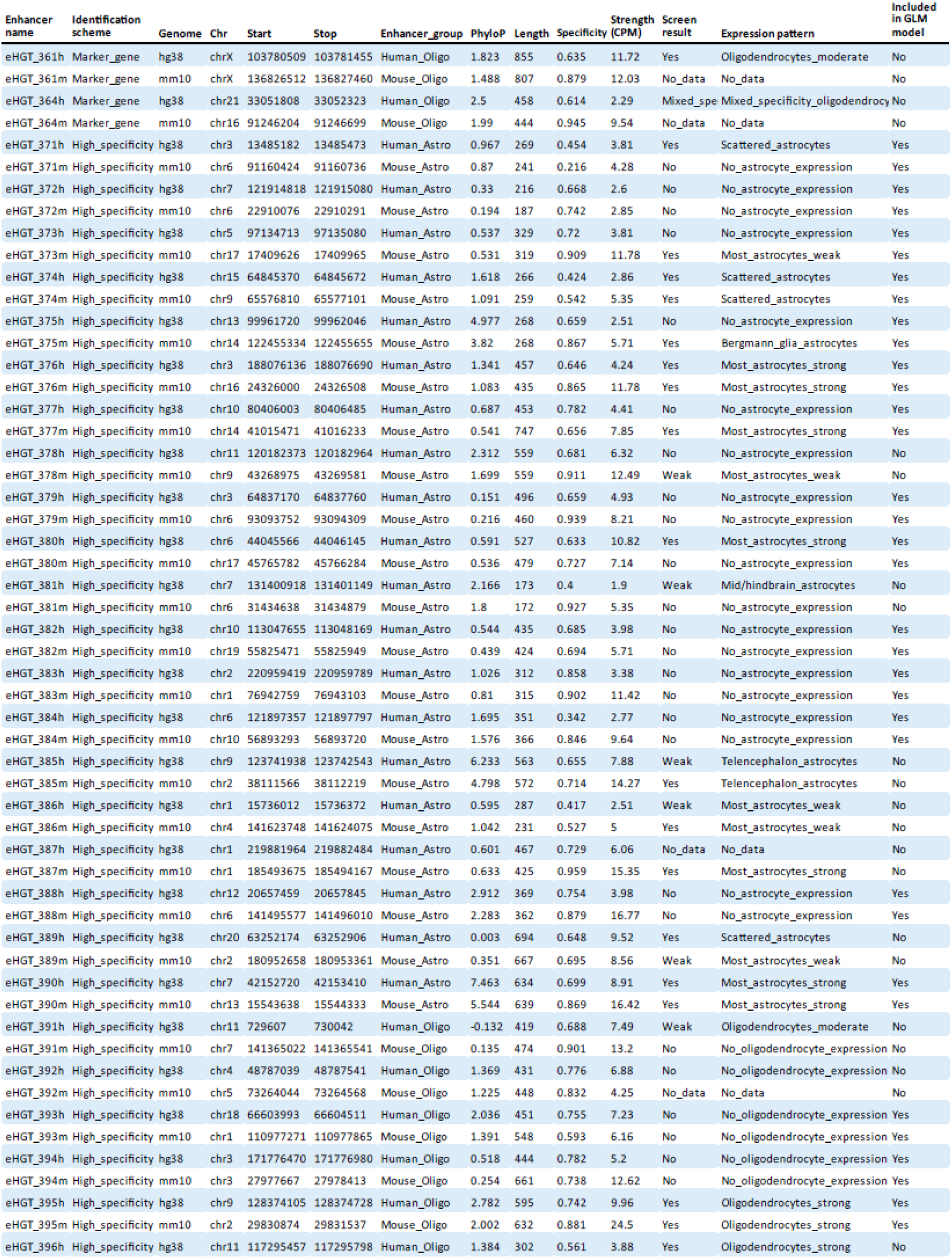
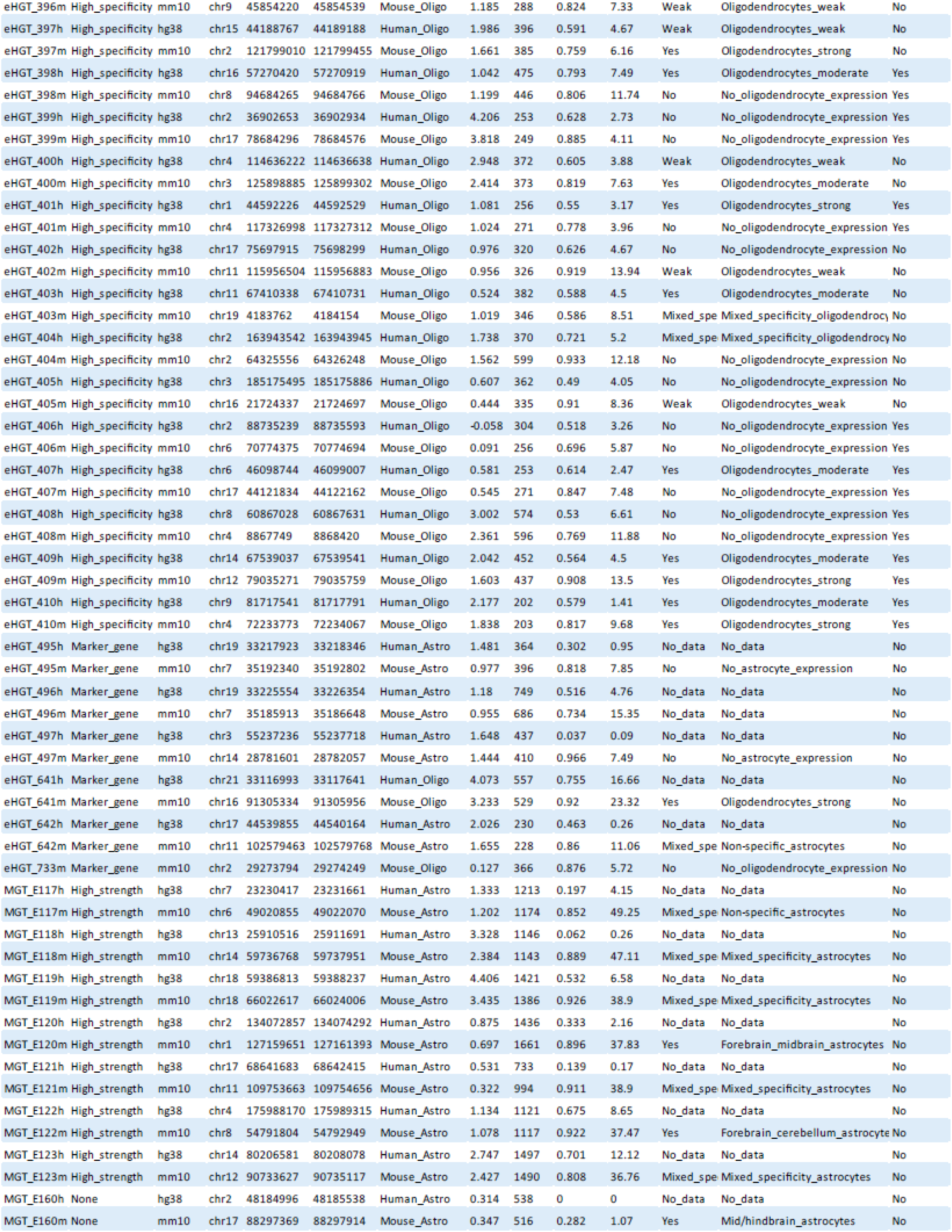
Genomic coordinates, sequence characterization, and mouse screening results of all tested astrocyte and oligodendrocyte enhancers. Calculations of parameters are as described in Methods section.

**Table S2:**
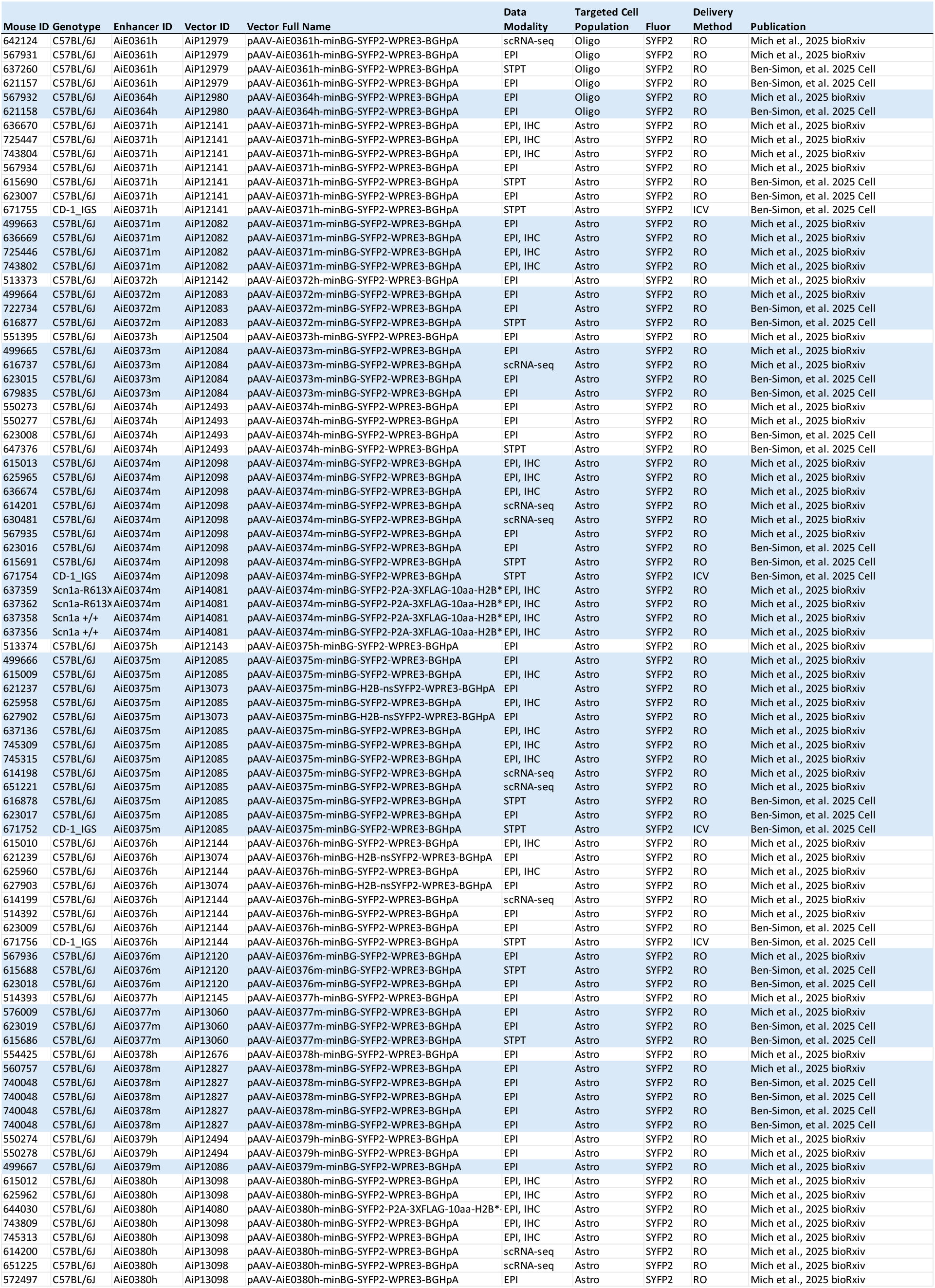
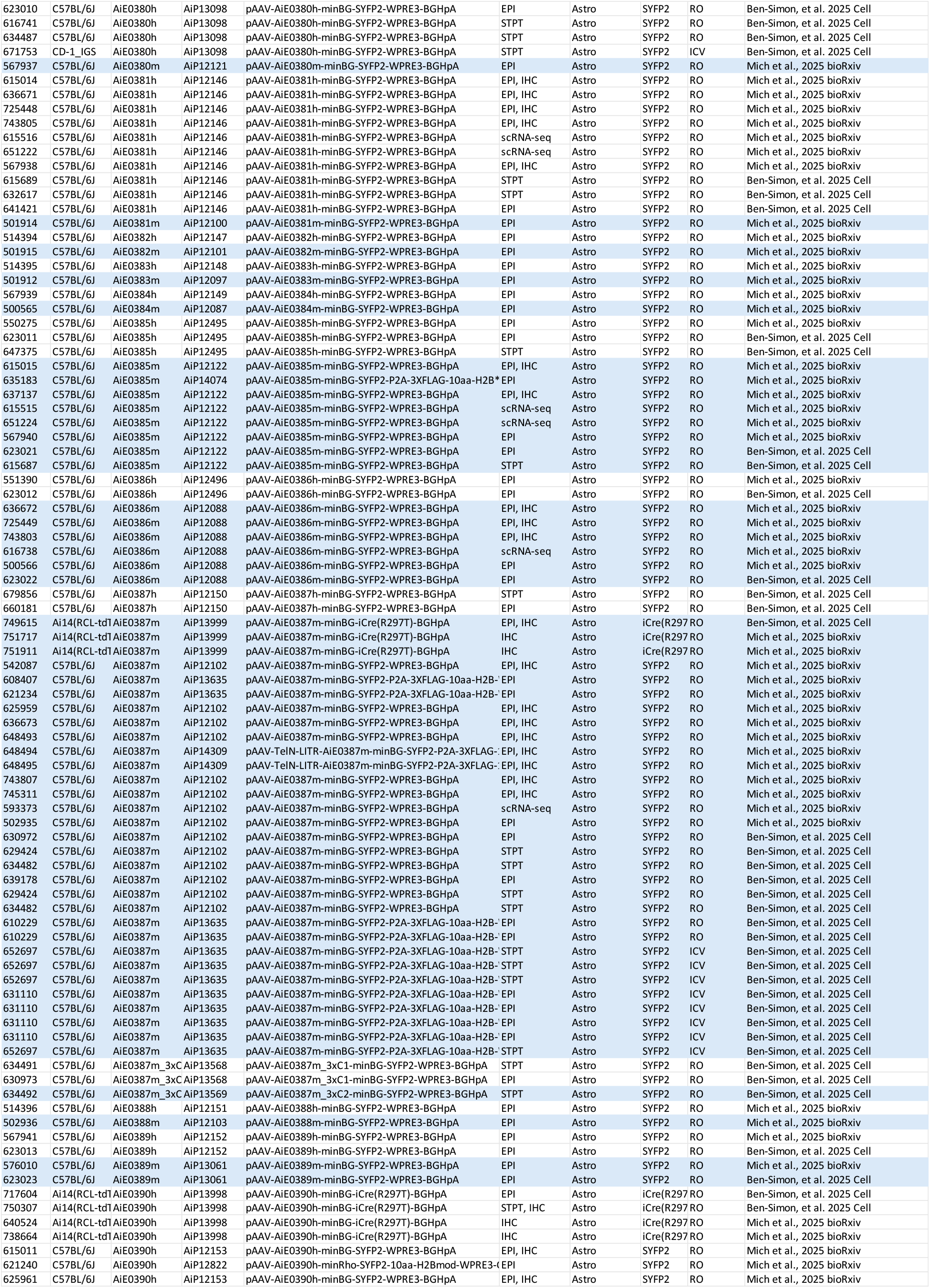
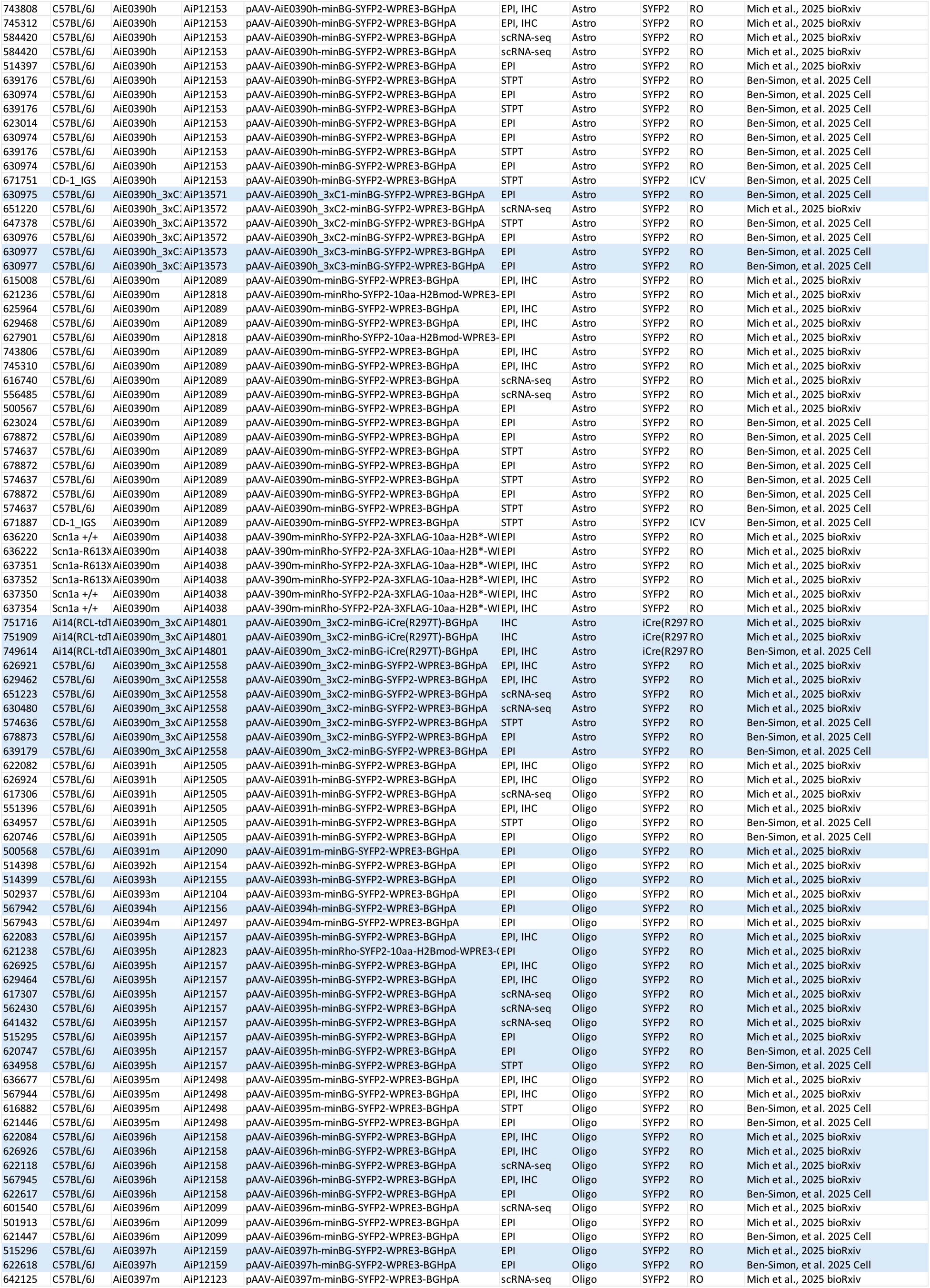
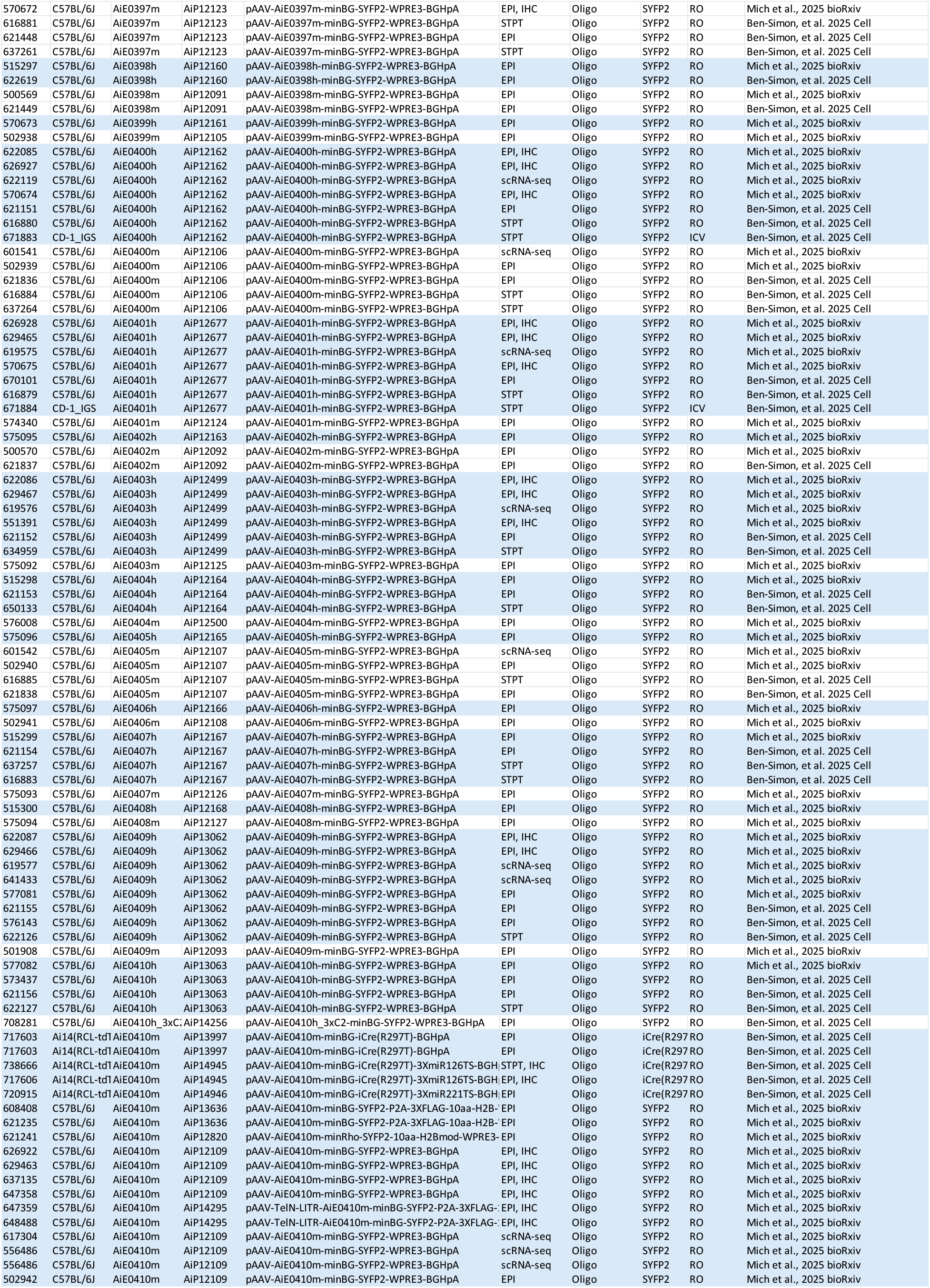
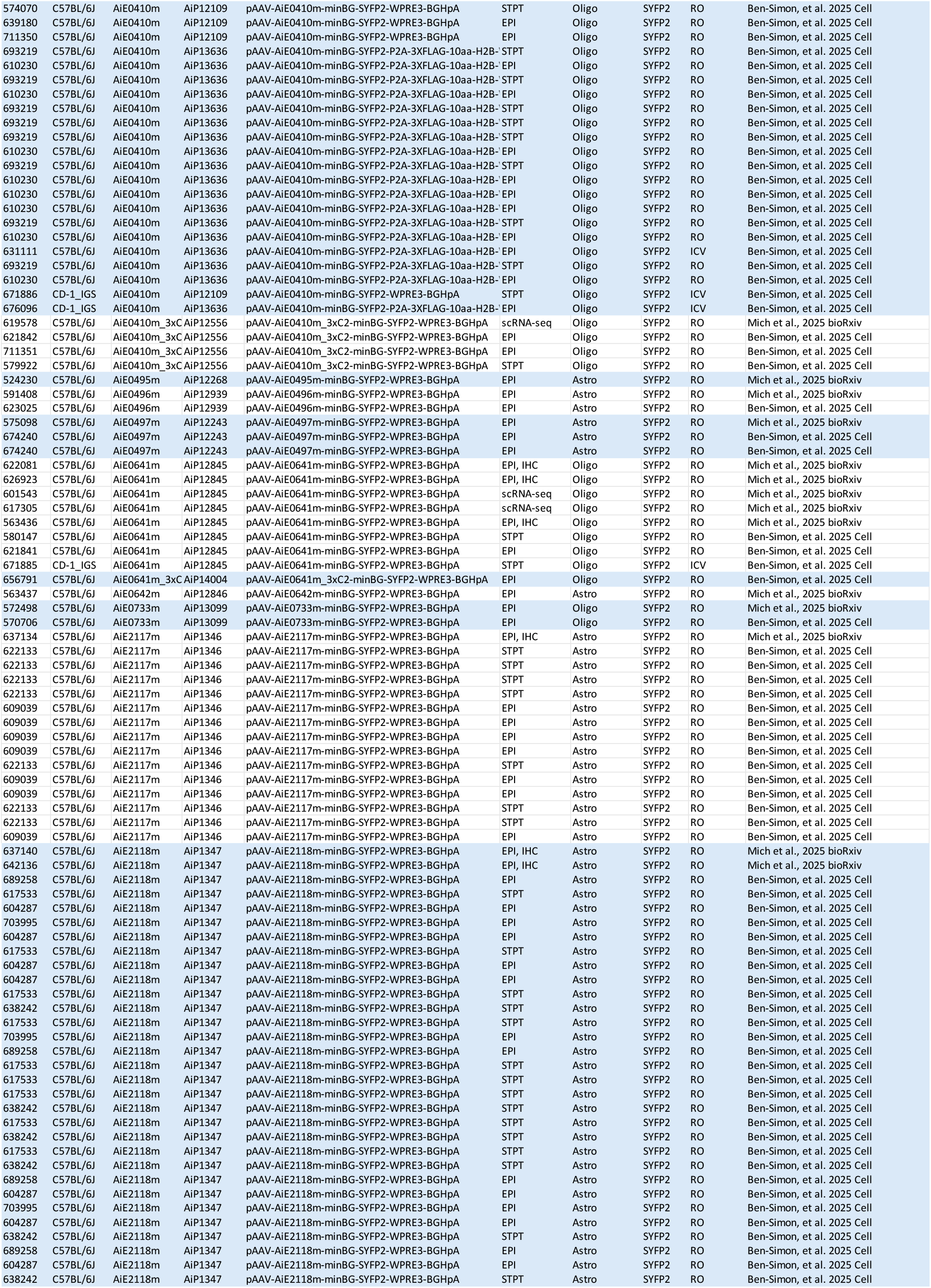
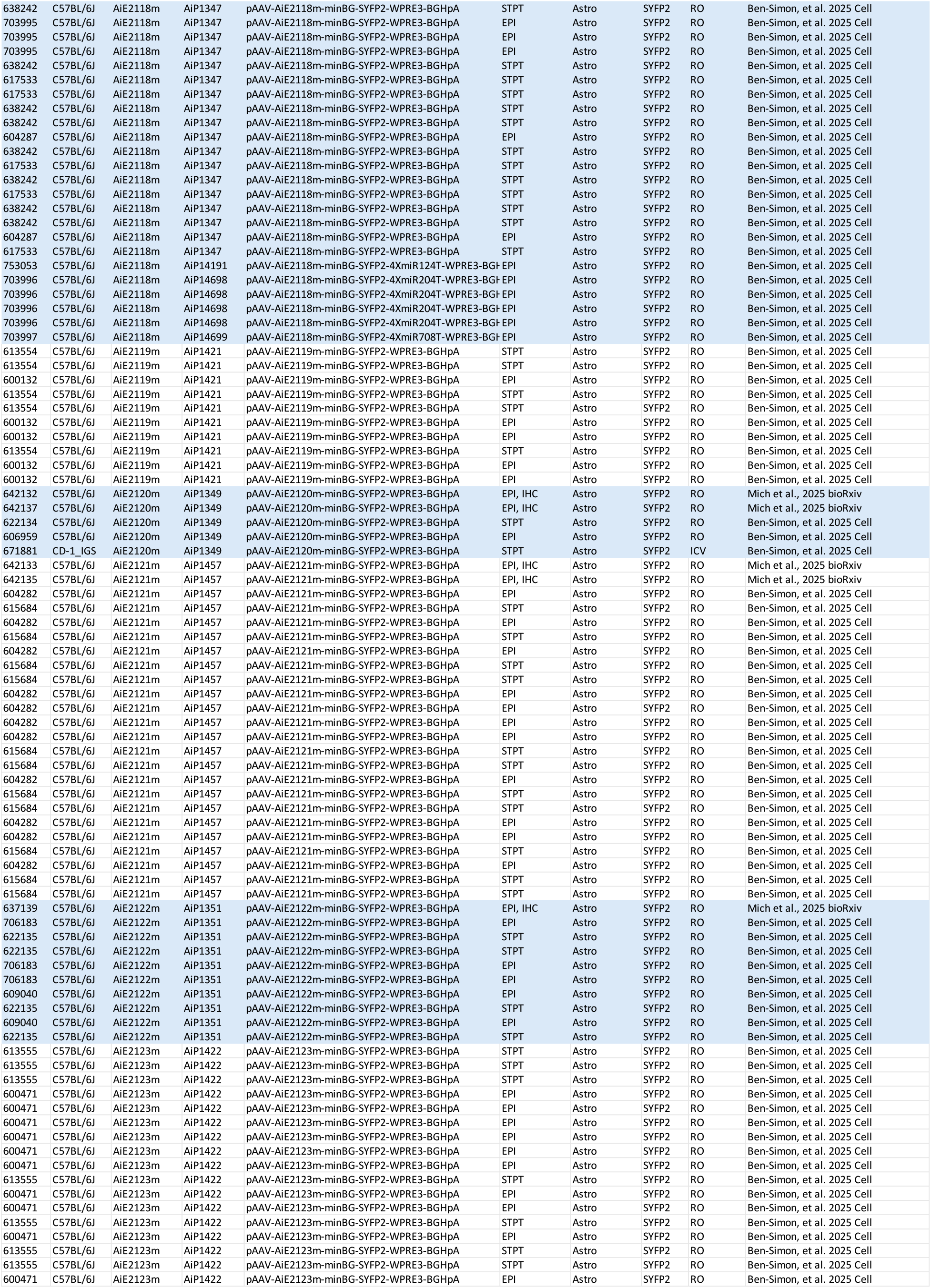
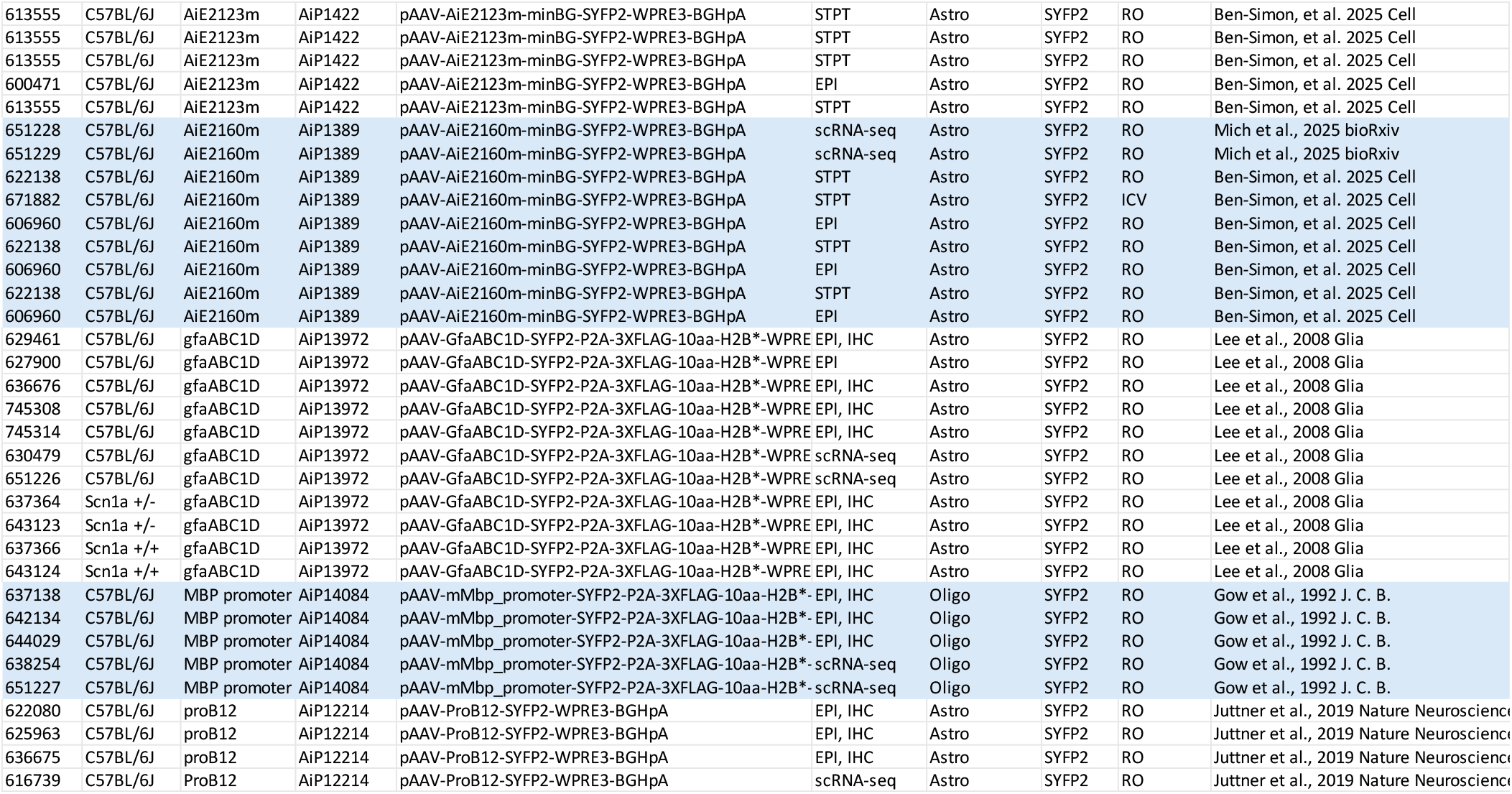
List of mice used for characterizing the activity of astrocyte and oligodendrocyte-specific enhancer-AAV reporter vectors. “Mouse ID” is a unique six-digit identifier for each mouse. “Genotype” describes the genotype. “Enhancer ID” is a unique label for each enhancer or promoter element tested (grouped by colors). “Vector ID” is a unique label for the enhancer-AAV DNA packaged into PHP.eB for testing. “Vector Full Name” is a descriptive name of the elements in each enhancer-AAV. “Data Modality” refers to the data collected from each mouse: EPI epifluorescence, IHC immunohistochemistry, STPT serial two-photon tomography, scRNA-seq single cell RNA-seq (SMARTerV4). “Targeted Cell Population” is Astrocytes or Oligodendrocytes. “Fluor” is SYFP2 or iCre(R297T) used to recombine tdTomato in Ai14 mouse line. “Delivery Method” is retro-orbital (RO) or intracerebroventricular (ICV). “Publication” refers to this work (Mich et al.), or if the mice were also reported on in a previous work (Ben-Simon et al.^41^), or whether the enhancer or promoter was previously described and used here as a positive control (Lee et al.^63^, Gow et al.^65^, Jüttner et al.^47^).

## References

1. Khakh, B. S. & Sofroniew, M. V. Diversity of astrocyte functions and phenotypes in neural circuits. Nat. Neurosci. 18, 942–952 (2015).

2. Zuchero, J. B. & Barres, B. A. Glia in mammalian development and disease. Dev. Camb. Engl. 142, 3805–3809 (2015).

3. Khakh, B. S. & Deneen, B. The Emerging Nature of Astrocyte Diversity. Annu. Rev. Neurosci. 42, 187–207 (2019).

4. Gegenhuber, B. et al. Astrocyte glucocorticoid receptor signalling restricts neuronal plasticity. Nature 655, 1233–1241 (2026).

5. Molofsky, A. V. et al. Astrocyte-encoded positional cues maintain sensorimotor circuit integrity. Nature 509, 189–194 (2014).

6. Kol, A. et al. Astrocytes contribute to remote memory formation by modulating hippocampal–cortical communication during learning. Nat. Neurosci. 23, 1229–1239 (2020).

7. Lioy, D. T. et al. A role for glia in the progression of Rett’s syndrome. Nature 475, 497– 500 (2011).

8. Doron, A. et al. Hippocampal astrocytes encode reward location. Nature 609, 772–778 (2022).

9. Williamson, M. R. et al. Learning-associated astrocyte ensembles regulate memory recall. Nature 637, 478–486 (2025).

10. Volkenhoff, A. et al. Glial Glycolysis Is Essential for Neuronal Survival in Drosophila. Cell Metab. 22, 437–447 (2015).

11. García-Cáceres, C. et al. Astrocytic Insulin Signaling Couples Brain Glucose Uptake with Nutrient Availability. Cell 166, 867–880 (2016).

12. Sabelström, H. et al. Resident Neural Stem Cells Restrict Tissue Damage and Neuronal Loss After Spinal Cord Injury in Mice. Science 342, 637–640 (2013).

13. Hasel, P., Rose, I. V. L., Sadick, J. S., Kim, R. D. & Liddelow, S. A. Neuroinflammatory astrocyte subtypes in the mouse brain. Nat. Neurosci. 24, 1475–1487 (2021).

14. Escartin, C. et al. Reactive astrocyte nomenclature, definitions, and future directions. Nat. Neurosci. 24, 312–325 (2021).

15. Lee, H.-G., Lee, J.-H., Flausino, L. E. & Quintana, F. J. Neuroinflammation: An astrocyte perspective. Sci. Transl. Med. 15, eadi7828 (2023).

16. Zuchero, J. B. et al. CNS Myelin Wrapping Is Driven by Actin Disassembly. Dev. Cell 34, 152–167 (2015).

17. Steadman, P. E. et al. Disruption of Oligodendrogenesis Impairs Memory Consolidation in Adult Mice. Neuron 105, 150–164.e6 (2020).

18. Gibson, E. M. et al. Neuronal Activity Promotes Oligodendrogenesis and Adaptive Myelination in the Mammalian Brain. Science 344, 1252304 (2014).

19. Looser, Z. J. et al. Oligodendrocyte–axon metabolic coupling is mediated by extracellular K+ and maintains axonal health. Nat. Neurosci. 27, 433–448 (2024).

20. Knowles, J. K. et al. Maladaptive myelination promotes generalized epilepsy progression. Nat. Neurosci. 25, 596–606 (2022).

21. Kang, S. H. et al. Degeneration and impaired regeneration of gray matter oligodendrocytes in amyotrophic lateral sclerosis. Nat. Neurosci. 16, 571–579 (2013).

22. Boisvert, M. M., Erikson, G. A., Shokhirev, M. N. & Allen, N. J. The Aging Astrocyte Transcriptome from Multiple Regions of the Mouse Brain. Cell Rep. 22, 269–285 (2018).

23. Endo, F. et al. Molecular basis of astrocyte diversity and morphology across the CNS in health and disease. Science 378, eadc9020 (2022).

24. Yao, Z. et al. A high-resolution transcriptomic and spatial atlas of cell types in the whole mouse brain. 2023.03.06.531121 Preprint at 10.1101/2023.03.06.531121 (2023).

25. Hodge, R. D. et al. Conserved cell types with divergent features in human versus mouse cortex. Nature 573, 61–68 (2019).

26. Chen, Z.-P. et al. Lipid-accumulated reactive astrocytes promote disease progression in epilepsy. Nat. Neurosci. 26, 542–554 (2023).

27. Wang, C. et al. Selective removal of astrocytic APOE4 strongly protects against tau-mediated neurodegeneration and decreases synaptic phagocytosis by microglia. Neuron 10.1016/j.neuron.2021.03.024 (2021) doi:10.1016/j.neuron.2021.03.024.

28. Monje, M. et al. Hedgehog-responsive candidate cell of origin for diffuse intrinsic pontine glioma. Proc. Natl. Acad. Sci. U. S. A. 108, 4453–4458 (2011).

29. Alcantara Llaguno, S., et al. Malignant Astrocytomas Originate from Neural Stem/Progenitor Cells in a Somatic Tumor Suppressor Mouse Model. Cancer Cell 15, 45–56 (2009).

30. Deverman, B. E. et al. Cre-dependent selection yields AAV variants for widespread gene transfer to the adult brain. Nat. Biotechnol. 34, 204–209 (2016).

31. Chan, K. Y. et al. Engineered AAVs for efficient noninvasive gene delivery to the central and peripheral nervous systems. Nat. Neurosci. 20, 1172–1179 (2017).

32. Nonnenmacher, M. et al. Rapid evolution of blood-brain-barrier-penetrating AAV capsids by RNA-driven biopanning. Mol. Ther. - Methods Clin. Dev. 20, 366–378 (2021).

33. De, A., El-Shamayleh, Y. & Horwitz, G. D. Fast and reversible neural inactivation in macaque cortex by optogenetic stimulation of GABAergic neurons. eLife 9, e52658 (2020).

34. El-Shamayleh, Y., Kojima, Y., Soetedjo, R. & Horwitz, G. D. Selective Optogenetic Control of Purkinje Cells in Monkey Cerebellum. Neuron 95, 51–62.e4 (2017).

35. Mendell, J. R. et al. Single-Dose Gene-Replacement Therapy for Spinal Muscular Atrophy. 10.1056/NEJMoa1706198 https://www.nejm.org/doi/10.1056/NEJMoa1706198?url_ver=Z39.88-2003&rfr_id=ori%3Arid%3Acrossref.org&rfr_dat=cr_pub%3Dwww.ncbi.nlm.nih.gov (2017) doi:10.1056/NEJMoa1706198.

36. Dimidschstein, J. et al. A viral strategy for targeting and manipulating interneurons across vertebrate species. Nat. Neurosci. 19, 1743–1749 (2016).

37. Vormstein-Schneider, D. et al. Viral manipulation of functionally distinct interneurons in mice, non-human primates and humans. Nat. Neurosci. 1–8 (2020) doi:10.1038/s41593-020-0692-9.

38. Hrvatin, S. et al. A scalable platform for the development of cell-type-specific viral drivers. eLife 8, e48089 (2019).

39. Mich, J. K. et al. Functional enhancer elements drive subclass-selective expression from mouse to primate neocortex. Cell Rep. 34, 108754 (2021).

40. Graybuck, L. T. et al. Enhancer viruses for combinatorial cell-subclass-specific labeling. Neuron 109, 1449–1464.e13 (2021).

41. Ben-Simon, Y. et al. A suite of enhancer AAVs and transgenic mouse lines for genetic access to cortical cell types. Cell 10.1016/j.cell.2025.05.002 (2025) doi:10.1016/j.cell.2025.05.002.

42. Johansen, N. J. et al. Evaluating methods for the prediction of cell-type-specific enhancers in the mammalian cortex. Cell Genomics 100879 (2025) doi:10.1016/j.xgen.2025.100879.

43. Kussick, E. et al. Enhancer AAVs for targeting spinal motor neurons and descending motor pathways in rodents and macaque. Cell Rep. 115730 (2025) doi:10.1016/j.celrep.2025.115730.

44. Hunker, A. C. et al. Enhancer AAV toolbox for accessing and perturbing striatal cell types and circuits. Neuron 113, 1507–1524.e17 (2025).

45. Furlanis, E. et al. An enhancer-AAV toolbox to target and manipulate distinct interneuron subtypes. Neuron 113, 1525–1547.e15 (2025).

46. Li, L. et al. Identification and application of cell-type-specific enhancers for the macaque brain. Cell 0, (2025).

47. Jüttner, J. et al. Targeting neuronal and glial cell types with synthetic promoter AAVs in mice, non-human primates and humans. Nat. Neurosci. 22, 1345–1356 (2019).

48. Lin, C.-H. et al. Identification of cis-regulatory modules for adeno-associated virus-based cell-type-specific targeting in the retina and brain. J. Biol. Chem. 298, (2022).

49. Gleichman, A. J., Kawaguchi, R., Sofroniew, M. V. & Carmichael, S. T. A toolbox of astrocyte-specific, serotype-independent adeno-associated viral vectors using microRNA targeting sequences. Nat. Commun. 14, 7426 (2023).

50. Wang, L.-L. et al. Revisiting astrocyte to neuron conversion with lineage tracing in vivo. Cell 184, 5465–5481.e16 (2021).

51. Xie, Y., Zhou, J. & Chen, B. Critical examination of Ptbp1-mediated glia-to-neuron conversion in the mouse retina. Cell Rep. 39, 110960 (2022).

52. Le, N., Appel, H., Pannullo, N., Hoang, T. & Blackshaw, S. Ectopic insert-dependent neuronal expression of GFAP promoter-driven AAV constructs in adult mouse retina. Front. Cell Dev. Biol. 10, (2022).

53. Borden, P. M. et al. A fast genetically encoded fluorescent sensor for faithful in vivo acetylcholine detection in mice, fish, worms and flies. 2020.02.07.939504 Preprint at 10.1101/2020.02.07.939504 (2020).

54. Fröhlich, D. et al. Dual-function AAV gene therapy reverses late-stage Canavan disease pathology in mice. Front. Mol. Neurosci. 15, 1061257 (2022).

55. Hillen, A. E. J. et al. In vivo targeting of a variant causing vanishing white matter using CRISPR/Cas9. Mol. Ther. - Methods Clin. Dev. 25, 17–25 (2022).

56. Hagemann, T. L. et al. Antisense therapy in a rat model of Alexander disease reverses GFAP pathology, white matter deficits, and motor impairment. Sci. Transl. Med. 13, eabg4711 (2021).

57. Lister, R. et al. Global Epigenomic Reconfiguration During Mammalian Brain Development. Science 341, 1237905 (2013).

58. Luo, C. et al. Single-cell methylomes identify neuronal subtypes and regulatory elements in mammalian cortex. Science 357, 600–604 (2017).

59. Lee, D.-S. et al. Simultaneous profiling of 3D genome structure and DNA methylation in single human cells. Nat. Methods 16, 999–1006 (2019).

60. Liu, H. et al. DNA methylation atlas of the mouse brain at single-cell resolution. Nature 598, 120–128 (2021).

61. Fullard, J. F. et al. Open chromatin profiling of human postmortem brain infers functional roles for non-coding schizophrenia loci. Hum. Mol. Genet. 10.1093/hmg/ddy229 (2019) doi:10.1093/hmg/ddy229.

62. Li, Y. E. et al. An atlas of gene regulatory elements in adult mouse cerebrum. Nature 598, 129–136 (2021).

63. Lee, Y., Messing, A., Su, M. & Brenner, M. GFAP promoter elements required for region-specific and astrocyte-specific expression. Glia 56, 481–493 (2008).

64. Sun, W. et al. SOX9 Is an Astrocyte-Specific Nuclear Marker in the Adult Brain Outside the Neurogenic Regions. J. Neurosci. 37, 4493–4507 (2017).

65. Gow, A., Friedrich, V. L., Jr & Lazzarini, R. A. Myelin basic protein gene contains separate enhancers for oligodendrocyte and Schwann cell expression. J. Cell Biol. 119, 605– 616 (1992).

66. Bin, J. M., Harris, S. N. & Kennedy, T. E. The oligodendrocyte-specific antibody ‘CC1’ binds Quaking 7. J. Neurochem. 139, 181–186 (2016).

67. Bailey, T. L. et al. MEME Suite: tools for motif discovery and searching. Nucleic Acids Res. 37, W202–W208 (2009).

68. Hume, M. A., Barrera, L. A., Gisselbrecht, S. S. & Bulyk, M. L. UniPROBE, update 2015: new tools and content for the online database of protein-binding microarray data on protein–DNA interactions. Nucleic Acids Res. 43, D117–D122 (2015).

69. Kulakovskiy, I. V. et al. HOCOMOCO: towards a complete collection of transcription factor binding models for human and mouse via large-scale ChIP-Seq analysis. Nucleic Acids Res. 46, D252–D259 (2018).

70. Castro-Mondragon, J. A. et al. JASPAR 2022: the 9th release of the open-access database of transcription factor binding profiles. Nucleic Acids Res. 50, D165 (2022).

71. Furushima, K., Murata, T., Kiyonari, H. & Aizawa, S. Characterization of Opr deficiency in mouse brain: Subtle defects in dorsomedial telencephalon and medioventral forebrain. Dev. Dyn. 232, 1056–1061 (2005).

72. Inoue, T., Ota, M., Ogawa, M., Mikoshiba, K. & Aruga, J. Zic1 and Zic3 Regulate Medial Forebrain Development through Expansion of Neuronal Progenitors. J. Neurosci. 27, 5461– 5473 (2007).

73. Hornig, J. et al. The Transcription Factors Sox10 and Myrf Define an Essential Regulatory Network Module in Differentiating Oligodendrocytes. PLOS Genet. 9, e1003907 (2013).

74. Turnescu, T. et al. Sox8 and Sox10 jointly maintain myelin gene expression in oligodendrocytes. Glia 66, 279–294 (2018).

75. Hinderer, C. et al. Severe Toxicity in Nonhuman Primates and Piglets Following High-Dose Intravenous Administration of an Adeno-Associated Virus Vector Expressing Human SMN. Hum. Gene Ther. 29, 285–298 (2018).

76. Zhang, K., et al. A Cell Atlas of Chromatin Accessibility across 25 Adult Human Tissues. 2021.02.17.431699 https://www.biorxiv.org/content/10.1101/2021.02.17.431699v1 (2021) doi:10.1101/2021.02.17.431699.

77. Goertsen, D. et al. AAV capsid variants with brain-wide transgene expression and decreased liver targeting after intravenous delivery in mouse and marmoset. Nat. Neurosci. 25, 106–115 (2022).

78. Liddelow, S. A. et al. Neurotoxic reactive astrocytes are induced by activated microglia. Nature 541, 481–487 (2017).

79. Martín-Suárez, S., Abiega, O., Ricobaraza, A., Hernandez-Alcoceba, R. & Encinas, J. M. Alterations of the Hippocampal Neurogenic Niche in a Mouse Model of Dravet Syndrome. Front. Cell Dev. Biol. 8, (2020).

80. Madisen, L. et al. A robust and high-throughput Cre reporting and characterization system for the whole mouse brain. Nat. Neurosci. 13, 133–140 (2010).

81. Hartung, M. & Kisters-Woike, B. Cre Mutants with Altered DNA Binding Properties*. J. Biol. Chem. 273, 22884–22891 (1998).

82. Wang, S. et al. The Endothelial-Specific MicroRNA miR-126 Governs Vascular Integrity and Angiogenesis. Dev. Cell 15, 261–271 (2008).

83. Sayeg, M. K. et al. Rationally Designed MicroRNA-Based Genetic Classifiers Target Specific Neurons in the Brain. ACS Synth. Biol. 4, 788–795 (2015).

84. Cho, F. S. et al. Enhancing GAT-3 in thalamic astrocytes promotes resilience to brain injury in rodents. Sci. Transl. Med. 14, eabj4310 (2022).

85. Hasel, P., Aisenberg, W. H., Bennett, F. C. & Liddelow, S. A. Molecular and metabolic heterogeneity of astrocytes and microglia. Cell Metab. 35, 555–570 (2023).

86. Aida, T. et al. Astroglial glutamate transporter deficiency increases synaptic excitability and leads to pathological repetitive behaviors in mice. Neuropsychopharmacol. Off. Publ. Am. Coll. Neuropsychopharmacol. 40, 1569–1579 (2015).

87. Li, Z. et al. Cell-Type-Specific Afferent Innervation of the Nucleus Accumbens Core and Shell. Front. Neuroanat. 12, (2018).

88. Beier, K. T. et al. Circuit Architecture of VTA Dopamine Neurons Revealed by Systematic Input-Output Mapping. Cell 162, 622–634 (2015).

89. Ren, J. et al. Anatomically Defined and Functionally Distinct Dorsal Raphe Serotonin Sub-systems. Cell 175, 472–487.e20 (2018).

90. Ren, J. et al. Single-cell transcriptomes and whole-brain projections of serotonin neurons in the mouse dorsal and median raphe nuclei. eLife 8, e49424 (2019).

91. Gabitto, M. I. et al. Integrated multimodal cell atlas of Alzheimer’s disease. 2023.05.08.539485 Preprint at 10.1101/2023.05.08.539485 (2023).

92. Kamath, T. et al. Single-cell genomic profiling of human dopamine neurons identifies a population that selectively degenerates in Parkinson’s disease. Nat. Neurosci. 25, 588–595 (2022).

93. Zamboni, M. et al. The regulatory code of injury-responsive enhancers enables precision cell-state targeting in the CNS. Nat. Neurosci. 1–13 (2025) doi:10.1038/s41593-025-02131-w.

94. Pagès, Hervé. BSgenome: Software infrastructure for efficient representation of full genomes and their SNPs. R package version 1.62.0. (2021).

95. Kim, J.-Y., Grunke, S. D., Levites, Y., Golde, T. E. & Jankowsky, J. L. Intracerebroventricular Viral Injection of the Neonatal Mouse Brain for Persistent and Widespread Neuronal Transduction. JoVE J. Vis. Exp. e51863 (2014) doi:10.3791/51863.

96. Ragan, T. et al. Serial two-photon tomography for automated ex vivo mouse brain imaging. Nat. Methods 9, 255–258 (2012).

97. Tasic, B. et al. Shared and distinct transcriptomic cell types across neocortical areas. Nature 563, 72 (2018).

98. Traag, V. A., Waltman, L. & van Eck, N. J. From Louvain to Leiden: guaranteeing well-connected communities. Sci. Rep. 9, 5233 (2019).

99. Choi, J.-H. et al. Optimization of AAV expression cassettes to improve packaging capacity and transgene expression in neurons. Mol. Brain 7, 17 (2014).

100. Bari, B. A. et al. Stable Representations of Decision Variables for Flexible Behavior. Neuron 103, 922–933.e7 (2019).

101. Akam, T. & Walton, M. E. pyPhotometry: Open source Python based hardware and software for fiber photometry data acquisition. Sci. Rep. 9, 3521 (2019).

102. Fullard, J. F. et al. An atlas of chromatin accessibility in the adult human brain. Genome Res. 28, 1243–1252 (2018).

